# Dicer deficiency in microglia leads to accelerated demyelination and failed remyelination

**DOI:** 10.1101/2023.10.17.562812

**Authors:** Ajai Tripathi, Nagendra Rai, Aaron Perles, Claire Jones, Ranjan Dutta

**Author notes:** Contact Information, **Ranjan Dutta, Ph.D.**, Department of Neurosciences, Lerner Research Institute, Cleveland Clinic 9500 Euclid Avenue/ NC30, Cleveland, OH 44195.

## Abstract

Microglia are the resident immune cells of the central nervous system (CNS) and are important regulators of normal brain functions. In CNS demyelinating diseases like multiple sclerosis (MS), the functions of these cells are of particular interest. Here we probed the impact of microRNA (miRNA)-mediated post-transcriptional gene regulation using a mouse model lacking microglia/macrophage-specific *Dicer* expression during demyelination and remyelination. Conditional *Dicer* ablation and loss of miRNAs in adult microglia led to extensive demyelination and impaired myelin processing. Interestingly, demyelination was accompanied by increased apoptosis of mature oligodendrocytes (OLs) and arresting OL progenitor cells (OPCs) in the precursor stage. At the transcriptional level, *Dicer*-deficient microglia led to downregulation of microglial homeostatic genes, increased cell proliferation, and a shift towards a disease-associated phenotype. Loss of remyelination efficiency in these mice was accompanied by stalling of OPCs in the precursor stage. Collectively, these results highlight a new role of microglial miRNAs in promoting a pro-regenerative phenotype in addition to promoting OPC maturation and differentiation during demyelination and remyelination.

**Graphical Abstract:** 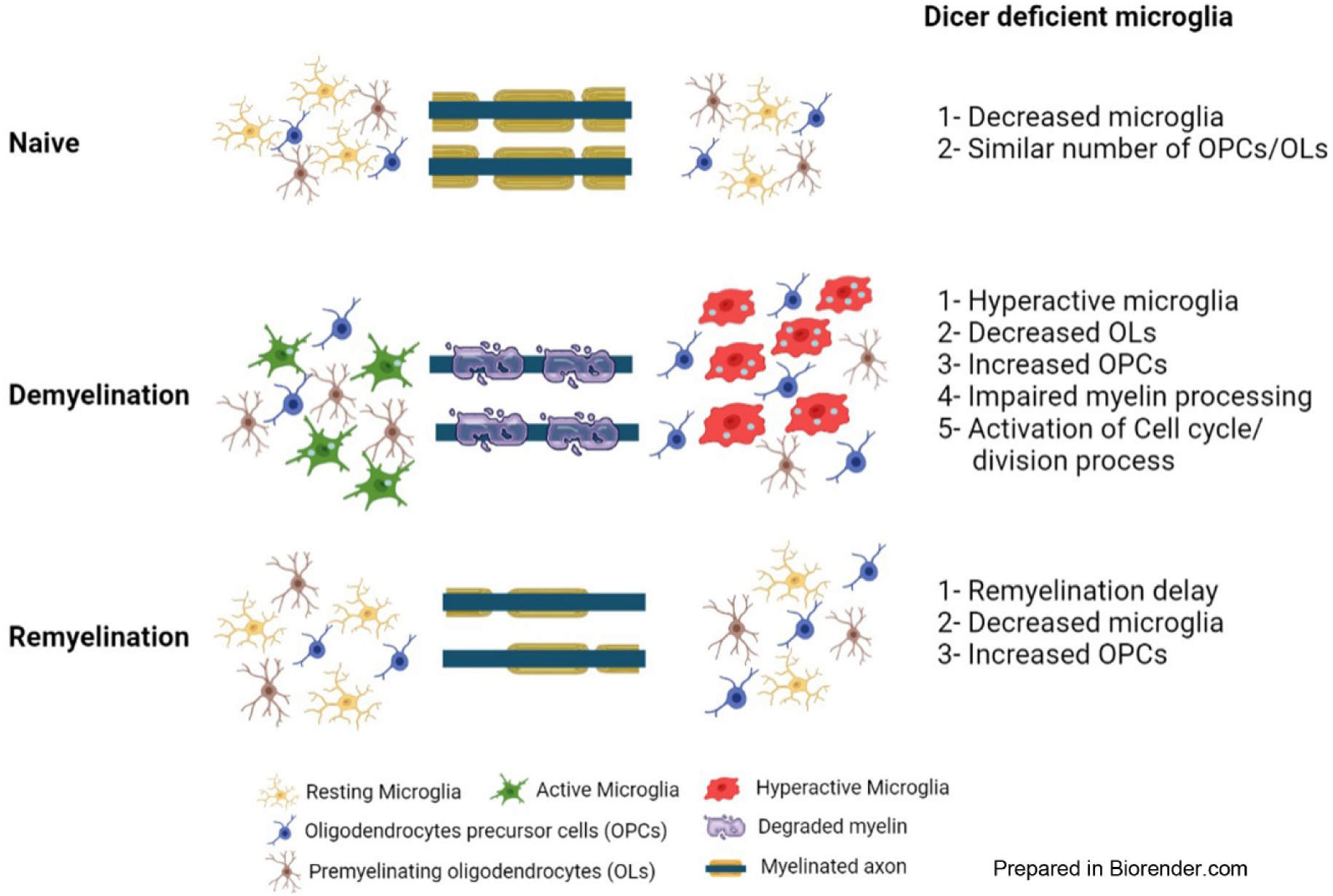

## Introduction

Microglia, the tissue resident macrophages of the central nervous system (CNS), arise from myeloid progenitor cells in the embryonic yolk sac. These innate immune cells play critical roles in normal brain development and homeostasis, including maintaining oligodendrocyte (OL) progenitor cell (OPCs) numbers and supporting OL maturation and myelination^1–3^. Recent studies have reported dysregulated microglial activity in various neurodegenerative diseases including multiple sclerosis (MS), a chronic inflammatory demyelinating disease of the CNS. In MS, OL damage, myelin destruction, and accompanying neuronal loss have been associated with activated microglia/macrophages in and around demyelinating lesions^4^. Nonetheless, microglia also repopulate remyelinating lesions and promote remyelination through the expression of anti-inflammatory molecules, phagocytosis of myelin debris, and repair of tissues^5–8^. Thus, identifying mechanisms regulating the pathogenic and protective functions of microglia is critical for designing therapeutic interventions for MS.

Emerging evidence suggests that epigenetic modulators (microRNAs (miRNAs) and histone modifications) regulate cellular behavior in both physiological and pathological conditions^9^. miRNAs, which are short non-coding RNAs, are processed to mature form by a ribonuclease type III, Dicer (DICER) enzyme, and regulate post-transcriptional gene expression by binding to the 3’UTR region of target genes. Various studies have identified critical direct and indirect roles for miRNAs in normal glial cell functions and development, including OL differentiation and maturation as well as axonal myelination/remyelination^10–13^. For example, astrocyte involvement in OL differentiation and myelination/remyelination has been found to be affected by miRNAs regulating target gene expression^14^. A recent study has also examined the importance of adult microglial miRNAs and reported that miRNAs are required for microglial responses to inflammatory challenges^15^. However, the effects of microglial miRNAs and their target genes in OL differentiation, myelination, and/or remyelination remain to be established.

In this study, we investigated the effect of microglia-specific *Dicer1* loss on the process of demyelination and remyelination. We found that loss of microglial *Dicer* expression did not alter adult CNS myelination. However, *Dicer*-deficient microglial cells assumed a hyperactive phenotype and caused extensive myelin loss following cuprizone-induced demyelination. Transcriptionally, *Dicer* deficiency in microglial cells led to dysregulation of pathways associated with cell cycle/cell division. Additionally, microglial myelin degradation and metabolizing capacity were also impaired, leading to diminished OPC differentiation and failure of remyelination.

## Results

### Adult microglial *Dicer* is dispensable for myelin maintenance

It has been previously reported that loss of postnatal microglial miRNA expression led to decreased microglial numbers in mouse brain cortex and hippocampal regions^15^. To test and distinguish the effect of tamoxifen (TMX) injection and loss of *Dicer* expression postnatally in CNS resident microglia/macrophages (Cx3cr1^creERT2^), we compared the morphology and microglial populations in brain white matter (WM) regions (corpus callosum, CC, **Supplementary Figure S1A**). IBA1-positive cell quantification showed that Cx3cr1^creERT2^ mutant animals had significantly less IBA1+ microglia in CC compared to littermate control (Cre−) animals (Cre− vs Cre+: 241±4.9 vs. 91.6±23.96, *p*=0.0036, **Figure 1A, 1B, and 1I**). Furthermore, three-dimensional morphometric analysis revealed no significant changes in cell volume or branch points/length in Cx3Cr1^creERT2^ mutant animals compared to controls (**Supplementary Figure S1B-S1D**) and they appeared as ramified, an indication of a resting phenotype. As Cx3cr1 is also expressed by peripheral immune cells and other tissue resident macrophages, we chose a CNS microglia-specific mouse line (Tmem119^creERT2^) to assess the effect of *Dicer* loss in resident microglia from the CC region^16^. Immunohistochemical staining analysis of TMX-treated Cre+ (mutant) animals showed similar decline in IBA1- positive microglia compared to the Cre− (control) group (Cre− vs Cre+: 175.9±11.41 vs. 138.2±8.27, *p*=0.0233, **Supplementary Figure S2A, S2B, and S2I**). Thus, these results suggest that loss of adult microglial miRNAs only affected the cell number, without affecting the level of cell activation.

**Figure 1:**
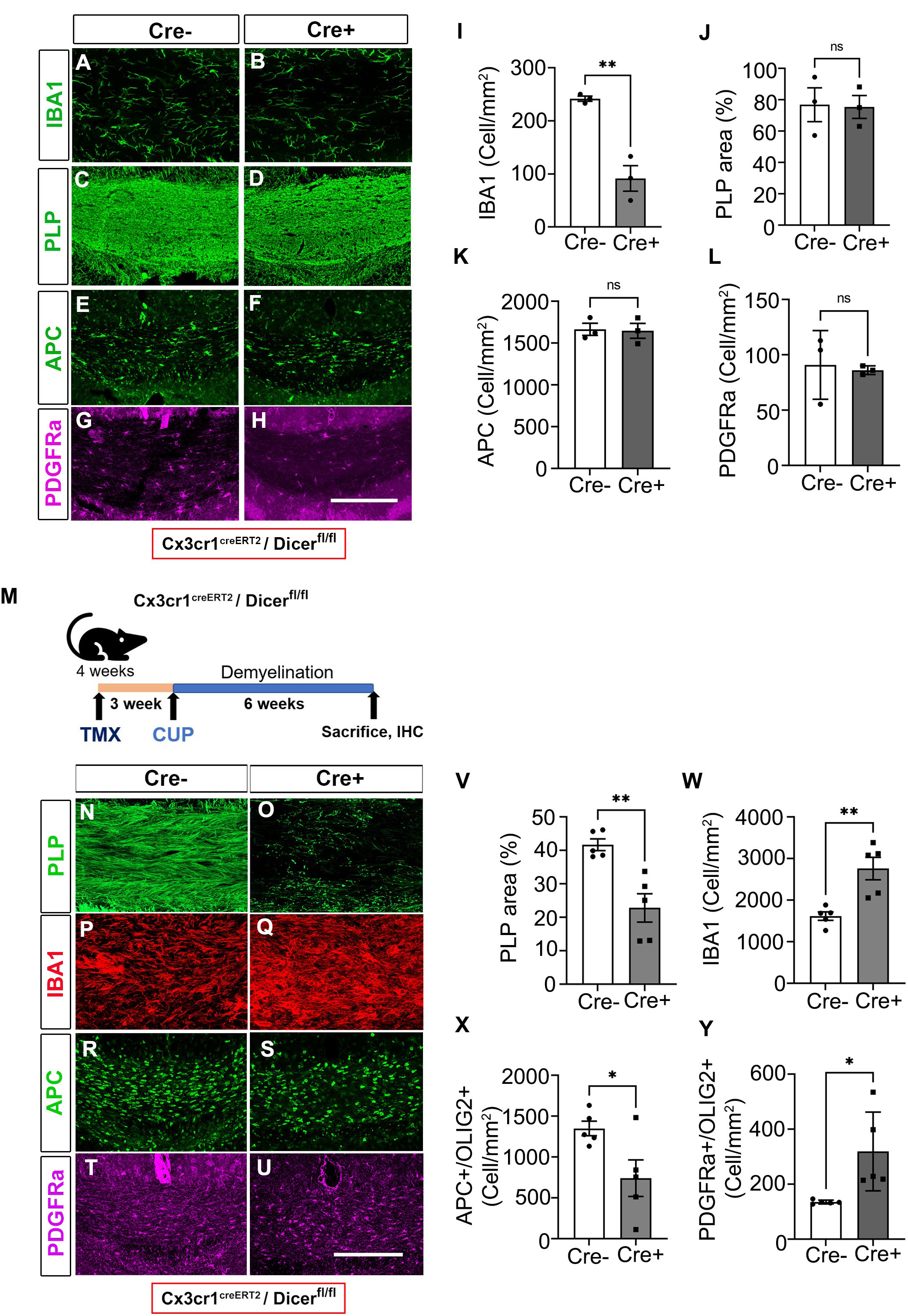
*Dicer1* is required for microglial maintenance and response to demyelination. **(A-H)** Representative immunostained confocal images of microglia (IBA1, A and B), myelin (PLP, C and D), OLs (APC, E and F), and OPCs (PDGFRα, G and H) from TMX- injected naïve Cre− control and Cre+ mutant animals. Scale bar - 100µm. **(I-L)** Bar graphs showing cell numbers (I, K and L) and myelin occupied area (J) in naïve mouse brain CC. Error bars indicate mean ± SEM. ** *p*< 0.01, ns- nonsignificant; Student’s *t*-test, two-sided. **(M)** Schematic illustration of Cx3Cr1^creERT2^/Dicer^fl/fl^ animals undergoing TMX treatment and CUP feeding. **(N-U)** Representative immunostained confocal images of myelin (PLP, N and O), microglia (IBA1, P and Q), OLs (APC, R and S), and OPCs (PDGFRα, T and U) from TMX-injected Cre− control and Cre+ mutant animals after six weeks of CUP feeding. Scale bar - 100µm. **(V-Y)** Bar graphs showing myelin occupied area (V) and cell numbers (W-Y) in CUP-fed mouse brain CC. Error bars indicate mean ± SEM. \**p*< 0.05, ** *p*< 0.01, ns- nonsignificant; Student’s *t*-test, two-sided.

To analyze how loss of microglial miRNAs affected myelin content and OL lineage cells (OPCs/OLs), we performed immunostaining for myelin (PLP), OLs (APC), OPCs (PDGFRα), and oligodendroglia lineage cells (OLIG2). Our results showed that the absence of miRNAs in microglia did not affect myelin content in either model of *Dicer* iKO animals (Cx3cr1^creERT2^ and Tmem119^creERT2^) compared to littermate controls (PLP-stained area, Cre− vs. Cre+: Cx3cr1^creERT2^-76.86±10.8 vs 75.41±7.26, *p*=0.9171, **Figure 1C, 1D, and 1J**; Tmem119^creERT2^- 57.17±2.58 vs. 54.36±3.08, *p*=0.0.5001, **Supplementary Figure S2C, S2D, and S2J**). Similarly, we did not find any significant change in cell numbers (OLs/OPCs) between the control and mutant groups in either genotype (Cx3cr1^creERT2^: APC+/OLIG2+, PDGFRα+/OLIG2+ and OLIG2+ cells, *p*= 0.8874, *p*= 0.8066 and *p*=0.497 and Tmem119^creERT2^: APC+, PDGFRα+ and OLIG2+cells, *p*= 0.8346, *p*= 0.9195, and *p*= 0.1639), respectively (**Figure 1E-1H, 1K and 1L, Supplementary Figure S1E-S1G, S2E-S2H, S2K and S2L**). Collectively, these findings indicate that deletion of *Dicer* expression in adult microglia does not significantly affect OL lineage cells or myelin content.

### *Dicer*-deficient microglia become hyperactive during demyelination

*Dicer-*ablated adult microglia show hyperactive response under inflammatory stimuli^15^. To examine microglial response during demyelination, we fed 0.3% cuprizone (CUP; a copper chelating agent)-supplemented diet to TMX-treated adult animals (Cre− and Cre+) for six weeks (**Figure 1M**). Comparing PLP immunostaining at the peak of demyelination showed significant myelin loss in Cx3cr1^creERT2^ mutant (Cre+) animals compared to control (Cre−) animals (PLP stained area, Cre− vs. Cre+: 41.69±1.7% vs 22.8±4.2%, *p*= 0.0033, **Figure 1N vs. 1O, and 1V**). Noticeable myelin loss was also found in CUP-fed Cre− control animals when compared to animals maintained on a normal diet (**Figure 1C vs. 1N**). As CUP-mediated demyelination has been correlated with gliosis (microgliosis and astrocytosis) due to increased cellular activity and cell proliferation^17^, PLP-stained serial sections were immunostained to identify microglial (IBA1) phenotype during demyelination (**Figure 1P vs. 1Q**). Strikingly, IHC showed a significantly increased number of microglia in mutant (Cre+) animals compared to controls (Cre− vs. Cre+: 1612.8±102 vs 2758±271, *p*=0.0042; **Figure 1W**).

Microglia accumulation in the demyelinated brain coincides with the upregulation of chemokines and cytokines, resulting in an inflammatory environment that propagates microglial proliferation and activation^18^. To confirm a significantly increased microglia population at the peak of demyelination, we investigated whether cell proliferation contributed to microglial expansion in Cx3cr1^creERT2^ mutant (Cre+) animals at the peak of demyelination. Mice were injected with BrdU for six consecutive days starting at the end of 5 weeks of CUP treatment before collecting brain tissue from experimental animals at the end of demyelination. Dual immunohistochemical staining with IBA1 and BrdU showed significantly increased proliferating microglia in Cx3cr1^creERT2^ mutant (Cre+) animals compared to control (Cre−) animals (Cre− vs. Cre+: 245±49 vs 1077±151, *p<*0.0001; **Supplementary Figure S3A-S3D**). Similarly, astrocytosis was also observed in CUP-fed animals; however, the hypertrophic phenotype of the astrocytes made it difficult to perform quantitative analysis in these two genotypes (data not shown).

Feeding CUP to mice triggers apoptosis in mature OLs while leaving the population of OPCs unaffected, and it does not alter the formation of new OLs during CUP-mediated WM (CC) demyelination^19^. Thus, to correlate enhanced demyelination of Cre+ mutant animals, we checked mature OLs with APC (CC1) labeling and found significantly less surviving OLs in microglia *Dicer* iKO animals (Cre+) compared to control animals (Cre− vs Cre+: 1349±89 vs. 739±224, *p*=0.0357; **Figure 1R vs 1S, and 1X**, **Supplementary Figure S4A-S4D**,). This loss was found to be associated with increased OLs apoptosis in *Dicer* mutant animals (**Supplementary Figure S4E-S4K**). Interestingly, we found significantly increased OPC numbers in Cre+ mutant animals compared to controls (Cre− vs. Cre+: Cx3cr1^creERT2^- 30.9±1.4 vs 72.1±12.9, *p*=0.0132; **Figure 1T vs. 1U, and 1Y,**). The previous finding of spontaneous remyelination occurring during ongoing demyelination during week six of CUP diet might be associated with increased OPCs in Cre+ mutant animals, which showed extensive demyelination^20^.

After CNS demyelination, microglia internalize and phagocytose myelin to degrade intracellularly by lysosomal structures^21^. Hence, we checked whether extensive myelin removal in Cre+ mutant animals was associated with increased myelin phagocytosis by hyperactive microglia and processing using the phagolysosomal protein, CD68, in CUP-fed mice (Ctr vs Cup-6w- IBA1:18.39±2.48 vs. 59.18±5.72, p=0.0023; CD68: 1.29±0.2 vs. 74.37±36.4, p=0.0253; **Supplementary Figure S4L-S4S)**. Quantitative confocal analysis of CD68+ and IBA1+ cells showed a significant increase in phagolysosome occupied area (Cre− vs Cre+: 12.6±1.4 vs 21.6±2.4, p=0.0079) and microglia population (Cre− vs Cre+: 42.3±2.6 vs 56.6±1.6, *p*=0.0015) in Cre+ mutants compared to control mice (**Figure 2A-2J**). This *in vivo* result was further confirmed with BV2 cells *in vitro*, as we observed impaired phagocytosed myelin processing in Crispr *Dicer*1 KO cells (**Figure 2K)**. We found increased MBP levels in *Dicer*-deficient BV2 cells after 24h of purified myelin treatment (**Figure 2L**, 1.9-fold change, *p*=0.0427) compared to Crispr control cells. Interestingly, endogenous *Dicer1*-deleted BV2 microglia also showed similar myelin processing impairments under inflammatory conditions (data not shown). Overall, these data suggest that loss of miRNAs from microglia, and the associated dysregulated gene expression, not only cause a hyperactive phenotype during demyelination, but these cells are also unable to metabolize and process phagocytosed myelin during demyelination.

**Figure 2:**
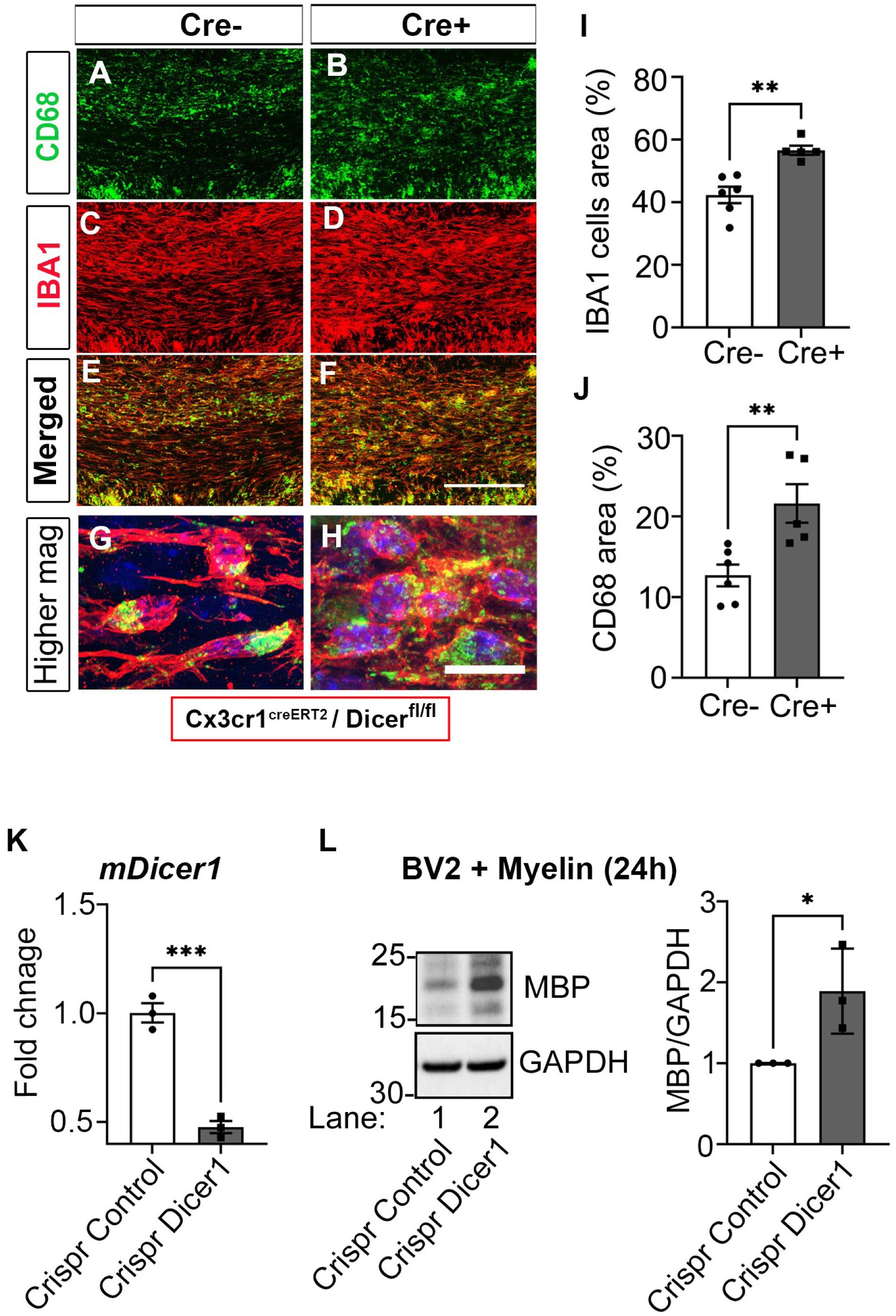
*Dicer1*-ablated microglia are hyperactive during demyelination and show impaired myelin processing. **(A-H)** Representative immunostained confocal images of phagolysosome (CD68, A and B)-positive microglia (IBA1, C and D) from TMX-injected Cre− control and Cre+ mutant animals fed CUP for six weeks. Merged images are shown in panels E and F. Higher magnified images show microglia (IBA1) contain phagolysosomes (G and H). Scale bars - 50µm (A-F) and 20µm (G and H). **(I and J)** Bar graphs showing CD68- and IBA1- occupied area in mouse brain CC. Error bars indicate mean ± SEM. ** *p*< 0.01; Student’s *t*-test, two-sided. **(K)** qPCR analysis of *mDicer1* expression in Crispr Control and Crispr Dicer1 BV2 cells. GAPDH was used as an endogenous control for qPCR analysis. Error bars indicate mean ± SEM. \*\*\**p*<0.001; Student’s *t*-test, two-sided. **(L)** Representative western blot images and quantification (from three independent experiments with similar results) of myelin-treated stable m*Dicer1* KO BV2 cells. GAPDH expression was used as a loading control. Data represent mean ± SEM of three independent experiments; \**p*<0.05; Student’s t-test, two-sided.

### Adult microglia *Dicer* ablation causes dysregulation of cell cycle pathways and activate genes related to disease-associated phenotype during demyelination

To evaluate the effect of miRNA loss on microglial gene expression in TMX-induced *Dicer* KO animals, adult Cx3cr1^creERT2^ mouse brain microglia (CD45^int^/Cd11b+) were flow-sorted and processed for RNA sequencing. Similar to the results from IHC staining (**Figure 1A, 1B and 1I**), we found significantly less microglia in Cre+ mutant animals in compared to controls after 8 weeks post-TMX administration (Cre− vs. Cre+: 206,338±23,383 vs. 94,594±9,885, p=0.0117; **Supplementary Figure S5A and S5B**). As expected, TMX-induced CRE-recombinase-mediated *Dicer* ablation led to a significant reduction in endogenous microglial *Dicer1* expression (−0.38-Fold, p<0.0001; **Supplementary Figure S5C**). To further ensure microglia-specific transcriptome enrichment, we checked cell-specific gene expression from other CNS cell types and found that enriched genes were mainly contributed by microglia/macrophages (**Supplementary Figure S5D**), thus confirming enriched microglial cell sorting. Comparative DEG analysis between Cx3cr1^creERT2^ Cre− control and Cre+ mutant animals revealed 594 upregulated and 554 downregulated genes (with >1 log2Fold change and p<0.01, **Figure 3A, Supplementary File 1**). Analysis of microglial cell-specific genes revealed a decreased expression in *Dicer1*-ablated microglial cells (**Figure 3B**), whereas in analyzing pro- and anti-inflammatory responses, we found no significant association to either phenotype in the absence of miRNA expression, respectively (data not shown). Further, functional categorization by the DAVID bioinformatics database^22^ for Gene Ontology (GO) annotation of enriched biological processes (p<0.05) showed altered genes to be associated with cell division/cycle, among others (**Figure 3C, Supplementary File 2**). Additionally, analyzing affected pathways with the ingenuity pathway analysis (IPA, p < 0.05) tool, we observed activated pathways (Z score >2) like Cardiac Hypertrophy Signaling (Enhanced), Insulin Secretion Signaling Pathway, Natural Killer Cell Signaling, Kinetochore Metaphase Signaling Pathway and many more, whereas pathways like Glutathione-mediated Detoxification, VDR/RXR Activation, PPARα/RXRα Activation, and the Xenobiotic Metabolism AHR Signaling Pathway were found to be inhibited (Z score <2) in *Dicer*- deficient microglia (**Supplementary File 3**).

**Figure 3:**
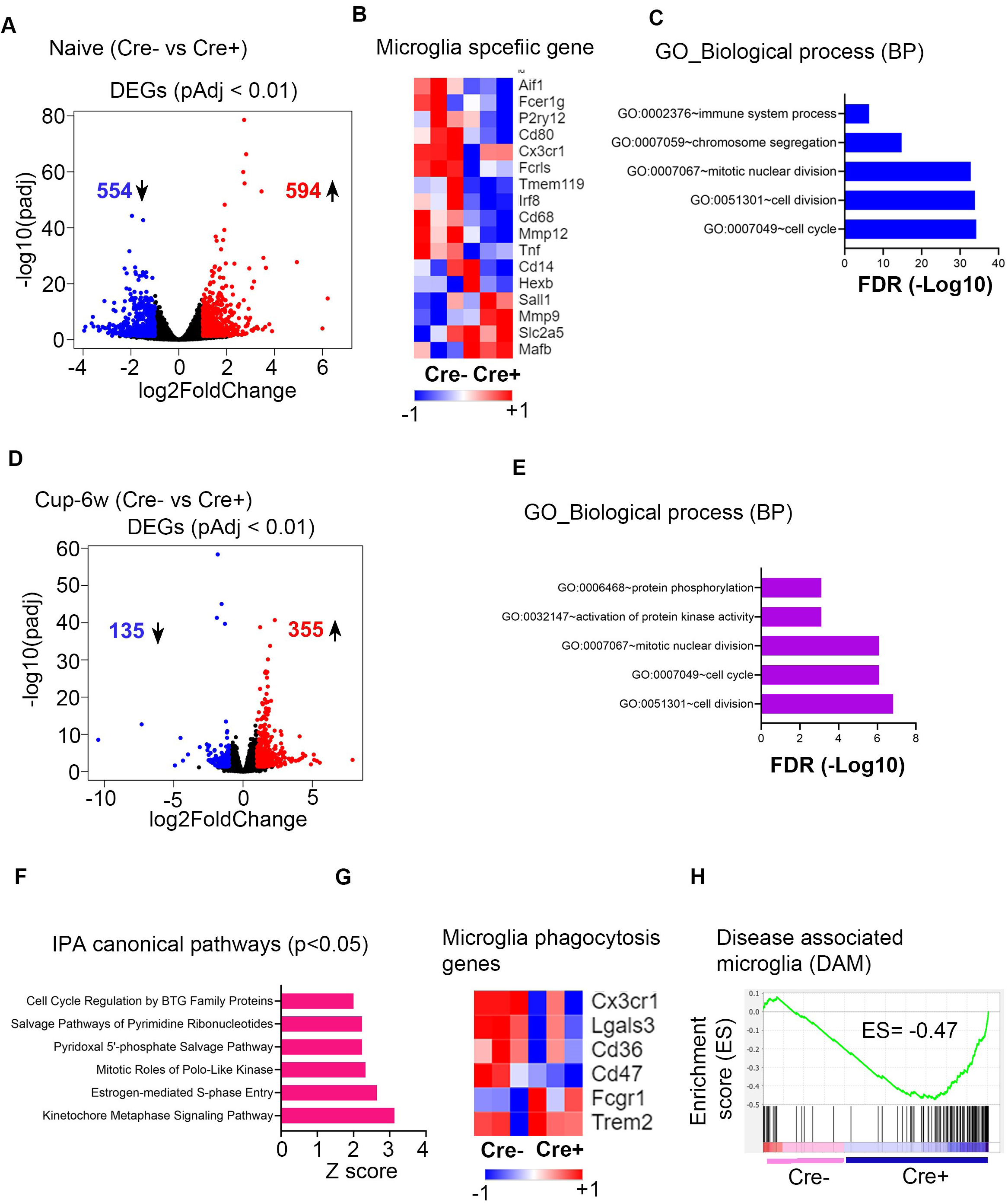
Microglial *Dicer1* is required for cell cycle activity and to regulate microglial disease phenotype. **(A)** Volcano plot showing significantly differentially-expressed genes (DEGs, log2Fold change >1 and *p*>0.01) in naïve *Dicer*-ablated adult microglia. Red dots represent upregulated transcripts and blue dots represent downregulated transcripts. **(B)** Heatmap showing microglia-specific transcript expression in *Dicer*-ablated adult mouse brain microglia. The scale indicates expression relative to CPM values across RNA samples. **(C)** The top five biological processes associated with significant DEGs of naïve adult *Dicer*-ablated microglia. **(D)** Volcano plot showing significantly differentially-expressed genes (DEGs, log2Fold change >1 and *p*>0.01) in *Dicer*-ablated adult microglia FACS sorted from CUP-fed animals. Red dots represent upregulated transcripts and blue dots represent downregulated transcripts. **(E)** The top five biological processes associated with significant DEGs from CUP-fed adult *Dicer*-ablated microglia. **(F)** The top IPA canonical pathways associated with significant DEGs from CUP-fed adult *Dicer*-ablated microglia. **(G)** Heatmap showing significantly regulated genes of microglial phagocytosis transcript expression in CUP-fed *Dicer*-ablated adult mouse microglia. The scale indicates expression relative to CPM values across RNA samples. **(H)** Gene set enrichment analysis showed increased proportions of transcripts associated with disease-associated phenotype in *Dicer*-ablated adult mouse microglia during demyelination. The x-axis with the pink line represents transcripts related to Cre− control microglia, while the purple line shows the presence of transcripts related to Cre+ mutant microglia (ES- Enrichment score is shown).

Next, we analyzed gene expression of flow-sorted hyperactive microglia during demyelination (CUP-6w) (**Supplementary Figure S5E**). Comparative DEG analysis showed 355 upregulated and 135 downregulated genes (with >1 log2Fold change and p<0.01, **Figure 3D, Supplementary File 4**) in Cre+ mutant animals compare to control groups (Cre−). GO term analysis of biological processes using the DAVID tool showed enrichment of processes like cell cycle, cell division, along with immune system process and immune response (**FDR< 0.01, Supplementary File 5**), in concert with earlier BrdU findings (**Supplementary Figure S3B-S3D**). Likewise, IPA canonical pathway analysis showed activation of Kinetochore Metaphase Signaling Pathway, Estrogen-mediated S-phase Entry, Mitotic Roles of Polo-Like Kinase, and others (Z score >2, p<0.05) in the absence of microglial miRNA expression during demyelination (**Figure 3F, Supplementary File 6**). Further, to confirm enhanced phagolysosome formation in *Dicer*1-ablated microglia during demyelination (**Figure 2**), we screened for genes related to phagocytic activity^23^ and found significant changes in transcript levels of *Cx3cr1*, *Lgals3*, *Cd36, Cd47, Fcgr1 and Trem2* (**Figure 3G**). As activated microglia attain a disease-specific phenotype in neurodegenerative diseases^24^, we compared microglia transcriptome signatures for the presence of genes specific to disease-associated microglia (DAM) during demyelination. Analyzing significantly dysregulated microglial genes using the Molecular Signatures database (v7.4) gene set enrichment analysis tool^25^ for the presence of DAM gene signatures, we found increased proportions of genes like *Cst7*, *Ctsd*, *Ctsz*, *Tyrobp*, and *Cadm1* in *Dicer*-deficient microglia (Cre+) compared to control animals (**Figure 3H**). Overall, RNA-seq data analysis indicated that enhanced microgliosis (cell number/hyperactive phenotype) during demyelination is an outcome of loss of miRNA-controlled target gene expression and upregulation of disease-associated transcripts in microglia.

### *Dicer*-deficient microglia inhibit OL differentiation and delay remyelination

Previous studies have reported that microglia contribute to OL differentiation and myelination by adopting a pro-regenerative phenotype and secreting growth factors^7,26^. Thus, to examine whether miRNA expression loss in microglia has any effect on OL differentiation, OPCs were co-cultured with *Dicer1*-inhibited primary microglia in OL differentiation media (**Figure 4A)**. After three days of differentiation, we found significantly less MBP+ cells in the presence of miRNA-deficient microglia (**Figure 4B and 4C**, siControl vs. siDicer1: 4.4±0.33 vs 3.3±.13, *p*=0.0052).

**Figure 4:**
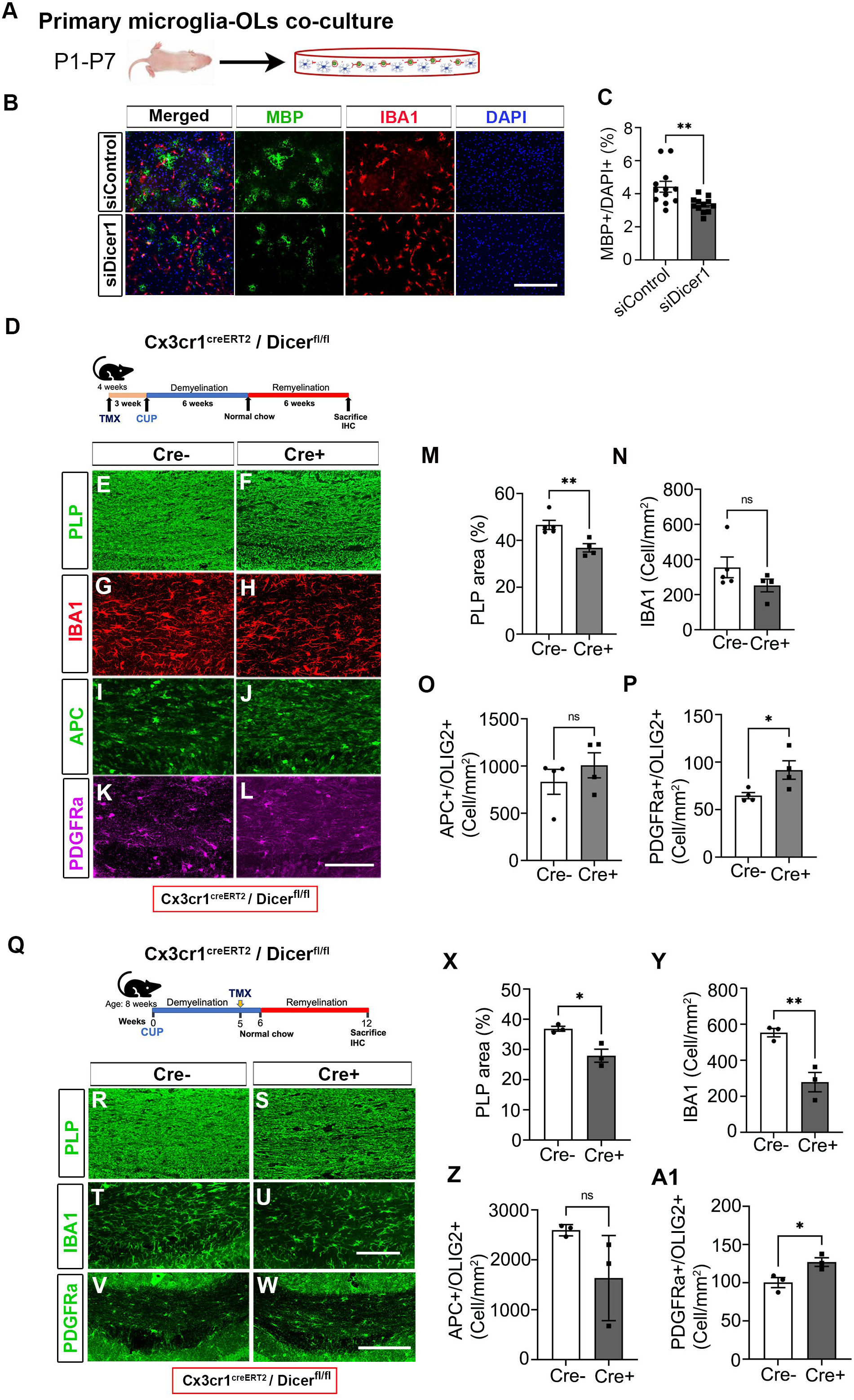
Microglia *Dicer1* is critical for successful OL differentiation and optimal remyelination. **(A)** Schematic illustration of siDicer1-treated primary microglia co-cultured with P7 mouse primary OPCs. **(B)** Representative immunofluorescence images (from three independent experiments with similar results) showing mouse primary OLs (MBP+ cells, green) and co-cultured with siDicer1 treated primary mouse microglia (IBA1+ cell, red). siControl- top row and siDicer1- bottom row. Nuclei were stained with DAPI (blue). Scale bar - 50µm. **(C)** Quantitative analyses of MBP+ cells from OL/microglia co-cultures. Data represent mean ± SEM of MBP+ cells populations from 3 independent experiments; \*\**p*<0.01; Student’s *t*-test, two-sided. **(D)** Schematic illustration of Cx3cr1^creERT2^/Dicer^fl/fl^ animals undergoing TMX treatment, CUP feeding, and remyelination. **(E-L)** Representative immunostained confocal images of myelin (PLP, E and F), microglia (IBA1, G and H), OLs (APC, I and J), and OPCs (PDGFRα, K and L) from TMX-injected Cre− control and Cre+ mutant animals after six weeks of CUP feeding and then 6 weeks maintained on a normal diet. Scale bar - 100µm. **(M-P)** Bar graphs showing myelin-occupied area (M) and cell numbers (N-P) in remyelinated mouse brain CC. Error bars indicate mean ± SEM. \**p*< 0.05, ** *p*< 0.01, ns- nonsignificant; Student’s *t*-test, two-sided. **(Q)** Schematic illustration of Cx3cr1^creERT2^/Dicer^fl/fl^ animals undergoing TMX treatment at 5 weeks of CUP feeding (6 weeks) and remyelination. **(R-W)** Representative immunostained confocal images of myelin (PLP, R and S), microglia (IBA1, T and U), and OPCs (PDGFRα, V and W) from TMX-injected (at five weeks of CUP feeding) Cre− control and Cre+ mutant animals after six weeks of CUP feeding followed by 6 weeks on normal diet. Scale bar - 100µm. **(X-A1)** Bar graphs showing myelin-occupied area (X) and cell numbers (Y-A1) in remyelinated mouse brain CC. Error bars indicate mean ± SEM. \**p*<0.05, \*\**p*< 0.01, ns-nonsignificant; Student’s *t*-test, two-sided.

Further, to examine the importance of microglia-specific miRNA expression during remyelination, TMX-injected control (Cre−) and mutant (Cre+) animals from both genotypes were transferred to normal diet for six additional weeks following six weeks of CUP-induced demyelination (**Figure 4D**). At the end of the remyelination period, brain tissue was collected from both groups and processed for immunohistochemical analysis. We started by analyzing the effect of *Dicer* loss on microglia during remyelination (**Figure 4G vs. 4H**) and we observed a decreased trend of IBA1-positive cells in Dicer KO animals compared to the control group (Cre− vs. Cre+: 355±59 vs 251±36, *p*=0.206; **Figure 4N**). To further examine remyelination, serial brain sections were immunostained for PLP (**Figure 4E vs. 4F**). Comparing between the two groups, we found significantly smaller PLP-stained areas in Cre+ mutant mice than in Cre− control animals (Cre− vs. Cre+: 46.6±1.9 vs 36.8±1.8, *p*=0.0086; **Figure 4M**). The reduced PLP staining was not found to be associated with axonal loss, as we did not observe any difference in axonal neurofilament, NF200, staining (data not shown). To understand whether decreased PLP staining (remyelination delay/failure) in Cre+ mutant animals was related to differences in the numbers of mature OLs, we immunostained serial brain sections with APC and found no significant differences in OL cell numbers between the two groups (**Figure 4I vs. 4J, and 4O**, *p*=0.3844). On the contrary, we observed significantly more OPCs in Cre+ mutant animals compared to the control group at end of the remyelination period despite similar OL lineage cell (OLIG2+) numbers in the two groups (Cre− vs. Cre+: 64.7±3.2 vs. 91.7±9.8, *p*=0.0394; **Figure 4K vs. 4L, and 4P, Supplementary Figure S6A-S6D**). Furthermore, to gain insight into microglial gene expression changes in the absence of miRNA expression during remyelination, microglia were FACS-sorted and processed for RNA-seq analysis (**Supplementary Figure S6E)**. Comparative DEGs analysis showed only 27 upregulated and 8 downregulated genes (with >1 log2Fold change and p<0.01, **Supplementary Figure S6F, Supplementary File 7**) in Cre+ mutant animals compared to control groups (Cre−). GO term analysis associated with biological process showed significant enrichment (p<0.01) of terms like response to heat (GO:0009408), cell adhesion (GO:0007155), response to unfolded protein (GO:0006986), cell surface receptor signaling pathway (GO:0007166), and negative regulation of angiogenesis (GO:0016525) (**Supplementary Figure S6G**, **Supplementary File 8**).

To further exclude effects of residual degraded myelin or impaired microglial myelin processing (**Figure 2**), we induced microglial *Dicer1* ablation in a separate cohort of animals where TMX was intraperitoneally administered to mice after 5 weeks of CUP feeding (**Figure 4Q**). At the end of TMX treatment, animals were transferred to a normal diet for an additional six weeks to remyelinate. At the end of the specified time, mice were euthanized and brain tissue was processed for IHC analysis. Comparing control (Cre−) and Cre+ mutant animals after 6 weeks of remyelination, we found significantly less PLP staining in Cre+ mutant CC compared to controls (**Figure 4R vs. 4S**; Cre− vs. Cre+: 36.9±0.8 vs 27.9±2.2, *p*=0.0183; **Figure 4X**). Interestingly, we also observed significant less IBA1+ cells in Cre+ mutant animals at the end of the remyelination period (**Figure 4T vs. 4U**; Cre− vs. Cre+: 553±23 vs 278±53, *p*=0.0094; **Figure 4Y**), highlighting the role of miRNAs in maintaining required microglial cell numbers in the mouse brain. We further examined the numbers of OLs in serial sections using APC immunostaining and found a decreasing trend in the number of mature OLs in Cre+ mutant animals (Cre− vs. Cre+: 2592±66 vs 1634±492, *p*=0.1253; **Figure 4Z**, **Supplementary Figure S7A-S7C**). In addition, we also observed significantly more PDGFRα+ cells in Cre+ mutant animals compared to controls at the end of six weeks of remyelination (**Figure 4V vs. 4W**; Cre− vs Cre+:100±6.6 vs. 127±5.7, *p*=0.0371; **Figure 4A1**;). To identify microglial genes reported to play role in OL differentiation/maturation/myelination/remyelination, sorted microglia were processed for RNA-seq analysis as described earlier. Comparative data analysis showed a total of 460 significantly dysregulated DEGs (293 upregulated and 167 downregulated, log2FoldChange> 1 and p<0.01) between Cre− control and Cre+ mutant animals (**Supplementary Figure S7D, Supplementary File 9**). Interestingly, the absence of miRNAs during remyelination caused significant downregulation of homeostatic markers like *Cx3cr1* and *Serpine2* (**Supplementary Figure S7E**), which might be associated with impaired remyelination after CUP treatment^27^. In addition, *Dicer*-deficient microglia also showed prominent enrichment of biological processes associated with cell division activity like regulation of attachment to spindle microtubule kinetochore, DNA replication, and others as evidenced by upregulation of *Nek2*, *Racgap1*, *Ect2*, *Knstrn,* and *Spag5* (**Supplementary Figure S7F and S7G, Supplementary File 10**). Overall, these results suggest that microglial *Dicer1-*mediated miRNA expression is critical for successful remyelination.

## Discussion

Various studies have reported cell intrinsic/extrinsic effects of miRNAs on target gene regulation in the contexts of both normal cell function and disease conditions, including a recent finding suggesting that miRNA target gene regulation contributes to intercellular cell signaling and thus affects proper functioning and development of target cells^28^. In this study, we demonstrated the importance of microglial miRNA expression in normal brain function and during ongoing demyelinating/remyelinating conditions using two independent inducible *Dicer*-ablated microglial mouse models of MS disease. *Dicer*- ablated adult microglia did not affect either brain myelin content or cause changes in OPCs, despite the reduction in IBA1-positive cells in both genotypes. However, CUP-mediated demyelination led to hyperactive microglia with impaired myelin processing ability. Transcriptionally, microglial *Dicer* loss led to increased expression of cell cycle/cell division genes in both healthy and demyelinating conditions, respectively. Interestingly, impaired gene expression due to loss of *Dicer* in microglia led to attaining a pathogenic phenotype, specifically in the demyelinating condition. Intriguingly, microglial miRNAs were also required for the OPC differentiation and maturation and necessary for timely and successful remyelination, irrespective of the stage of *Dicer* ablation. Thus, the results reported here highlight optimal microglial miRNA-mediated cell intrinsic gene regulation required for clearing myelin, as well as cell extrinsic effects on OLs by promoting differentiation/maturation required for successful remyelination in MS disease.

*Dicer1*, a terminal endonuclease, is responsible for generation of mature miRNAs and is associated with post-transcriptional regulation of target gene expression. *Dicer*- dependent miRNAs have been reported to play roles in various aspects of cell development like cell cycle progression, cell division, cell maturation, and survival processes, including neurons, glia, and immune cells^29–31^. In the context of reduced microglial cell numbers in naïve *Dicer* KO mouse brains, recent finding of miRNA-dependent DNA repair and preservation of genome integrity and *Dicer*-dependent cell survival may explain the differences with the control group^15,32^. Interestingly, IPA analysis of RNA-seq data also showed dysregulated expression of genes related to cell cycle/division/DNA damage checkpoint regulation functions (*Ccne1, Ccne2, E2f1, E2f7, E2f8, Myc, Aurka, Ccnb1, Ccnb2, Cdc25c, Pkmyt1, Plk1*) ^33,34^. A previous detailed study from Varol and colleagues also reported a similar decline in adult microglia numbers in conditional *Dicer* KO mouse brains^15^. Future studies will be needed to confirm whether similar declines in microglia numbers are present during early postnatal periods.

In demyelinating diseases like MS, degraded myelin accumulates in the lesion area and phagocytosed cells like microglia and infiltrating macrophages clear myelin debris, which is a prerequisite process for a successful repair process^35–37^. We observed heightened myelin loss and increased phagolysosome content at the peak of demyelination in *Dicer* iKO animals, along with impairment of *Dicer*-ablated microglia to process phagocytosed myelin, suggesting that *Dicer*-mediated miRNA expression is a prerequisite for optimal microglial activity in such an environment. To support this concept, RNA-seq analysis from sorted microglia at 6 weeks of demyelination showed significant dysregulation in genes (*Cx3cr1, Lgals3, Cd36, Cd47, Sirpa*) associated with phagocytic function^38,39^. Interestingly, Cx3cr1 deficiency in mice significantly reduced the clearance of myelin and inhibited proper remyelination after CUP treatment, highlighting the association between myelin clearance and demyelination/degenerative mechanisms^6^. In addition, GSEA analysis also confirmed that *Dicer*-ablated microglia acquire a hyperactive disease-associated transcriptional state^40^, which could be associated with the impaired myelin processing ability of microglia as DAM response is required for optimal lysosomal degradation and lipid processing, among other activities^41^.

Clearance of myelin debris, followed by recruitment and proliferation of OPCs and differentiation into mature myelinating OLs, are critical steps for successful remyelination in demyelinating diseases like MS^42^. We noticed increased OPC numbers during demyelinating and remyelinating stages in *Dicer* iKO animals. As the presence of degraded myelin impairs the recruitment and differentiation of OPCs^21^, excessive demyelination and leftover degraded myelin might be responsible for arresting OPCs at the precursor stage in the lesion area, leading remyelination delay/failure in *Dicer* iKO animals. Furthermore, *in vitro* microglia/OPC co-cultures showed significantly fewer differentiated OLs when microglia *Dicer1* expression was inhibited using a pool of siRNAs. The role of microglia in myelination/remyelination has not only been related to myelin debris clearance, but failing paracrine stimulation of OL differentiation is another key factor contributing to inefficient remyelination^43–45^. While looking for microglia-specific myelinating factors in the RNA-seq data, we found only a few associated transcripts (e.g. *Tgm2*) in remyelination cohorts. The absence of microglia-specific OL differentiation/myelination factors might be associated with sample selection during the cell sorting stage, as we took a whole-brain microglia sorting approach during bulk RNA-seq analysis, rather than focusing on only the affected CC (WM) region. Future studies using single-cell RNA sequencing approaches from affected CC regions of demyelinating/remyelinating cohorts has been planned to probe for microglia-specific factors.

Collectively, we demonstrate the importance of miRNAs in maintaining an optimal CNS resident immune cell population and regulating activation in the healthy brain. On the contrary, in neurodegenerative diseases like MS, miRNAs regulate microglial disease phenotype and promote microgliosis. Furthermore, using *in vivo* Dicer-regulated miRNA expression, we showed that microglial miRNAs are required for myelin phagocytosis and its optimal processing. The presence of degraded myelin during remyelination stalls OPCs at the precursor stage and delays efficient remyelination. As *Dicer* expression has been found to be dysregulated in different biological samples from MS patients ^46,47^, it would be of great interest to consider optimal miRNA levels when designing any specific remyelinating therapeutic intervention for this disease.

## Acknowledgements

This work was supported by NINDS R01 NS123532 and NINDS R21 NS123546 to RD. The authors would like to acknowledge editorial support from Dr. Christopher Nelson.

## Author Contributions

AT, NR, RD designed the study. AT, NR, AP and CJ performed the experiments and analyzed most of the data. AT, NR, AP, CJ and RD wrote the manuscript.

## RESOURCE AVAILABILITY

### Lead Contact

Further information will be addressed by the Lead Contact, Ranjan Dutta.

### Materials Availability

For further requests for reagents, please contact the Lead Contact, Ranjan Dutta.

### Data and code Availability

This study did not generate datasets or codes.

### Experimental mouse models

All procedures were approved by the Institutional Animal Care and Use Committee (IACUC) at the Lerner Research Institute (LRI), Cleveland Clinic Foundation (Cleveland, OH). Wild-type (C57BL/6J, Stock #00664) and transgenic mice (CXrCR1^creERT2^, stock# 020940; Tmem119_creERT2_, stock# 031820; and Dicer^fl/fl^, stock# 06001) were procured from Jackson Laboratory and maintained on a 12h light/dark cycle with access to food and water *ad libitum*. All mouse experiments were performed from CNS tissue collected from P1-adult stage.

### Transgenic animal models and genotyping Dicer Floxed recombined alleles

The inducible Cx3cr1^creERT2^ /Tmem119^creERT2^xDicerf^l/fL^ (iKO) mouse line was generated by breeding Cx3cr1^Cre−ERT2^ /Tmem119^creERT2^ and Dicer^fl//fl^ mice and genotyped using primers targeting CRE insert and flox site **(Supplementary Table 1)**. Littermate CRE-negative (*Cx3cr1 Cre−: Dicer^fl/fl^ and Tmem119 Cre−: Dicer^fl/fl^*) animals were used as controls for *in vivo* experiments and labeled as Cre(-) or control, whereas mutant (*Dicer1* ablated), CRE positive (*Cx3cr1 Cre+: Dicer^fl/fl^ and Tmem119 Cre+: Dicer^fl/fl^*), were labeled as Cre(+), mutant.

### Tamoxifen treatment

To induce recombination in transgenic mice, TMX was suspended in warm corn oil (Sigma, # T5648, 10mg/ml) and administered intraperitoneally for five constitutive days at 75mg/kg BW. All animals were TMX-treated at 4-5 weeks of age and examined for 3-4 weeks before starting cuprizone (CUP) feeding.

### Cuprizone (bis–cyclohexanone-oxaldihydrazone)-induced demyelination

Seven-to eight-week-old animals were maintained on 0.3% cuprizone (CUP) supplemented diet (ENVIGO) for 6 weeks. For controls, littermate animals were maintained on normal chow. After 6 weeks, animals were euthanized and brain samples were collected. For remyelination groups, animals were transferred to normal chow for additional six weeks.

### BrdU incorporation assay

To identify proliferating glial cells during demyelination, BrdU (100μg, Sigma) was administered intraperitoneally for six consecutive days at five weeks of cuprizone treatment ^48^. At the end of the demyelination period (2 hrs post-last BrdU injection), animals were euthanized and brain samples were collected for IHC analysis^49^.

### Immunohistochemistry

Cre−/+ animals were euthanized and transcardially perfused with 4% paraformaldehyde (PFA). Brains were fixed in 4% PFA (48 hr) followed by equilibration in 30% sucrose solution for 48 hr. Subsequently, brains were coronally sliced with a cooled microtome into 30µm-thick sections. Sections were stained with rat anti-PLP (1:300, Gift from Dr. Wendy Macklin), rabbit anti-IBA1 (1:750, Abcam), mouse anti-APC (1:250, Millipore), goat anti-PDGFRα (1:250, R&D system), rabbit-cleaved caspase 3 (1:250, Cell signal), mouse anit-GFAP (1:2500, Millipore), rat anti-CD68 (1:500, BioLegend), rat anti-BrdU (1:250, Abcam), or rabbit anti-TMEM119 (1:500, Abcam) as previously described^50^. Primary antibody incubations were followed by corresponding secondary antibodies labelling using respective Alexa fluor-labelled donkey anti-rat/rabbit/mouse/goat (Thermo Scientific) antibodies.

### BV2 cell cultures and primary microglia cultures

BV2 cells (kindly provided by Dr. Feng Ling, Cleveland Clinic) were cultured in Dulbecco’s modified Eagle’s medium (DMEM) high glucose containing 5% fetal bovine serum (FBS), 100 IU/mL penicillin (pen), and 100 mg/mL streptomycin (strep) and incubated at 37^0^C and 5% CO_2_. Mouse brain primary microglia were isolated from P1 pups of either sex by mixed glia culture methods. Briefly, pup brains were dissected and cut into small 1 mm^3^ pieces before digesting in HBSS containing 200U papain and 2500U DNAseI at 37^0^C for 30 min. The cell suspension was filtered through a 40µm-strainer and re-suspended in growth media (DMEM+10%FBS+1x Pen-strep). The mixed glia cells were maintained in growth media at 37 ^0^C and 5% CO_2_ with media changes every other day until day 10-12. To collect loosely-attached microglia cells, flask was shaken at 150rpm for 1hr before transferring media with detached microglia cells to 12 mm poly-D-lysine-coated coverslips in a new 24-well tissue culture plate.

### Primary microglia siRNA transfection

Primary microglia were transfected by Dicer1 siRNA (Dharmacon siRNAs SMARTpool) and Lipofectamine RNAiMAX Transfection Reagent following manufacturer’s instructions using a siRNA to Lipofectamine ratio of 1:2. Before microglia transfection, fresh growth media was added to cells and transfected with 5 pmol siRNAs per well of a 24-well plate. Scrambled siRNA was used as an experimental control.

### Lenti-Dicer1 Crispr/Cas9 generation

Mouse *Dicer1* guide RNA (gRNA), AACCGTACACTGTCCATCGG, was designed using the ChopChop online guide RNA design tool^51^ and cloned in Lenti-CRISPR-Cas9 plasmid (lentiCRISPR v2 was a gift from Feng Zhang, Addgene plasmid # 52961; http://n2t.net/addgene:52961; RRID:Addgene_52961) as previously reported^52^. To generate lentivirus expressing *Dicer1* gRNA, cloned lentiCRISPR plasmid and 2^nd^ generation packaging system (psPAX2 and pCMV-VSV-G) were co-transfected to HEK cells using Lipofectamine 3000 transfection reagent (Thermo Fisher) as per the manufacturer’s protocol (psPAX2 was a gift from Didier Trono (Addgene plasmid # 12260; http://n2t.net/addgene:12260; RRID: Addgene_12260, pCMV-VSV-G was a gift from Bob Weinberg (Addgene plasmid # 8454; http://n2t.net/addgene:8454; RRID: Addgene_8454). After 24 hr post-transfection (day 0), fresh growth media (DMEM GlutaMax+10% FBS+1x antibiotics) was added and lentivirus particles containing media were collected over the following two days (day 1 and day 2) and further concentrated using an ultracentrifugation method as previously reported^13^.

### Stable BV2 *Dicer* knockout cell line

To create the *Dicer1* knockout cell line, BV2 cells were transduced with lentivirus expressing *Dicer1* gRNA-Cas9 using polybrene (10µg/ml, Sigma) transfection reagent as previously described^13^. Transduced BV2 cells were further selected with puromycin antibiotic (4µg/ml) to create a stable cell line. For controls, lentiCripsrV2-Cas9 empty vector expressing lentivirus was used.

### Myelin phagocytosis

For the phagocytosis experiment, myelin was isolated from adult mouse brain using a percoll density gradient method^53^. To analyze myelin uptake by Dicer1 KO BV2 cells, myelin (10µg/ml) was added to the culture medium. After 24 hr incubation, cells were lysed and protein was isolated for western blot analysis.

### Mouse brain primary OPC cultures

Mouse brain primary OPCs were isolated from P6-P7 mouse pups of either sex by an immunopanning method as previously described^13^. Briefly, CD140a-positive cells were serially immunopanned to enrich OPCs and cultured in serum-free OPC proliferation medium (DMEM, glutamine- 2 mM, sodium pyruvate- 1 mM, insulin- 5 µg/ml, n-acetyl-cysteine- 5 µg/ml, trace element B-1X, d- biotin- 10 ng/ml, B27- 1X, antibiotic-antimycotic- 1X, BSA-10 µg/ml, transferrin-10 µg/ml, putrescine- 1.6 µg/ml, progesterone- 60 ng/ml, selenite- 40 ng/ml, PDGFα- 20ng/ml, NT3- 2ng/ml, forskolin- 4.2 µg/ml, CNTF-10 ng/ml) and seeded on a microglia monolayer cultured on poly-D-lysine-coated 12-mm diameter glass coverslips in a 24-well plates (50,000 cells/coverslips). After 24 hr in proliferation conditions, old media was replaced with fresh OL differentiation media (OPC media +T3 (40ng/ml).

### RNA extraction and reverse transcription quantitative polymerase chain reaction (RT-qPCR)

Total RNA was isolated from sorted primary mouse microglia (CD45low/Cd11b+ cells) and stable BV2 cells using Qiagen RNA isolation kits (Qiagen) as per the manufacturer’s instructions. Total RNA was reverse-transcribed to cDNA by SuperScript™ VILO™ cDNA Synthesis Kits (Applied Biosystems) as recommended, respectively. The expression of *Dicer1* was assessed using TaqMan Gene expression assays (ThermoFisher). GAPDH was used as an endogenous control in the reaction. Delta Ct values were used to determine relative expression changes (2^-ΔΔCT^) and are presented as fold change (FC).

### Western blotting

Total protein was extracted from transfected primary BV2 cells in RIPA lysis buffer (Thermo Fisher) supplemented with 1X Halt protease and phosphatase inhibitor cocktails (Thermo Fisher). Ten micrograms (10µg) of total protein was resolved on 4-12% SDS-PAGE gels and transferred to PVDF membranes. Membranes were blocked in 5% non-fat dry milk in tris-buffered saline with Tween-20 (TBST) for 1 hr at room temp followed by incubating membranes overnight at 4^0^C in the following primary antibodies: rat anti-MBP (1:500) and mouse anti-GAPDH (1:5000). After washing in TBS-T, blots were then incubated with peroxidase-conjugated anti-mouse (1:10000) and anti-rat IgG (1:1000) for 1 hour at RT. Chemiluminescence bands were detected with Clarity Western ECL substrate (Bio-Rad Laboratories, Hercules, CA) and imaged using a Bio-Rad Chemidoc MP and analyzed using Image Lab software (ver. 5).

### Immunofluorescence

Fixed cells (OLs/microglia) were permeabilized with 0.1% Triton-X100 for 10 min at room temperature. After washing in PBS (3X), samples were blocked in 5% normal goat serum (NGS) and 1% BSA for 1 hr at RT. Samples were incubated with primary antibodies (MBP-1:300 and IBA1-1:500) overnight at 4^0^C. The next day, after washing in PBS (3X), cells were incubated in Alexa fluorophore-tagged compatible secondary antibodies for 1 hr at room temp. After washing (PBS), cells were mounted in prolong gold antifade (±DAPI) mounting media.

### Image acquisition and quantification

Fluorescent images were obtained with a Leica DM5500 fluorescence microscope (Leica, New Jersey, USA) and a confocal laser-scanning microscope (Zeiss, Oberkochen, Germany) at appropriate excitation wavelengths and magnification. For statistical analyses, representative images from comparable locations were acquired for each animal (control and mutant). Images were processed and quantified using FIJI (IMAGE J) software^54^.

### Microglia fluorescence-activated single cell sorting (FACS)

Brains from control, demyelination (CUP-6w), and remyelination (6+6w) mice were homogenized using a Neural Tissue Dissociation Kit (Papain, Miltenyi Auburn, CA) following the manufacturer’s protocol. Cells were re-suspended in RPMI containing 25mM HEPES (pH-7.2), adjusted to 30% Percoll (Pharmacia, Uppsala, Sweden), underlaid with 1 ml 70% Percoll and centrifuged at 850xg for 30 min at 4^0^C. Cells were collected from the 30%/70% interface, washed with RPMI, counted, and suspended in FACS buffer (0.1% Bovine serum albumin in DPBS). Fcγ receptors were blocked with 1% mouse serum and rat anti-mouse CD16/32 mAb (clone 2.4G2: BD Biosciences, San Diego, CA) for 20 min on ice prior to staining with CD45 (clone 30-F11) and CD11b (clone M1/70) antibodies in FACS buffer. Microglia were sorted based upon their CD45^int^CD11b+ phenotypes using a FACS Aria III (BD Biosciences) and FACS Diva software (BD Biosciences). Sorted cells were collected in 1.7ml Eppendorf tubes containing RNase inhibitor before subsequent centrifugation. Cell pellets were re-suspended in Trizol (Invitrogen, Carlsbad, CA) and stored at −80^0^C until RNA-seq.

### RNA-seq and data analysis

RNA was isolated from FACS microglia (CD45^int^/CD11b+) and quantified using a Qubit 2.0 Fluorometer (Life Technologies, Carlsbad, CA, USA). RNA integrity was checked with 2100 TapeStation (Agilent Technologies, Palo Alto, CA, USA). SMART-Seq v4 Ultra Low Input Kit for Sequencing was used for full-length cDNA synthesis and amplification (Clontech, Mountain View, CA), and an Illumina Nextera XT library was used for sequencing library preparation. The final library was assessed with Qubit 2.0 Fluorometer and Agilent TapeStation. The samples were sequenced using a 2x150 Paired End (PE) configuration on the Illumina HiSeq instrument according to the manufacturer’s instructions. Image analysis and base calling were conducted by the HiSeq Control Software (HCS) on the HiSeq instrument. Raw sequence data (.bcl files) generated from Illumina HiSeq were converted into fastq files and de-multiplexed using Illumina bcl2fastq v. 2.17 program. One mis-match was allowed for index sequence identification. After demultiplexing, sequence data was checked for overall quality and yield. Then, sequence reads were trimmed to remove possible adapter sequences and nucleotides with poor quality using Trimmomatic v.0.36. The trimmed reads were mapped to the *Mus musculus* mm10 reference genome available on ENSEMBL using the STAR aligner v.2.5.2b. The STAR aligner uses a splice aligner that detects splice junctions and incorporates them to help align the entire read sequences. BAM files were generated as a result of this step. Unique gene hit counts were calculated by using featureCounts from the Subread package v.1.5.2. Only unique reads that fell within exon regions were counted. After extraction of gene hit counts, the gene hit counts table was used for downstream differential expression analysis. Using DESeq2, a comparison of gene expression between the groups of samples was performed. The Wald test was used to generate p-values and Log2 fold changes. Genes with adjusted p-values < 0.05 and absolute log2 fold changes > (±) 1 were called as differentially expressed genes (DEGs) for each comparison.

### Quantification and statistical analysis

All data analyses were performed using GraphPad Prism 8.0. Quantifications were performed from at least three independent experiments and in a blinded fashion. Statistical analysis was performed using Student’s *t*-tests to compare between two groups and ANOVAs with Dunnett’s *post hoc* test (one-way)/ Bonferroni’s (two-way) for two or more samples compared to controls, respectively. *P*<0.05 was considered to be statistically significant. Data are shown as mean ± standard error of mean (s.e.m.).

## Supplementary Figures

**Supplementary Figure S1: Microglial morphology and volume measurements**

**(A)** Schematic illustration of Cx3Cr1^creERT2^/Dicer^fl/fl^ animals undergoing TMX treatment and cartoon of a coronal mouse brain section showing confocal imaged area for this study. **(B and C)** Representative IMARIS software-isolated single microglia cells (green, top and purple, bottom) from naïve TMX-injected Cx3cr1^creERT2^ Cre−control and Cre+ mutant mouse brain corpus callosum. Scale bar - 1µm. **(D)** Bar graphs showing microglial cell volume in naïve TMX-injected Cx3cr1^creERT2^ Cre− control and Cre+ mutant mouse brain corpus callosum. Error bars indicate mean ± SEM. ns- nonsignificant; Student’s *t*-test, two-sided. **(E and F)** Representative immunostained confocal images of OL (OPCs+OLs) population (Olig2+) from naïve TMX-injected Cx3cr1^creERT2^ Cre− control and Cre+ mutant animals. Scale bar - 100µm. **(G)** Bar graphs showing OL cell numbers in mouse brain corpus callosum. Error bars indicate mean ± SEM. ns- nonsignificant; Student’s *t*-test, two-sided.

**Supplementary Figure S2: *Dicer1* is required for maintaining CNS resident microglia number**

**(A-H)** Representative immunostained confocal images of microglia (IBA1, A and B), myelin (PLP, C and D), OLs (APC, E and F), and OPCs (PDGFRα, G and H) from TMX-injected naïve Cre− control and Cre+ mutant animals. Scale bar - 100µm. **(I-L)** Bar graphs showing cell numbers (I, K and L) and myelin-occupied area (J) in naïve Tmem119^creERT2^ mouse brain CC. Error bars indicate mean ± SEM. \**p*< 0.05, ns- nonsignificant; Student’s *t*-test, two-sided.

**Supplementary Figure S3: *Dicer*-dependent miRNA expression is essential for regulating cell proliferation during the process of demyelination**

**(A)** Schematic illustration of Cx3cr1^creERT2^/Dicer^fl/fl^ animals undergoing TMX and BrdU treatment at the initiation of and 5 weeks into CUP feeding (6 weeks), respectively. **(B and C)** Representative immunostained confocal images of proliferating microglia (IBA1 and BrdU) from TMX-injected Cre− control and Cre+ mutant animals after six weeks of CUP feeding. Scale bar - 100µm.

**Supplementary Figure S4: Cuprizone feeding leads to death of mature OLs and enhanced microglial phagocytosis at the peak of demyelination**

**(A)** Schematic illustration of Cx3Cr1^creERT2^/Dicer^fl/fl^ animals undergoing TMX treatment and CUP feeding. **(B and C)** Representative immunostained confocal images of OL (OPCs+OLs) population (Olig2+) from TMX-injected Cre− control and Cre+ mutant animals after six weeks of CUP feeding. Scale bar - 100µm. **(D)** Bar graphs showing OL cell numbers in mouse brain CC. Error bars indicate mean ± SEM. ns- nonsignificant; Student’s *t*-test, two-sided. **(E-J)** Representative immunostained confocal images of OLs (APC, E and F), and apoptotic markers (CC3, G and H) from TMX-injected Cre− control and Cre+ mutant animals after six weeks of CUP feeding. Merged images are shown in panels I and J. Scale bar - 100µm. **(K)** Bar graphs showing apoptotic mature OLs in mouse brain corpus callosum. Error bars indicate mean ± SEM. \*\*\**p*< 0.001; Student’s *t*-test, two-sided. **(L-Q)** Representative immunostained confocal images of phagolysosome (CD68, L and M)-positive microglia (IBA1, N and O) from control C57 and animals fed with CUP for six weeks. Merged images are shown in panels P and Q. Scale bars - 50µm (A-F) and 20µm (G and H). **(R and S)** Bar graphs showing microglia-(R) and phagolysosome-occupied area (S) in control and CUP-fed mouse brain CC. Error bars indicate mean ± SEM. \**p*< 0.05, \*\**p*< 0.01; Student’s *t*-test, two-sided.

**Supplementary Figure S5: Adult microglia-specific *Dicer1* loss leads to decreased brain-specific resident microglia/macrophages**

**(A)** FACS sorting gating strategy to isolate CD45^int^/CD11b+ microglia from Cx3cr1^creERT2^ (Cre− and Cre+) animals. **(B)** Bar graphs showing CD45^int^/CD11b+ microglial cell numbers from naïve Cx3cr1^creERT2^ (Cre− and Cre+) animals treated with TMX only. Error bars indicate mean ± SEM. \**p*< 0.05; Student’s *t*-test, two-sided. **(C)** qPCR analysis of *mDicer1* expression in FASC-sorted CD45^int^/CD11b+ microglia from naïve Cx3cr1^creERT2^ (Cre− and Cre+) animals treated with TMX only. GAPDH was used as an endogenous control for qPCR analysis. Error bars indicate mean ± SEM. \*\**\*\*p*<0.0001; Student’s *t*- test, two-sided. **(D)** Representative cell-specific gene expression showing enrichment of microglia FACS samples over other CNS resident cells. **(E)** Bar graphs showing CD45^int^/CD11b+ microglia cell numbers from TMX-treated Cx3cr1^creERT2^ (Cre− and Cre+) animals fed with CUP diet for six weeks to induce demyelination. Error bars indicate mean ± SEM. Ns- nonsignificant; Student’s *t*-test, two-sided.

**Supplementary Figure S6: Evaluating the effects of *Dicer1* loss in adult microglia undergoing CUP feeding and remyelination**

**(A)** Schematic illustration of Cx3cr1^creERT2^/Dicer^fl/fl^ animals undergoing TMX treatment, CUP feeding, and remyelination. **(B and C)** Representative immunostained confocal images of OL (OPCs+OLs) population (Olig2+) from TMX-injected Cre− control and Cre+ mutant animals after six weeks of remyelination. Scale bar - 100µm. **(D)** Bar graphs showing OL cell numbers in mouse brain corpus callosum. Error bars indicate mean ± SEM. ns- nonsignificant; Student’s *t*-test, two-sided. **(E)** Bar graphs showing CD45^int^/CD11b+ flow-sorted cells from remyelinated Cre− and Cre+ mouse brain. Error bars indicate mean ± SEM. \**p*< 0.05; Student’s *t*-test, two-sided. **(F)** Volcano plot showing significant differentially-expressed genes (DEGs, log2Fold change >1 and *p*>0.01) in Dicer-ablated adult microglia FACS-sorted from remyelinated animals. Red dots represent upregulated transcripts and blue dots represent downregulated transcripts. **(G)** The top five biological processes associated with significant DEGs of Dicer-ablated FACS-sorted microglia from remyelinated animals.

**Supplementary Figure S7: Adult microglia lacking *Dicer1* expression during remyelination show delayed/failed remyelination**

**(A)** Schematic illustration of Cx3cr1^creERT2^/Dicer^fl/fl^ animals undergoing CUP feeding, TMX treatment, and remyelination. **(B and C)** Representative immunostained confocal images of mature OL population (APC) from TMX-injected Cre− control and Cre+ mutant animals after six weeks of remyelination. Scale bar - 100µm. **(D)** Volcano plot showing significant differentially-expressed genes (DEGs, log2Fold change >1 and *p*>0.01) in *Dicer*-ablated adult microglia FACS-sorted from remyelinated animals. **(E)** Bar graphs showing microglial homeostatic transcripts in mouse brain CC. Error bars indicate mean ± SEM. \**p*< 0.05 and \*\**p*< 0.01; Student’s *t*-test, two-sided. Red dots represent upregulated transcripts and blue dots represent downregulated transcripts. **(F)** The top five biological processes associated with significant DEGs of Dicer-ablated FACS-sorted microglia from remyelinated animals. **(G)** Heatmap showing significantly altered genes associated to regulation of attachment to spindle microtubule kinetochore (GO:005198) in remyelinated Cre− and Cre+ mutant animals. The scale indicates expression relative to CPM values across RNA samples.

## Supplementary Files

**Supplementary File 1:** List of differentially-expressed genes (DEGs) from flow-sorted microglia isolated from naïve Cx3cr1^creERT2^ Cre− control and Cre+ mutant animals.

**Supplementary File 2:** List of enriched biological processes of DEGs from flow-sorted microglia isolated from naïve Cx3cr1^creERT2^ Cre− control and Cre+ mutant animals.

**Supplementary File 3:** Ingenuity Canonical Pathways (IPA) analysis of DEGs from flow-sorted microglia isolated from naïve Cx3cr1^creERT2^ Cre− control and Cre+ mutant animals.

**Supplementary File 4:** List of differentially-expressed genes (DEGs) from flow-sorted microglia isolated from Cx3cr1^creERT2^ Cre− control and Cre+ mutant animals fed with CUP diet for 6 weeks.

**Supplementary File 5:** List of enriched biological processes of DEGs from flow-sorted microglia isolated from Cx3cr1^creERT2^ Cre− control and Cre+ mutant animals fed with CUP diet for 6 weeks.

**Supplementary File 6:** Ingenuity Canonical Pathways (IPA) analysis of DEGs from flow-sorted microglia isolated from Cx3cr1^creERT2^ Cre− control and Cre+ mutant animals fed with CUP diet for 6 weeks.

**Supplementary File 7:** List of differentially-expressed genes (DEGs) from flow-sorted microglia isolated from Cx3cr1^creERT2^ Cre− control and Cre+ mutant animals undergoing remyelination after six weeks of CUP feeding.

**Supplementary File 8:** List of enriched biological processes of DEGs from flow-sorted microglia isolated from Cx3cr1^creERT2^ Cre− control and Cre+ mutant animals undergoing remyelination after six weeks of CUP feeding.

**Supplementary File 9:** List of differentially-expressed genes (DEGs) from flow-sorted microglia isolated from Cx3cr1^creERT2^ Cre− control and Cre+ mutant animals treated with TMX at the peak of demyelination and undergoing remyelination after six weeks of CUP feeding.

**Supplementary File 10:** List of enriched biological processes of DEGs from flow-sorted microglia isolated from Cx3cr1^creERT2^ Cre− control and Cre+ mutant animals treated with TMX at the peak of demyelination and undergoing remyelination after six weeks of CUP feeding.

