## Supplementary figures and images for "Dicer deficiency in microglia leads to accelerated demyelination and failed remyelination"

### Supplementary Figure 1

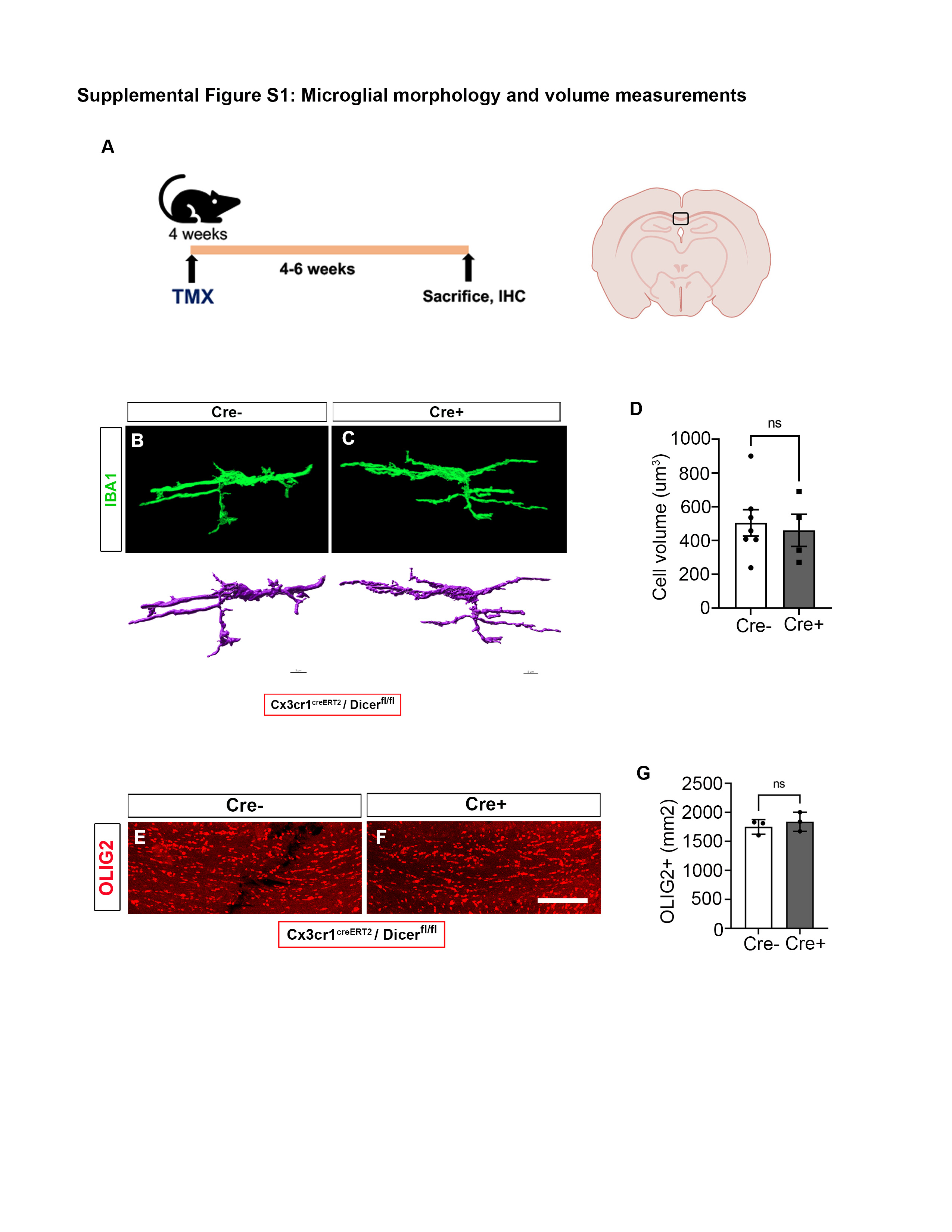

### Supplementary Figure 2

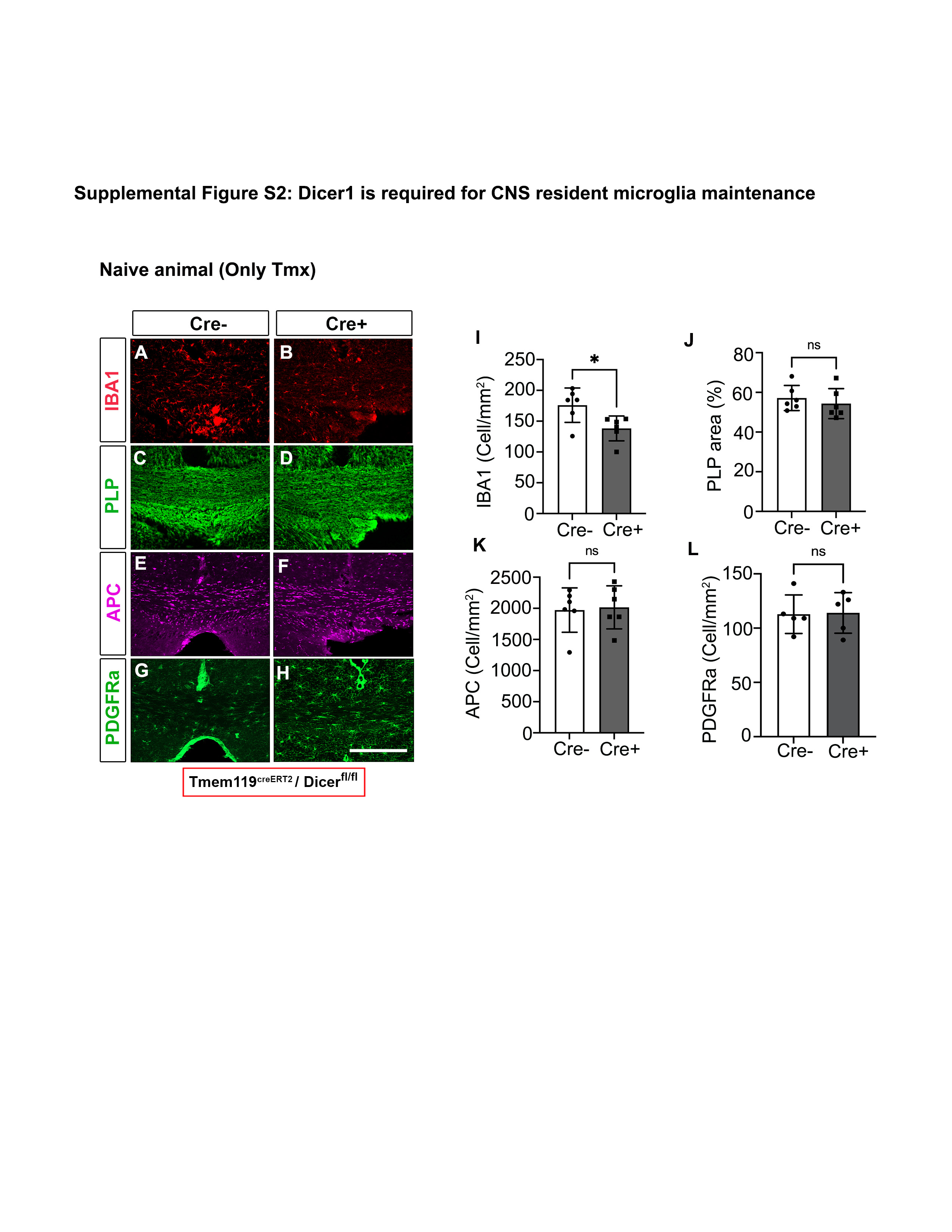

### Supplementary Figure 3

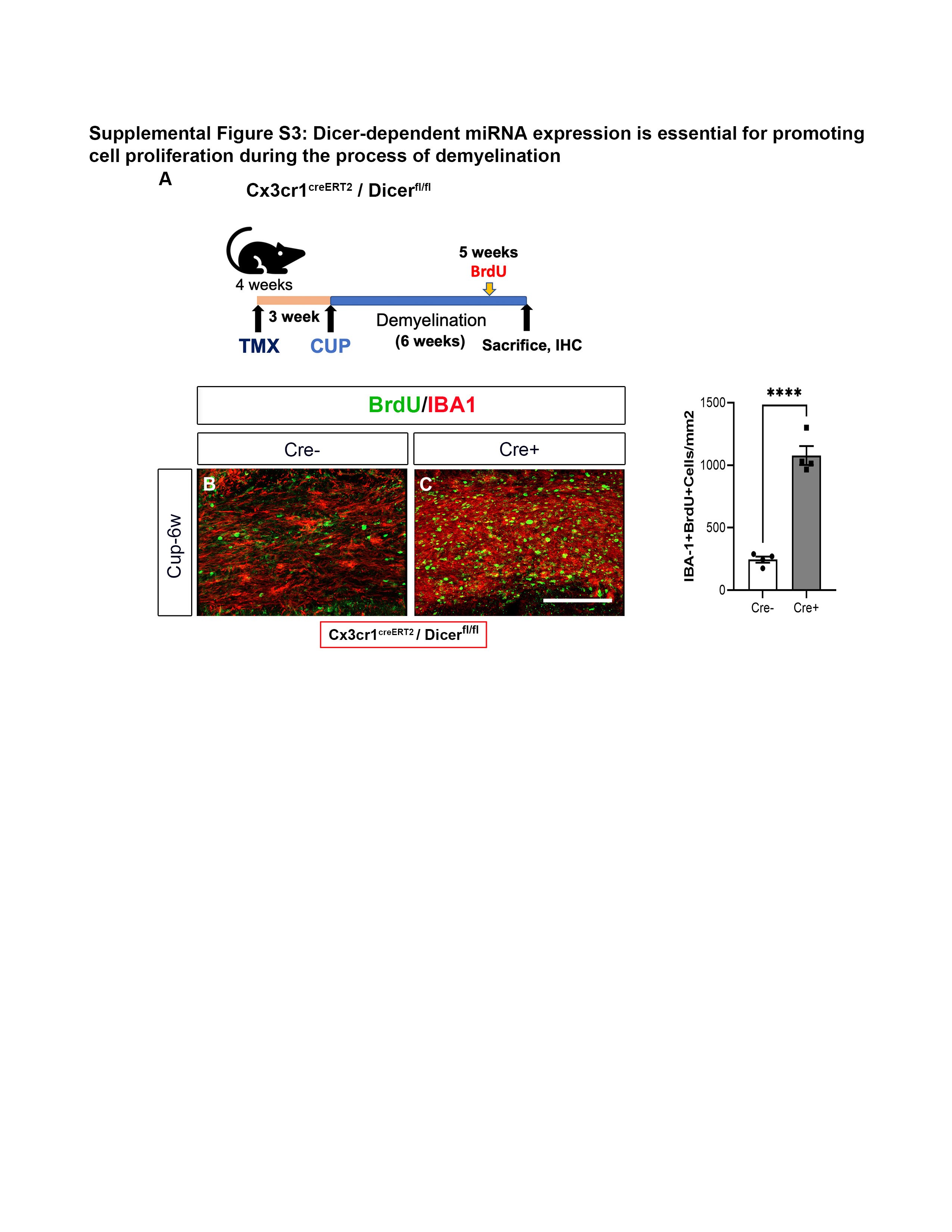

### Supplementary Figure 4

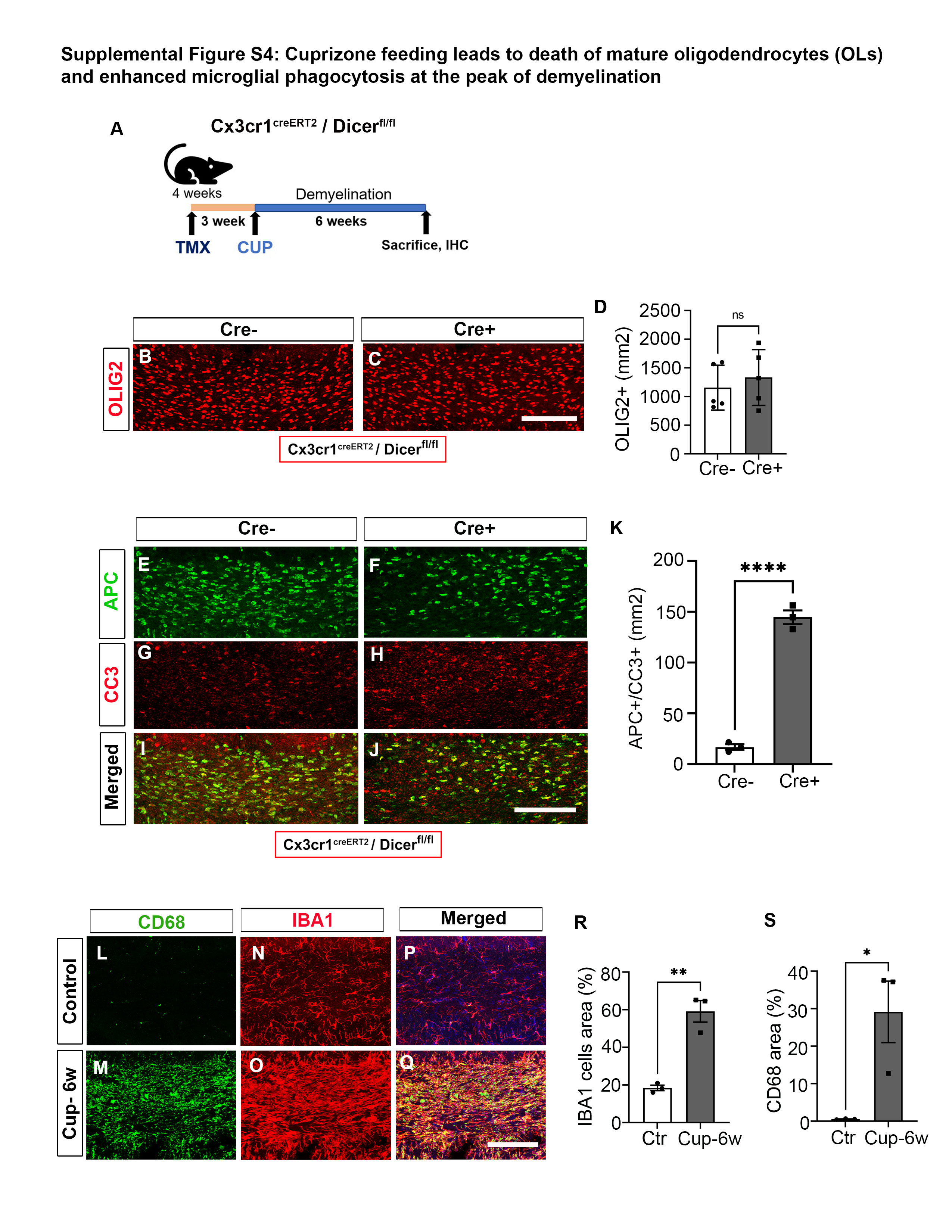

### Supplementary Figure 5

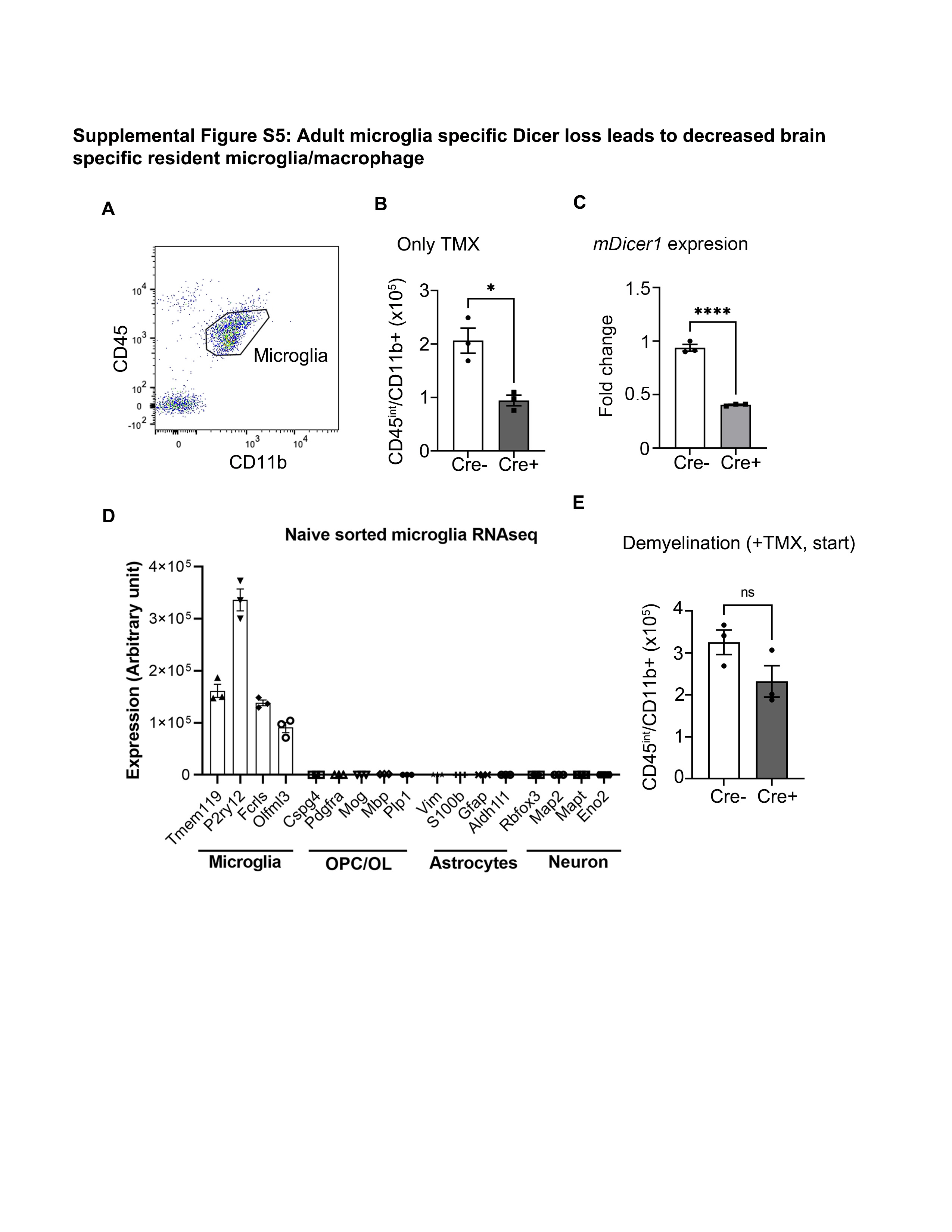

### Supplementary Figure 6

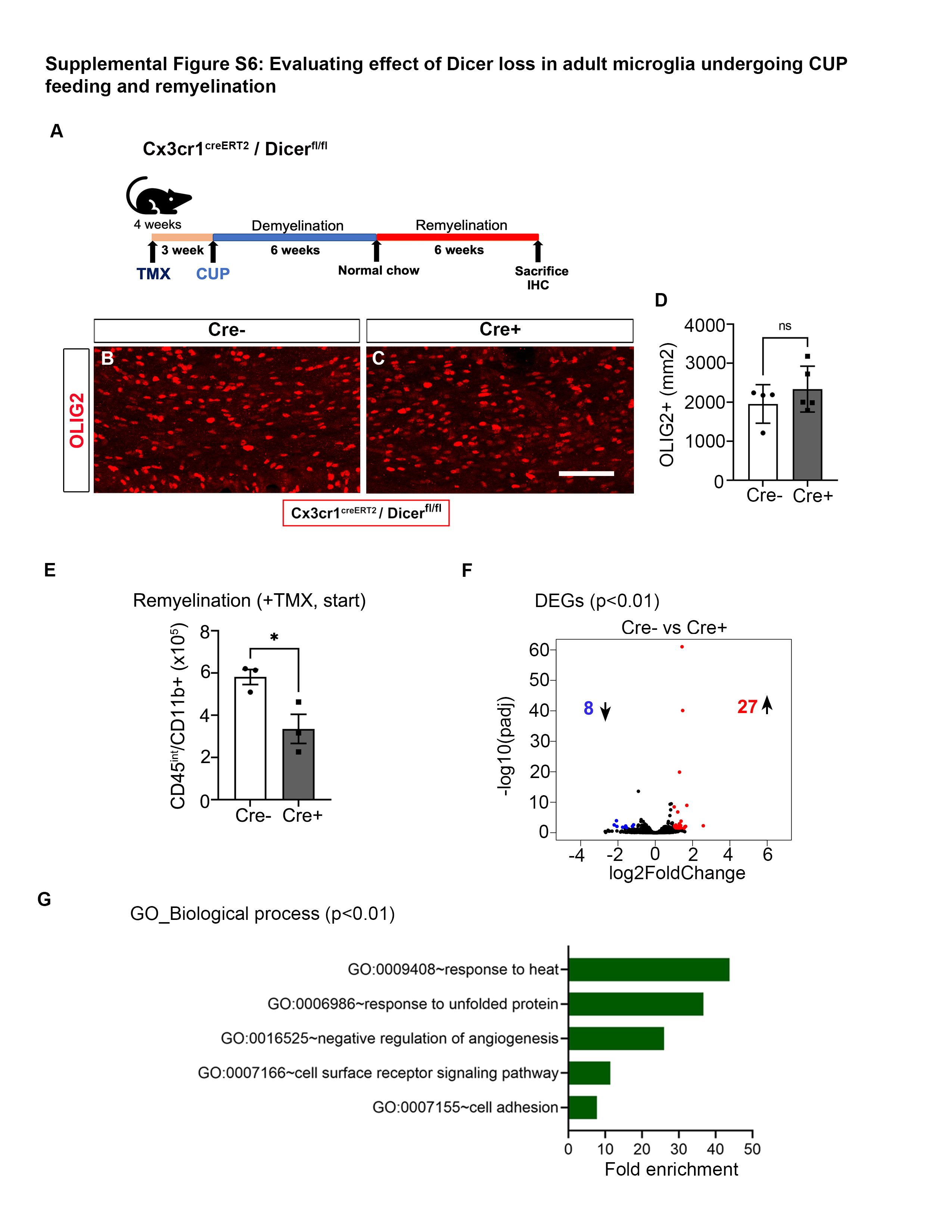

### Supplementary Figure 7

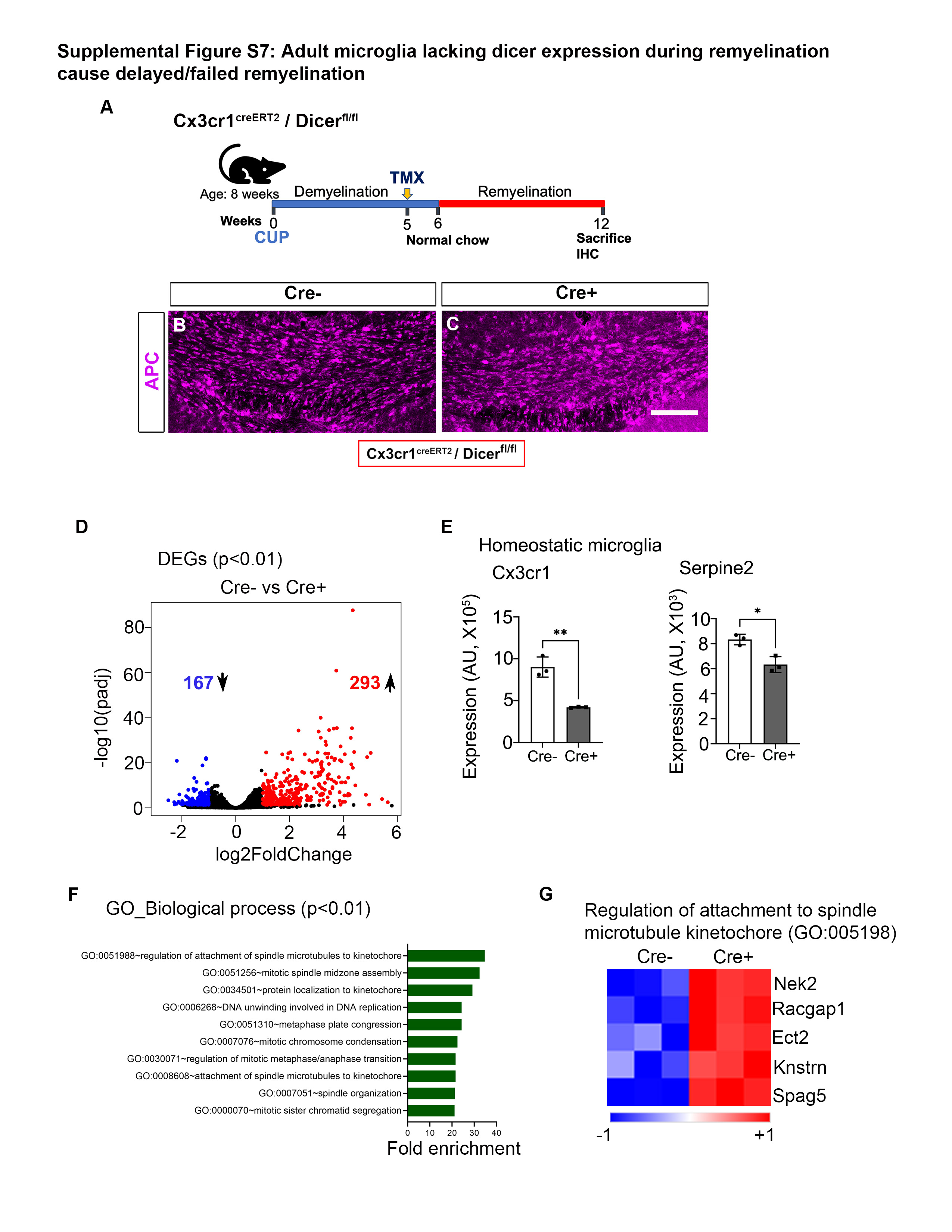
