## Supplementary File 10 for "Dicer deficiency in microglia leads to accelerated demyelination and failed remyelination"

**Supplementary file 10:** List of enriched biological processed of DEGs from flow sorted microglia isolated from Cx3cr1$^{creERT2}$ Cre- control and Cre+ mutant animals treated with TMX at peak of demyelination and undergoing remyelination after six weeks of CUP feeding.

| Category | Term | Count | % | PValue | Genes | List Total | Pop Hits | Pop Total | Fold Enrichment | Bonferroni | Benjamini | FDR |
|---|---|---|---|---|---|---|---|---|---|---|---|---|
| GOTERM_I | GO:0007049~cell cycle | 102 | 23.18182 | 3.47E-63 | 240641, 14 | 372 | 614 | 18082 | 8.074866029 | 6.11E-60 | 6.11E-60 | 5.97E-60 |
| GOTERM_I | GO:0051301~cell division | 75 | 17.04545 | 1.23E-51 | 240641, 14 | 372 | 374 | 18082 | 9.747498706 | 2.17E-48 | 1.09E-48 | 1.06E-48 |
| GOTERM_I | GO:0007067~mitotic nuclear division | 62 | 14.09091 | 2.28E-45 | 240641, 14 | 372 | 277 | 18082 | 10.87966306 | 4.02E-42 | 1.34E-42 | 1.31E-42 |
| GOTERM_I | GO:0007059~chromosome segregation | 23 | 5.227273 | 4.29E-18 | 66442, 719 | 372 | 89 | 18082 | 12.56149571 | 7.55E-15 | 1.89E-15 | 1.85E-15 |
| GOTERM_I | GO:0006974~cellular response to DNA damage stimulus | 36 | 8.181818 | 2.02E-12 | 18973, 710 | 372 | 420 | 18082 | 4.166359447 | 3.56E-09 | 7.12E-10 | 6.96E-10 |
| GOTERM_I | GO:0006260~DNA replication | 20 | 4.545455 | 7.47E-12 | 23834, 201 | 372 | 123 | 18082 | 7.903662908 | 1.31E-08 | 2.19E-09 | 2.14E-09 |
| GOTERM_I | GO:0000281~mitotic cytokinesis | 11 | 2.5 | 2.44E-10 | 16571, 193 | 372 | 30 | 18082 | 17.82275986 | 4.30E-07 | 6.14E-08 | 6.01E-08 |
| GOTERM_I | GO:0000070~mitotic sister chromatid segregation | 10 | 2.272727 | 3.71E-10 | 12704, 180 | 372 | 23 | 18082 | 21.13370734 | 6.53E-07 | 8.16E-08 | 7.98E-08 |
| GOTERM_I | GO:0006270~DNA replication initiation | 10 | 2.272727 | 5.83E-10 | 23834, 700 | 372 | 24 | 18082 | 20.2531362 | 1.03E-06 | 1.14E-07 | 1.11E-07 |
| GOTERM_I | GO:0006281~DNA repair | 27 | 6.136364 | 2.35E-09 | 18973, 193 | 372 | 318 | 18082 | 4.127054169 | 4.14E-06 | 4.14E-07 | 4.05E-07 |
| GOTERM_I | GO:0007080~mitotic metaphase plate congression | 10 | 2.272727 | 1.95E-08 | 56742, 229 | 372 | 34 | 18082 | 14.29633144 | 3.43E-05 | 3.12E-06 | 3.05E-06 |
| GOTERM_I | GO:0007052~mitotic spindle organization | 9 | 2.045455 | 6.32E-08 | 66442, 761 | 372 | 28 | 18082 | 15.62384793 | 1.11E-04 | 9.28E-06 | 9.07E-06 |
| GOTERM_I | GO:0007051~spindle organization | 7 | 1.590909 | 4.82E-07 | 51791, 165 | 372 | 16 | 18082 | 21.26579301 | 8.48E-04 | 6.13E-05 | 5.99E-05 |
| GOTERM_I | GO:0000082~G1/S transition of mitotic cell cycle | 11 | 2.5 | 4.88E-07 | 15904, 189 | 372 | 62 | 18082 | 8.62391606 | 8.58E-04 | 6.13E-05 | 5.99E-05 |
| GOTERM_I | GO:0007018~microtubule-based movement | 12 | 2.727273 | 5.32E-07 | 240641, 22 | 372 | 78 | 18082 | 7.478081059 | 9.36E-04 | 6.24E-05 | 6.10E-05 |
| GOTERM_I | GO:0000910~cytokinesis | 9 | 2.045455 | 6.68E-07 | 12704, 233 | 372 | 37 | 18082 | 11.82345248 | 0.001175821 | 7.35E-05 | 7.19E-05 |
| GOTERM_I | GO:0034501~protein localization to kinetochore | 6 | 1.363636 | 8.20E-07 | 76464, 703 | 372 | 10 | 18082 | 29.16451613 | 0.001441474 | 8.49E-05 | 8.30E-05 |
| GOTERM_I | GO:0051310~metaphase plate congression | 6 | 1.363636 | 2.49E-06 | 229841, 71 | 372 | 12 | 18082 | 24.30376344 | 0.004372707 | 2.43E-04 | 2.38E-04 |
| GOTERM_I | GO:0032467~positive regulation of cytokinesis | 8 | 1.818182 | 3.90E-06 | 12704, 240 | 372 | 33 | 18082 | 11.78364288 | 0.006842271 | 3.50E-04 | 3.42E-04 |
| GOTERM_I | GO:0007076~mitotic chromosome condensation | 6 | 1.363636 | 3.98E-06 | 14211, 700 | 372 | 13 | 18082 | 22.43424318 | 0.006977007 | 3.50E-04 | 3.42E-04 |
| GOTERM_I | GO:0051988~regulation of attachment of spindle microtubules to kinetochore | 5 | 1.136364 | 5.81E-06 | 18005, 136 | 372 | 7 | 18082 | 34.71966206 | 0.010177729 | 4.87E-04 | 4.76E-04 |
| GOTERM_I | GO:0030261~chromosome condensation | 6 | 1.363636 | 1.28E-05 | 21973, 760 | 372 | 16 | 18082 | 18.22782258 | 0.022331614 | 0.00102657 | 0.001004 |
| GOTERM_I | GO:0051726~regulation of cell cycle | 12 | 2.727273 | 1.93E-05 | 21335, 124 | 372 | 112 | 18082 | 5.207949309 | 0.0333572 | 0.00147504 | 0.001442 |
| GOTERM_I | GO:0006268~DNA unwinding involved in DNA replication | 5 | 1.136364 | 3.32E-05 | 21973, 172 | 372 | 10 | 18082 | 24.30376344 | 0.056770177 | 0.00243518 | 0.002381 |
| GOTERM_I | GO:0032147~activation of protein kinase activity | 7 | 1.590909 | 4.97E-05 | 72119, 136 | 372 | 33 | 18082 | 10.31068752 | 0.08369274 | 0.00336159 | 0.003287 |
| GOTERM_I | GO:0000086~G2/M transition of mitotic cell cycle | 7 | 1.590909 | 4.97E-05 | 12704, 125 | 372 | 33 | 18082 | 10.31068752 | 0.08369274 | 0.00336159 | 0.003287 |
| GOTERM_I | GO:0006468~protein phosphorylation | 27 | 6.136364 | 1.55E-04 | 11481, 114 | 372 | 576 | 18082 | 2.278477823 | 0.239087809 | 0.00993243 | 0.009712 |
| GOTERM_I | GO:0010212~response to ionizing radiation | 8 | 1.818182 | 1.58E-04 | 77011, 116 | 372 | 57 | 18082 | 6.822109036 | 0.243072428 | 0.00993243 | 0.009712 |
| GOTERM_I | GO:0051256~mitotic spindle midzone assembly | 4 | 0.909091 | 1.64E-04 | 16571, 718 | 372 | 6 | 18082 | 32.40501792 | 0.250286517 | 0.00993243 | 0.009712 |
| GOTERM_I | GO:0006334~nucleosome assembly | 10 | 2.272727 | 2.89E-04 | 66929, 126 | 372 | 104 | 18082 | 4.673800662 | 0.399091905 | 0.01692097 | 0.016546 |
| GOTERM_I | GO:0006302~double-strand break repair | 8 | 1.818182 | 2.98E-04 | 71988, 116 | 372 | 63 | 18082 | 6.172384366 | 0.408224722 | 0.01692097 | 0.016546 |
| GOTERM_I | GO:0006310~DNA recombination | 9 | 2.045455 | 3.54E-04 | 208084, 16 | 372 | 85 | 18082 | 5.146679317 | 0.463901657 | 0.01947898 | 0.019047 |
| GOTERM_I | GO:0016310~phosphorylation | 27 | 6.136364 | 3.98E-04 | 11481, 114 | 372 | 612 | 18082 | 2.144449715 | 0.503335817 | 0.02120309 | 0.020733 |
| GOTERM_I | GO:0000724~double-strand break repair via homologous recombination | 8 | 1.818182 | 4.78E-04 | 623474, 19 | 372 | 68 | 18082 | 5.718532574 | 0.569282633 | 0.0247677 | 0.024219 |
| GOTERM_I | GO:0008283~cell proliferation | 14 | 3.181818 | 6.44E-04 | 71878, 213 | 372 | 220 | 18082 | 3.093206256 | 0.678187973 | 0.03122756 | 0.030536 |
| GOTERM_I | GO:0008608~attachment of spindle microtubules to kinetochore | 4 | 0.909091 | 6.56E-04 | 229841, 76 | 372 | 9 | 18082 | 21.60334528 | 0.685194139 | 0.03122756 | 0.030536 |
| GOTERM_I | GO:0030071~regulation of mitotic metaphase/anaphase transition | 4 | 0.909091 | 6.56E-04 | 23834, 229 | 372 | 9 | 18082 | 21.60334528 | 0.685194139 | 0.03122756 | 0.030536 |
| GOTERM_I | GO:0090307~mitotic spindle assembly | 6 | 1.363636 | 6.92E-04 | 72119, 180 | 372 | 35 | 18082 | 8.332718894 | 0.704445139 | 0.03206523 | 0.031355 |
