## Supplementary File 1 for "Dicer deficiency in microglia leads to accelerated demyelination and failed remyelination"

**Supplementary file 1: List of differentially expressed gene (DEGs, log2FoldChange <-1/>+1, pAdj<0.05) from flow microglia isolated from naïve Cx3cr1creERT2 Cre- control and Cre+ mutant animals.**

| Gene.name | log2FoldC | pvalue | padj | Gene.name | log2FoldChange | pvalue | padj |
|---|---|---|---|---|---|---|---|
| Lgi2 | 6.22021 | 1.01E-17 | 1.93E-15 | Ifit3b | -1.000619474 | 2.20E-05 | 0.000243085 |
| Gm37199 | 6.00187 | 7.96E-06 | 0.000101 | Anxa4 | -1.001868714 | 1.95E-14 | 1.91E-12 |
| Gm8091 | 4.9342 | 1.98E-31 | 1.90E-28 | Ppa1 | -1.001914706 | 8.92E-08 | 2.01E-06 |
| Cebpb | 3.89623 | 0.000112 | 0.000976 | Lrrc8b | -1.002486035 | 0.0073915 | 0.030848988 |
| Tmc5 | 3.78309 | 1.44E-07 | 3.04E-06 | Lca5 | -1.002981829 | 0.0053092 | 0.023785386 |
| Skint3 | 3.6406 | 3.07E-29 | 2.13E-26 | Impdh1 | -1.003557227 | 0.0033564 | 0.016335999 |
| Dyrk3 | 3.61138 | 1.82E-06 | 2.82E-05 | Cxcl14 | -1.004567445 | 0.0044102 | 0.020433052 |
| Chst2 | 3.53575 | 5.72E-33 | 5.84E-30 | Slmap | -1.005099717 | 1.47E-13 | 1.20E-11 |
| Hist1h2bf | 3.50441 | 2.17E-05 | 0.00024 | Snhg8 | -1.006925856 | 0.0011129 | 0.006632535 |
| Alb | 3.45705 | 3.40E-57 | 1.04E-53 | Timp4 | -1.009615757 | 0.0001829 | 0.001487029 |
| Myt1l | 3.30426 | 9.15E-06 | 0.000114 | 4930515G01F | -1.010079192 | 0.0030411 | 0.01506509 |
| Retnlg | 3.26626 | 1.71E-06 | 2.67E-05 | Gm38843 | -1.011766206 | 2.40E-08 | 6.39E-07 |
| Mmp8 | 3.15473 | 8.52E-06 | 0.000107 | Ltb | -1.012616477 | 2.03E-05 | 0.00022724 |
| Hyal6 | 3.14297 | 4.75E-24 | 1.69E-21 | Gng7 | -1.012925415 | 0.0003453 | 0.002518855 |
| Spam1 | 3.09007 | 7.25E-05 | 0.000668 | Tmem221 | -1.013111431 | 2.03E-07 | 4.17E-06 |
| Gck | 3.06235 | 8.88E-22 | 2.72E-19 | Zfp248 | -1.013512002 | 7.78E-05 | 0.000709141 |
| Fam83d | 3.04116 | 3.84E-05 | 0.000392 | Mgst3 | -1.013744974 | 2.35E-07 | 4.71E-06 |
| Eid2 | 2.98578 | 3.88E-10 | 1.54E-08 | Stard8 | -1.014175778 | 1.35E-08 | 3.83E-07 |
| Gm8696 | 2.94441 | 5.27E-29 | 3.45E-26 | Tnfrsf13b | -1.015212387 | 1.26E-14 | 1.33E-12 |
| Il1r2 | 2.92688 | 3.48E-09 | 1.12E-07 | Capn2 | -1.015931029 | 4.80E-05 | 0.000474634 |
| 5330426L24Ril | 2.91824 | 1.14E-13 | 9.49E-12 | Zfp558 | -1.017042248 | 4.23E-06 | 5.86E-05 |
| Gm22267 | 2.88258 | 8.95E-08 | 2.01E-06 | Fam171b | -1.017409967 | 0.0006611 | 0.004292205 |
| Klri1 | 2.86042 | 4.95E-07 | 8.92E-06 | Gm13212 | -1.018424708 | 0.0011181 | 0.006650802 |
| 9130214F15Ril | 2.82678 | 0.000958 | 0.005874 | 4933433G15F | -1.019079648 | 0.0132578 | 0.048873544 |
| Gm5431 | 2.81392 | 6.89E-71 | 5.28E-67 | Tuft1 | -1.019482292 | 7.87E-06 | 0.000100424 |
| Mrap2 | 2.80202 | 3.37E-10 | 1.36E-08 | Psma8 | -1.019935832 | 3.51E-05 | 0.000361738 |
| Otud1 | 2.79364 | 1.02E-06 | 1.69E-05 | Rsad2 | -1.020857937 | 0.0006949 | 0.004488874 |
| Slc6a20b | 2.76309 | 0.001744 | 0.009643 | Trpv4 | -1.022653992 | 0.0002542 | 0.001973339 |
| Coro2a | 2.74838 | 3.69E-60 | 1.41E-56 | Ighj3 | -1.025603078 | 0.0073659 | 0.030758659 |
| Kif14 | 2.73003 | 0.001891 | 0.010294 | Gm39469 | -1.02683312 | 4.31E-07 | 7.93E-06 |
| Hspa1b | 2.7297 | 1.89E-83 | 2.89E-79 | Ccdc114 | -1.026981681 | 0.0083796 | 0.03419035 |
| Tesk1 | 2.71385 | 4.98E-17 | 7.94E-15 | Cldn10 | -1.027453443 | 0.0018363 | 0.010052535 |
| Gm4316 | 2.70277 | 0.001848 | 0.010103 | Agpat4 | -1.028472942 | 2.68E-05 | 0.0002879 |
| Hspa1a | 2.69208 | 2.45E-64 | 1.25E-60 | Gm15991 | -1.030319126 | 0.0022075 | 0.011706594 |
| Gm44135 | 2.67779 | 0.00026 | 0.002006 | Tmem44 | -1.03179878 | 0.0123372 | 0.046384044 |
| Mxd3 | 2.66738 | 0.000137 | 0.001167 | Chchd6 | -1.034049311 | 4.58E-07 | 8.37E-06 |
| Kif18b | 2.62786 | 0.004221 | 0.019685 | Ube2l6 | -1.035717893 | 8.93E-12 | 4.97E-10 |
| Adora1 | 2.52196 | 1.59E-15 | 2.02E-13 | Pigz | -1.036009535 | 0.0001824 | 0.001486366 |
| Nptxr | 2.51025 | 5.52E-10 | 2.10E-08 | Svip | -1.036855032 | 2.43E-07 | 4.85E-06 |
| Gm38317 | 2.49913 | 1.91E-06 | 2.95E-05 | Gstm4 | -1.038473325 | 4.18E-07 | 7.78E-06 |
| Gm48483 | 2.49427 | 0.003032 | 0.015037 | Lyplal1 | -1.041931278 | 9.45E-06 | 0.000117031 |
| 4930479D17Ri | 2.48688 | 1.76E-14 | 1.73E-12 | Irak1bp1 | -1.042725887 | 0.0090588 | 0.036332407 |
| Hyal5 | 2.46499 | 9.53E-15 | 1.03E-12 | Hilpda | -1.043127604 | 3.66E-09 | 1.17E-07 |

| | | | | | | | |
|---|---|---|---|---|---|---|---|
| Ano8 | 2.46062 | 0.003152 | 0.015539 | Angptl6 | -1.044488797 | 0.0005719 | 0.003808639 |
| Gm44620 | 2.45371 | 4.55E-05 | 0.000454 | Ift43 | -1.044793084 | 0.002835 | 0.014246558 |
| L3mbtl1 | 2.44401 | 0.001117 | 0.006648 | Zfp354c | -1.047858091 | 2.21E-05 | 0.00024344 |
| E2f7 | 2.38728 | 0.000738 | 0.004709 | Rida | -1.048154396 | 9.80E-10 | 3.55E-08 |
| Depdc1b | 2.38447 | 0.000444 | 0.003101 | Tor3a | -1.048793395 | 2.35E-12 | 1.49E-10 |
| Gm43414 | 2.38375 | 0.000115 | 0.001003 | Rwdd3 | -1.048908975 | 4.20E-06 | 5.83E-05 |
| Gata2 | 2.37352 | 2.07E-07 | 4.24E-06 | Tes | -1.04977957 | 1.11E-12 | 7.51E-11 |
| Mogs | 2.36646 | 2.15E-05 | 0.000238 | Gstm1 | -1.049815853 | 4.08E-07 | 7.63E-06 |
| C1qtnf7 | 2.33622 | 0.002361 | 0.012357 | Zfp37 | -1.050011309 | 1.14E-08 | 3.27E-07 |
| 1700020N01Ri | 2.32422 | 8.10E-31 | 6.90E-28 | 1700006E09R | -1.050935098 | 0.0048521 | 0.022112185 |
| Itgb5 | 2.31406 | 6.10E-05 | 0.000579 | Rtl8c | -1.054758536 | 4.41E-06 | 6.05E-05 |
| Cdc25c | 2.30821 | 0.001369 | 0.007897 | Glrx | -1.056452707 | 4.03E-08 | 1.01E-06 |
| Dnaaf5 | 2.3023 | 3.47E-12 | 2.14E-10 | Sat2 | -1.059487827 | 5.15E-09 | 1.58E-07 |
| Aspm | 2.29247 | 2.61E-05 | 0.000282 | Gm8113 | -1.059671702 | 0.0002803 | 0.002140276 |
| Ankle1 | 2.29082 | 0.003898 | 0.018449 | Rnu1a1 | -1.059851846 | 0.0057802 | 0.02533181 |
| Efcab11 | 2.28938 | 0.00351 | 0.01697 | Tmem237 | -1.062358843 | 1.82E-11 | 9.40E-10 |
| Klrd1 | 2.28641 | 8.87E-07 | 1.50E-05 | Ighj4 | -1.065607115 | 0.0005813 | 0.003864179 |
| Neil3 | 2.2658 | 0.000158 | 0.00132 | Prodh | -1.066309085 | 2.10E-07 | 4.30E-06 |
| Fbln1 | 2.25952 | 3.11E-19 | 7.55E-17 | Mrc1 | -1.066618016 | 1.41E-08 | 3.97E-07 |
| Adra2a | 2.23975 | 3.04E-05 | 0.00032 | Slc39a8 | -1.067022839 | 3.59E-06 | 5.09E-05 |
| P3h2 | 2.23549 | 5.12E-07 | 9.20E-06 | AB124611 | -1.068077372 | 0.0051406 | 0.02319931 |
| Acvr2b | 2.22076 | 3.63E-22 | 1.16E-19 | Hist1h1e | -1.068793398 | 0.0001784 | 0.001459194 |
| Asb4 | 2.21281 | 3.78E-11 | 1.85E-09 | Aph1b | -1.070991521 | 0.000403 | 0.002867345 |
| Hcar2 | 2.21139 | 6.33E-08 | 1.48E-06 | Arhgap18 | -1.071071987 | 7.85E-11 | 3.72E-09 |
| Lsm14b | 2.21084 | 1.31E-07 | 2.79E-06 | Tbc1d4 | -1.07279846 | 3.01E-06 | 4.39E-05 |
| Gm17110 | 2.19548 | 0.009452 | 0.037505 | Rrad | -1.073969975 | 1.29E-13 | 1.06E-11 |
| Crebzf | 2.19373 | 4.29E-07 | 7.91E-06 | Rab34 | -1.075092347 | 1.23E-10 | 5.54E-09 |
| Sapcd2 | 2.18327 | 0.001516 | 0.008602 | Tmem159 | -1.076978483 | 2.73E-14 | 2.60E-12 |
| Gm26714 | 2.17296 | 4.93E-13 | 3.58E-11 | Mfge8 | -1.078321737 | 1.10E-12 | 7.47E-11 |
| Plekha3 | 2.16707 | 2.55E-15 | 3.07E-13 | Atp10b | -1.079472825 | 0.0047951 | 0.021898199 |
| Spc24 | 2.16281 | 9.06E-06 | 0.000113 | Blvrb | -1.080433956 | 4.21E-12 | 2.50E-10 |
| Knl1 | 2.15059 | 2.71E-06 | 4.01E-05 | Rnasel | -1.083555382 | 9.50E-11 | 4.39E-09 |
| Syn2 | 2.14651 | 0.008737 | 0.03531 | Kbtbd12 | -1.084621053 | 0.0001534 | 0.001287072 |
| Gm29781 | 2.14259 | 0.005505 | 0.024459 | Batf3 | -1.085681514 | 8.40E-10 | 3.07E-08 |
| Hyal4 | 2.13798 | 1.87E-12 | 1.21E-10 | 1810021B22R | -1.087179917 | 1.44E-07 | 3.05E-06 |
| Ska3 | 2.13322 | 9.39E-06 | 0.000116 | Aph1c | -1.088782769 | 0.0001208 | 0.00104091 |
| 2610306O10Ri | 2.12662 | 0.010417 | 0.040477 | Rragd | -1.089642097 | 0.0006 | 0.003971578 |
| Sox12 | 2.10733 | 0.013095 | 0.0484 | P3h4 | -1.091161833 | 4.22E-06 | 5.85E-05 |
| Foxm1 | 2.09772 | 1.79E-08 | 4.87E-07 | Zfp993 | -1.091686462 | 0.0011695 | 0.006908502 |
| Hist3h2ba | 2.09344 | 2.01E-05 | 0.000226 | Map2k3os | -1.093514239 | 0.0081411 | 0.033404057 |
| Foxr1 | 2.09257 | 1.64E-10 | 7.20E-09 | Lgals1 | -1.094001824 | 0.0005642 | 0.003765393 |
| Csnk1g2 | 2.07959 | 0.001456 | 0.008316 | Vegfa | -1.094888318 | 0.0024953 | 0.012887439 |
| Fzd8 | 2.0665 | 4.09E-05 | 0.000416 | Cxcl16 | -1.095376444 | 2.99E-12 | 1.87E-10 |
| Ckap4 | 2.06642 | 0.000117 | 0.001014 | Myo15 | -1.096648326 | 0.0126993 | 0.047338978 |
| Crls1 | 2.06212 | 5.64E-10 | 2.12E-08 | Creb3l1 | -1.096981226 | 0.0013613 | 0.007859292 |
| Enpp6 | 2.06211 | 0.001732 | 0.009594 | Zfp992 | -1.097092747 | 2.33E-11 | 1.19E-09 |
| Gm12934 | 2.0587 | 0.003551 | 0.017134 | Clec7a | -1.100304188 | 1.62E-11 | 8.61E-10 |

| | | | | | | | |
|---|---|---|---|---|---|---|---|
| Gm14314 | 2.04131 | 0.002467 | 0.012775 | Lbh | -1.100795736 | 0.0004564 | 0.003170033 |
| Plac8 | 2.03326 | 1.51E-05 | 0.000176 | Cd81 | -1.102991796 | 2.24E-25 | 8.38E-23 |
| Fbxo45 | 2.01921 | 6.13E-15 | 6.91E-13 | Ripk3 | -1.103571579 | 2.02E-06 | 3.10E-05 |
| 2510039O18Ri | 2.00527 | 0.001144 | 0.006783 | Ptgr1 | -1.103915812 | 1.41E-06 | 2.24E-05 |
| Rnf149 | 1.99157 | 6.53E-07 | 1.14E-05 | Akap17b | -1.104709028 | 0.0097371 | 0.038409233 |
| Trim43c | 1.99001 | 0.003314 | 0.016181 | Srl | -1.104984528 | 5.35E-07 | 9.55E-06 |
| Ptchd1 | 1.97676 | 1.78E-22 | 5.79E-20 | Fam43a | -1.108287054 | 0.0004507 | 0.003139323 |
| Tatdn2 | 1.96872 | 1.66E-09 | 5.75E-08 | Gadd45b | -1.108987335 | 4.85E-06 | 6.59E-05 |
| Prkcq | 1.96297 | 6.34E-31 | 5.72E-28 | Sdf2l1 | -1.110977373 | 4.46E-17 | 7.34E-15 |
| Gm38832 | 1.95141 | 4.91E-17 | 7.92E-15 | Fblim1 | -1.112593391 | 2.56E-09 | 8.51E-08 |
| Lockd | 1.944 | 0.001445 | 0.008263 | Il12rb2 | -1.112947845 | 0.0007195 | 0.004612734 |
| Ccdc71l | 1.94082 | 1.37E-12 | 9.09E-11 | Gale | -1.114031501 | 5.60E-05 | 0.000540021 |
| Dvl1 | 1.93647 | 4.39E-13 | 3.26E-11 | Maoa | -1.114343337 | 0.0001552 | 0.001300432 |
| Gm28151 | 1.93142 | 2.36E-08 | 6.33E-07 | Prnp | -1.11554969 | 1.01E-07 | 2.24E-06 |
| E2f1 | 1.92838 | 1.95E-12 | 1.25E-10 | Cox6a2 | -1.115557613 | 0.0008444 | 0.00527933 |
| Prc1 | 1.92725 | 0.007784 | 0.032239 | 2900060B14F | -1.115618504 | 0.0001675 | 0.001383904 |
| Hnrnpd | 1.92538 | 0.000347 | 0.002526 | Rnu2-10 | -1.120513475 | 8.14E-05 | 0.000737867 |
| Mark3 | 1.92426 | 0.000138 | 0.00117 | Zfp72 | -1.121065871 | 0.0001837 | 0.001492447 |
| Gm23722 | 1.92102 | 0.011095 | 0.042538 | Lst1 | -1.121272989 | 2.88E-09 | 9.48E-08 |
| Tbc1d5 | 1.91803 | 2.34E-52 | 5.98E-49 | Sardh | -1.122132358 | 9.39E-07 | 1.57E-05 |
| Gm8817 | 1.91438 | 8.88E-14 | 7.68E-12 | Thbs2 | -1.122257866 | 0.0011163 | 0.006645295 |
| Ube2c | 1.91046 | 9.18E-06 | 0.000114 | Tnf | -1.122969134 | 5.07E-08 | 1.22E-06 |
| Basp1 | 1.90635 | 0.000272 | 0.002083 | Gm29395 | -1.125327631 | 7.41E-05 | 0.000679826 |
| Phf19 | 1.90543 | 0.002427 | 0.012615 | Gm37352 | -1.126600742 | 1.71E-07 | 3.58E-06 |
| Dock9 | 1.90172 | 3.60E-43 | 6.12E-40 | Ephx1 | -1.126931242 | 2.85E-14 | 2.69E-12 |
| Lemd2 | 1.88415 | 6.75E-05 | 0.000628 | Bend5 | -1.127120117 | 8.78E-08 | 1.98E-06 |
| Kifc1 | 1.88381 | 0.003763 | 0.017961 | Slamf8 | -1.127818148 | 1.77E-11 | 9.20E-10 |
| Kif11 | 1.88023 | 0.010146 | 0.039605 | E130317F20R | -1.128259977 | 2.39E-06 | 3.59E-05 |
| Dapk3 | 1.87464 | 0.001347 | 0.007791 | Gm128 | -1.131576259 | 0.0001291 | 0.001104339 |
| Mtch1 | 1.87025 | 0.010565 | 0.040926 | Lyzl4 | -1.132882055 | 7.02E-05 | 0.000649019 |
| Fam8a1 | 1.86073 | 2.14E-25 | 8.18E-23 | Sox15 | -1.134956713 | 0.0047459 | 0.021751166 |
| Rab12 | 1.86041 | 0.000393 | 0.002811 | Gnpda1 | -1.135255759 | 8.19E-15 | 8.89E-13 |
| Siglec5 | 1.851 | 1.74E-39 | 2.42E-36 | Slc16a9 | -1.13580644 | 7.00E-05 | 0.00064794 |
| Mki67 | 1.84662 | 0.009143 | 0.036612 | Lrrc75aos2 | -1.136250414 | 0.0126614 | 0.047220598 |
| Prkar2a | 1.84281 | 1.17E-21 | 3.51E-19 | Zkscan7 | -1.138049213 | 0.0042574 | 0.019826898 |
| Gm2065 | 1.83956 | 4.22E-05 | 0.000426 | Aldh2 | -1.139823869 | 2.33E-10 | 9.86E-09 |
| Gas2 | 1.83522 | 0.000122 | 0.00105 | Nat1 | -1.140121048 | 0.0038715 | 0.018358908 |
| Hlf | 1.8315 | 1.02E-17 | 1.93E-15 | Adamts1 | -1.142049197 | 7.06E-08 | 1.64E-06 |
| Cecr2 | 1.82554 | 1.05E-07 | 2.31E-06 | Adam33 | -1.145609209 | 0.0094467 | 0.037505036 |
| Ccna2 | 1.82355 | 0.007323 | 0.030633 | Atp6v0e2 | -1.147273699 | 0.0043055 | 0.020022388 |
| Nek2 | 1.82353 | 0.000316 | 0.00235 | Fam26f | -1.147804854 | 2.91E-08 | 7.55E-07 |
| Frat2 | 1.82176 | 2.70E-06 | 4.00E-05 | Rell1 | -1.148495853 | 7.61E-05 | 0.000695274 |
| Zdhhc18 | 1.82174 | 3.10E-13 | 2.36E-11 | Gm23849 | -1.148602942 | 1.04E-09 | 3.75E-08 |
| Gm15706 | 1.82169 | 7.27E-06 | 9.36E-05 | A930007I19R | -1.150956832 | 3.83E-07 | 7.23E-06 |
| Gm2396 | 1.81893 | 2.97E-09 | 9.75E-08 | Smim10l2a | -1.151385475 | 1.38E-10 | 6.18E-09 |
| Gm43254 | 1.81482 | 5.37E-05 | 0.000523 | Gm26887 | -1.152741337 | 0.0004069 | 0.00288795 |
| Map3k9 | 1.81126 | 2.02E-15 | 2.49E-13 | Kank2 | -1.153614856 | 0.0085556 | 0.034744132 |

| | | | | | | | |
|---|---|---|---|---|---|---|---|
| Esr1 | 1.81027 | 8.44E-22 | 2.64E-19 | Guca1a | -1.156882492 | 6.75E-05 | 0.000628398 |
| Psrc1 | 1.80741 | 0.011106 | 0.042558 | Ly9 | -1.158237311 | 3.64E-13 | 2.74E-11 |
| Arf6 | 1.8023 | 0.001428 | 0.008179 | D930028M14 | -1.158694417 | 0.0064809 | 0.027736178 |
| Chad | 1.78957 | 1.38E-18 | 3.10E-16 | C4b | -1.161671345 | 0.0019936 | 0.010757156 |
| Troap | 1.78535 | 0.000921 | 0.005694 | Rab4a | -1.162521193 | 1.35E-05 | 0.000158875 |
| 9530046B11Ri | 1.78449 | 0.009323 | 0.037138 | Slc7a4 | -1.16298448 | 0.0022484 | 0.01188754 |
| Grk6 | 1.78299 | 0.000273 | 0.002093 | Padi2 | -1.163640469 | 6.83E-18 | 1.36E-15 |
| Gm24225 | 1.78038 | 1.26E-16 | 1.89E-14 | 4933421O10I | -1.16672334 | 6.04E-06 | 7.95E-05 |
| Ccne1 | 1.77814 | 0.002641 | 0.013465 | Dctd | -1.167681289 | 0.0002698 | 0.002070729 |
| Fbxo33 | 1.77664 | 3.57E-15 | 4.24E-13 | Lyz2 | -1.171306849 | 6.09E-14 | 5.42E-12 |
| Gm19582 | 1.77428 | 0.00653 | 0.027898 | Prelid2 | -1.175831765 | 3.08E-09 | 1.01E-07 |
| Cpeb3 | 1.77199 | 2.87E-11 | 1.43E-09 | Opn3 | -1.178463664 | 0.0007036 | 0.004526299 |
| Gm47205 | 1.77185 | 0.000142 | 0.001205 | H2-K2 | -1.179608783 | 2.62E-10 | 1.09E-08 |
| Ckap2 | 1.76893 | 2.43E-05 | 0.000265 | Trappc6a | -1.180210918 | 2.24E-10 | 9.59E-09 |
| Prim2 | 1.76663 | 4.29E-36 | 5.05E-33 | Gm43144 | -1.181403324 | 0.0029546 | 0.014717145 |
| Tmed7 | 1.76543 | 0.002044 | 0.010967 | Ifitm3 | -1.184384337 | 2.58E-07 | 5.11E-06 |
| Phf21b | 1.76065 | 4.29E-05 | 0.000431 | Lypd6 | -1.185961173 | 5.42E-06 | 7.23E-05 |
| Oip5 | 1.7604 | 0.004836 | 0.022044 | Lrrc8c | -1.186230686 | 0.0024833 | 0.012846143 |
| Mmp9 | 1.75406 | 0.001014 | 0.006132 | Hdhd3 | -1.195223782 | 0.0034226 | 0.016616324 |
| Hist1h1c | 1.75402 | 2.03E-19 | 5.11E-17 | Pyroxd2 | -1.195482451 | 6.50E-09 | 1.96E-07 |
| Esco2 | 1.75327 | 0.000261 | 0.002013 | Crip1 | -1.195746842 | 6.27E-05 | 0.000591981 |
| Rnpepl1 | 1.75035 | 8.56E-13 | 5.94E-11 | Ropn1l | -1.197794548 | 0.0067844 | 0.028730062 |
| Gm13071 | 1.74966 | 0.008236 | 0.033732 | Gm45191 | -1.198436063 | 4.88E-05 | 0.000482056 |
| Grsf1 | 1.749 | 0.000996 | 0.006052 | Ifi207 | -1.200525109 | 0.0003282 | 0.002416135 |
| Ube2m | 1.74872 | 0.001005 | 0.006095 | Emp3 | -1.20176088 | 3.91E-14 | 3.58E-12 |
| Smap1 | 1.74138 | 3.50E-10 | 1.40E-08 | C230037L18R | -1.202999649 | 0.0009916 | 0.006031942 |
| Pimreg | 1.73865 | 0.000258 | 0.001999 | Adcyap1r1 | -1.203331307 | 0.0003517 | 0.002555493 |
| F830045P16Ril | 1.7325 | 4.34E-05 | 0.000435 | Mterf1b | -1.206488744 | 0.0002874 | 0.002180508 |
| Cdyl | 1.73087 | 1.74E-12 | 1.14E-10 | Rab3d | -1.208416606 | 2.36E-05 | 0.000257683 |
| Crtap | 1.72857 | 2.37E-15 | 2.91E-13 | Fuom | -1.208665376 | 0.002628 | 0.013422225 |
| Agap3 | 1.72712 | 4.57E-10 | 1.79E-08 | D630045J12R | -1.209642918 | 2.03E-06 | 3.12E-05 |
| Cds1 | 1.71973 | 1.50E-25 | 5.89E-23 | Armcx2 | -1.209967727 | 2.53E-08 | 6.73E-07 |
| Itgb3 | 1.714 | 5.40E-29 | 3.45E-26 | Cers4 | -1.214819815 | 1.50E-05 | 0.000175546 |
| Pbk | 1.71102 | 0.008614 | 0.034932 | Rnf17 | -1.216394729 | 3.75E-05 | 0.000384583 |
| Ppp1r12c | 1.71027 | 2.95E-05 | 0.000312 | Gtf2h3 | -1.217229037 | 1.89E-09 | 6.50E-08 |
| Mocs3 | 1.70961 | 3.24E-05 | 0.000339 | Arhgap29 | -1.217546883 | 2.75E-09 | 9.07E-08 |
| Heg1 | 1.70379 | 1.74E-05 | 0.000199 | 4933431E20R | -1.221687859 | 0.0038766 | 0.01837756 |
| Cldn12 | 1.70094 | 7.68E-20 | 1.99E-17 | Mx1 | -1.221737277 | 4.42E-09 | 1.38E-07 |
| Anln | 1.69776 | 6.96E-05 | 0.000645 | Fkbp14 | -1.222794484 | 1.07E-07 | 2.34E-06 |
| Fam91a1 | 1.69768 | 3.30E-19 | 7.71E-17 | Stk26 | -1.222952602 | 4.54E-07 | 8.31E-06 |
| Kif22 | 1.69707 | 1.52E-05 | 0.000178 | E030042O20I | -1.224927136 | 0.0002887 | 0.002183551 |
| Prkaca | 1.69563 | 0.001994 | 0.010757 | Zfp647 | -1.227071999 | 0.0005635 | 0.003762154 |
| Spen | 1.69019 | 1.38E-12 | 9.14E-11 | Tmem38a | -1.227135871 | 1.03E-05 | 0.000126205 |
| Whamm | 1.69002 | 5.16E-07 | 9.25E-06 | Gm44789 | -1.22944674 | 0.0052491 | 0.023584948 |
| B230311B06Ri | 1.68636 | 5.62E-08 | 1.34E-06 | 1700123M08 | -1.230053 | 0.0080614 | 0.033157088 |
| Inpp5a | 1.68623 | 6.19E-08 | 1.45E-06 | Ftl2 | -1.230993477 | 0.001703 | 0.009460046 |
| Irs3 | 1.68357 | 4.36E-08 | 1.07E-06 | Acot6 | -1.234156438 | 0.0123362 | 0.046384044 |

| | | | | | | | |
|---|---|---|---|---|---|---|---|
| Map3k19 | 1.6824 | 4.35E-13 | 3.25E-11 | Thnsl2 | -1.234677597 | 6.96E-08 | 1.62E-06 |
| Cpeb2 | 1.67993 | 1.87E-28 | 1.10E-25 | Slc39a11 | -1.234683154 | 3.44E-13 | 2.61E-11 |
| 4930579C12Ri | 1.67538 | 0.000648 | 0.004223 | Bcar3 | -1.235100856 | 0.0062684 | 0.027015425 |
| Stil | 1.67158 | 0.004149 | 0.019424 | Apbb1 | -1.237730598 | 7.07E-08 | 1.64E-06 |
| Gm7854 | 1.66989 | 0.006453 | 0.027639 | Igsf10 | -1.238837823 | 0.0009907 | 0.006031313 |
| Ncoa1 | 1.66698 | 9.13E-13 | 6.27E-11 | Eif4e3 | -1.239563744 | 7.50E-13 | 5.29E-11 |
| Vps54 | 1.66606 | 3.27E-19 | 7.71E-17 | Speg | -1.244015131 | 2.88E-15 | 3.45E-13 |
| Nid2 | 1.66091 | 2.82E-17 | 5.09E-15 | Gm4961 | -1.244282859 | 0.0132514 | 0.048873388 |
| E2f8 | 1.65745 | 0.007358 | 0.030734 | Gnb5 | -1.244354564 | 8.86E-08 | 2.00E-06 |
| Prr11 | 1.65612 | 0.006196 | 0.02681 | Mxra8 | -1.24573626 | 0.0002229 | 0.001770249 |
| Gm35339 | 1.65465 | 0.002937 | 0.014649 | Tnni3 | -1.245816263 | 6.83E-05 | 0.000634468 |
| Clspn | 1.6545 | 0.009558 | 0.03786 | R74862 | -1.246572291 | 7.76E-08 | 1.77E-06 |
| Efcab6 | 1.65205 | 0.000377 | 0.002709 | Gm11638 | -1.248068091 | 0.0016931 | 0.00941654 |
| Cdca2 | 1.65128 | 0.000315 | 0.002336 | Arl6 | -1.251049311 | 1.60E-08 | 4.43E-07 |
| Dact3 | 1.64986 | 9.38E-07 | 1.57E-05 | Dse | -1.252367856 | 1.50E-16 | 2.19E-14 |
| Lrrc47 | 1.64906 | 0.004662 | 0.021427 | Gm38140 | -1.256641341 | 9.35E-05 | 0.000833952 |
| Tlr11 | 1.64336 | 7.30E-05 | 0.000671 | Dnase1l1 | -1.257517116 | 8.23E-10 | 3.02E-08 |
| Hmmr | 1.6432 | 0.000112 | 0.000974 | Gm44745 | -1.262454375 | 0.0001121 | 0.000977576 |
| Ccnb2 | 1.64241 | 2.41E-06 | 3.61E-05 | Fcrls | -1.26412259 | 2.73E-11 | 1.39E-09 |
| Mknk2 | 1.63859 | 1.45E-15 | 1.87E-13 | Pter | -1.26572903 | 8.11E-12 | 4.57E-10 |
| Kcnk12 | 1.63419 | 0.010065 | 0.039409 | Cdc42ep1 | -1.265948973 | 0.0030409 | 0.01506509 |
| Gm826 | 1.63159 | 4.54E-08 | 1.11E-06 | Ccdc80 | -1.267120091 | 0.0096042 | 0.037973106 |
| Abca13 | 1.63134 | 9.72E-14 | 8.37E-12 | Rab42 | -1.267423577 | 0.0006082 | 0.004012149 |
| Axin1 | 1.62968 | 2.01E-08 | 5.45E-07 | Gm3164 | -1.268267589 | 2.03E-06 | 3.12E-05 |
| Gm16894 | 1.62826 | 0.000215 | 0.001719 | Thap8 | -1.272751744 | 0.0010114 | 0.006120924 |
| Pi4k2a | 1.62628 | 0.004062 | 0.019101 | Stx1a | -1.272847643 | 2.86E-05 | 0.000304094 |
| Pwwp2a | 1.62507 | 2.85E-13 | 2.19E-11 | Hmgn3 | -1.274855787 | 1.56E-05 | 0.000181381 |
| Adamtsl2 | 1.62447 | 2.69E-18 | 5.88E-16 | Nipsnap1 | -1.276437574 | 8.63E-07 | 1.46E-05 |
| Mcm10 | 1.62445 | 0.003937 | 0.018603 | Gm37560 | -1.277539416 | 0.0066048 | 0.028156361 |
| Bcr | 1.6156 | 3.28E-08 | 8.43E-07 | Ifit1 | -1.282078857 | 3.21E-05 | 0.000336071 |
| Aurka | 1.61481 | 4.93E-06 | 6.68E-05 | Olr1 | -1.283765707 | 0.0004342 | 0.003049351 |
| Scarf2 | 1.60258 | 0.002849 | 0.014284 | Ccdc122 | -1.284259529 | 4.20E-09 | 1.32E-07 |
| Gm37909 | 1.60186 | 0.013066 | 0.048341 | Zswim7 | -1.287644202 | 5.46E-07 | 9.69E-06 |
| Kdm1a | 1.60186 | 0.000714 | 0.004583 | Myom1 | -1.288491587 | 3.10E-11 | 1.53E-09 |
| Dlgap5 | 1.59739 | 0.004104 | 0.019263 | 1700011I03Ri | -1.289100815 | 1.50E-05 | 0.000175989 |
| Gm20696 | 1.59606 | 1.34E-11 | 7.25E-10 | Dmtn | -1.29241673 | 1.87E-12 | 1.21E-10 |
| Kcnn4 | 1.59387 | 2.64E-16 | 3.74E-14 | Fbxo36 | -1.298130716 | 2.73E-05 | 0.000292567 |
| Usp25 | 1.59265 | 4.21E-05 | 0.000426 | Klrg2 | -1.298175897 | 0.0003249 | 0.002398016 |
| Ccnb1 | 1.58867 | 0.007872 | 0.032496 | Gm2000 | -1.299658868 | 0.0016059 | 0.009000472 |
| Kctd2 | 1.58866 | 0.002871 | 0.01438 | Stard6 | -1.30088126 | 0.0123916 | 0.046531377 |
| Khsrp | 1.58759 | 1.10E-11 | 5.97E-10 | 4430402I18Ri | -1.305220569 | 0.0010908 | 0.006526353 |
| Carm1 | 1.58284 | 5.13E-10 | 1.98E-08 | Ddah2 | -1.308431272 | 2.81E-11 | 1.41E-09 |
| Hsp90aa1 | 1.58057 | 3.48E-39 | 4.45E-36 | Mgmt | -1.309528691 | 1.15E-06 | 1.88E-05 |
| Rac1 | 1.58016 | 0.009341 | 0.037182 | Gm3667 | -1.31138665 | 1.36E-07 | 2.90E-06 |
| Ythdf1 | 1.56921 | 0.000296 | 0.002229 | Tceal9 | -1.311464433 | 9.97E-18 | 1.93E-15 |
| Gm37317 | 1.56661 | 0.00018 | 0.001468 | Plekhh2 | -1.312752857 | 0.0001151 | 0.001002117 |
| Ifi30 | 1.56542 | 1.26E-13 | 1.04E-11 | Tctn2 | -1.313415336 | 6.87E-05 | 0.000638147 |

| | | | | | | | |
|---|---|---|---|---|---|---|---|
| Cdk2ap1 | 1.56395 | 4.67E-08 | 1.14E-06 | Tlr1 | -1.315139797 | 1.67E-14 | 1.66E-12 |
| Arglu1 | 1.56177 | 0.000288 | 0.002181 | AA414768 | -1.317071973 | 0.0113181 | 0.043276603 |
| Klf2 | 1.55313 | 0.000207 | 0.001663 | Ankrd37 | -1.318148487 | 6.07E-05 | 0.000576585 |
| Aurkb | 1.55238 | 1.48E-05 | 0.000174 | Hist2h2ac | -1.318509606 | 0.0096514 | 0.038129977 |
| Samd4b | 1.55023 | 8.37E-12 | 4.70E-10 | 4930523C07F | -1.320383091 | 4.95E-05 | 0.000487506 |
| Sspo | 1.54756 | 0.010905 | 0.041926 | Fgl2 | -1.321420898 | 1.36E-12 | 9.09E-11 |
| Gtse1 | 1.54749 | 0.003023 | 0.015012 | Chst10 | -1.323994345 | 3.87E-06 | 5.43E-05 |
| 1810026B05Ri | 1.54574 | 8.46E-13 | 5.89E-11 | Fam213a | -1.32486473 | 1.86E-15 | 2.32E-13 |
| Sft2d3 | 1.54541 | 4.15E-07 | 7.75E-06 | Ighj2 | -1.327080772 | 5.63E-05 | 0.000541669 |
| C5ar2 | 1.54193 | 9.24E-41 | 1.42E-37 | S100a1 | -1.32917295 | 2.79E-11 | 1.40E-09 |
| Cpd | 1.54189 | 1.20E-15 | 1.56E-13 | Tnfsf13b | -1.32938996 | 2.34E-13 | 1.83E-11 |
| Gsk3a | 1.54123 | 1.04E-08 | 3.04E-07 | Ctla2b | -1.333914419 | 2.99E-14 | 2.79E-12 |
| Hecw2 | 1.53857 | 6.31E-05 | 0.000595 | S100a10 | -1.337578429 | 6.05E-07 | 1.06E-05 |
| Tbc1d10b | 1.53751 | 5.51E-14 | 4.99E-12 | Rpl39l | -1.342883712 | 0.0068117 | 0.028837509 |
| Gm43148 | 1.53353 | 0.001246 | 0.007289 | Entpd2 | -1.345457027 | 0.0066262 | 0.02820811 |
| Gm17586 | 1.53166 | 1.29E-06 | 2.08E-05 | H2-Ob | -1.347136382 | 4.06E-18 | 8.51E-16 |
| Mybph | 1.5316 | 3.88E-12 | 2.35E-10 | Tfpi | -1.352779505 | 3.00E-20 | 8.26E-18 |
| Plk1 | 1.53037 | 0.000154 | 0.001291 | Slc25a43 | -1.352962157 | 4.98E-05 | 0.000489768 |
| Ttyh2 | 1.52776 | 8.36E-13 | 5.84E-11 | Enpp1 | -1.354242834 | 8.08E-07 | 1.38E-05 |
| Abhd17b | 1.52769 | 8.38E-10 | 3.07E-08 | Ccdc102a | -1.356717899 | 4.24E-06 | 5.87E-05 |
| Slc35e1 | 1.52711 | 0.000604 | 0.003991 | Id2 | -1.357964235 | 8.93E-26 | 3.60E-23 |
| Map3k5 | 1.52272 | 1.22E-12 | 8.23E-11 | Pilra | -1.358172234 | 4.03E-17 | 6.79E-15 |
| Cdca8 | 1.52213 | 0.000777 | 0.00493 | Fndc4 | -1.358624861 | 9.66E-06 | 0.000119176 |
| Mcl1 | 1.51861 | 0.00222 | 0.011756 | Exoc3l4 | -1.359105951 | 0.0097151 | 0.038332547 |
| Tcf24 | 1.51666 | 0.002783 | 0.014024 | Cxcl10 | -1.359552188 | 1.22E-06 | 1.99E-05 |
| Rybp | 1.51524 | 7.60E-10 | 2.82E-08 | 9330104G04F | -1.359993498 | 3.62E-06 | 5.13E-05 |
| Otulin | 1.51157 | 0.011418 | 0.043558 | Slamf9 | -1.361473761 | 1.67E-27 | 8.52E-25 |
| Gm16323 | 1.5114 | 0.011944 | 0.045148 | Jam3 | -1.361856348 | 6.86E-09 | 2.06E-07 |
| Upk1b | 1.50926 | 8.52E-30 | 6.87E-27 | Selenbp1 | -1.362960434 | 1.53E-14 | 1.55E-12 |
| Dusp7 | 1.50538 | 0.00013 | 0.001108 | Ly6g6d | -1.363707946 | 2.10E-05 | 0.000233741 |
| Zfpm1 | 1.49984 | 1.21E-05 | 0.000145 | Ica1 | -1.363949112 | 0.0008774 | 0.005465142 |
| Slc30a1 | 1.49802 | 5.59E-06 | 7.41E-05 | Gm3716 | -1.367223442 | 4.29E-07 | 7.91E-06 |
| Entpd6 | 1.49755 | 1.72E-29 | 1.32E-26 | Lpl | -1.368393837 | 5.52E-15 | 6.27E-13 |
| Spock2 | 1.49672 | 0.011375 | 0.043459 | Gm16486 | -1.372055238 | 0.002189 | 0.011629636 |
| Clstn1 | 1.48942 | 0.000309 | 0.002302 | Paox | -1.373860192 | 1.06E-13 | 9.00E-12 |
| B3gnt3 | 1.48856 | 0.000279 | 0.002135 | Stard4 | -1.374866914 | 1.30E-05 | 0.000154445 |
| Gm38318 | 1.48753 | 0.003932 | 0.01859 | Tmem220 | -1.378131798 | 9.93E-06 | 0.00012208 |
| Pclaf | 1.48735 | 0.000304 | 0.002275 | Pstpip2 | -1.378276754 | 0.000472 | 0.003258057 |
| Hist2h3c2 | 1.48683 | 0.009562 | 0.037867 | Ly6g6e | -1.381841829 | 5.69E-05 | 0.000546889 |
| Zmynd19 | 1.48212 | 0.000413 | 0.002923 | Pmepa1os | -1.38290034 | 0.0026168 | 0.013373719 |
| Fancf | 1.48011 | 0.001305 | 0.007594 | Dnaaf3 | -1.384598542 | 2.68E-05 | 0.0002879 |
| Zfp651 | 1.47686 | 4.45E-05 | 0.000444 | B3gnt7 | -1.385243814 | 0.001298 | 0.007559411 |
| Mapkapk2 | 1.47482 | 0.002274 | 0.011986 | Dennd2d | -1.387749828 | 0.0017491 | 0.009665002 |
| Tsga10ip | 1.47471 | 0.007324 | 0.030633 | Fbxo17 | -1.388751669 | 1.09E-06 | 1.80E-05 |
| Mlycd | 1.47401 | 0.004306 | 0.020022 | Nfkbiz | -1.390321791 | 1.95E-13 | 1.56E-11 |
| Birc5 | 1.46226 | 0.000132 | 0.00113 | Dhtkd1 | -1.394173294 | 0.0004464 | 0.003118164 |
| Sipa1l1 | 1.45943 | 2.52E-27 | 1.24E-24 | Cers1 | -1.397737535 | 0.0024971 | 0.012891397 |

| | | | | | | | |
|---|---|---|---|---|---|---|---|
| Gm4524 | 1.45492 | 0.003989 | 0.018811 | Zfp819 | -1.400135634 | 0.0067039 | 0.02846796 |
| Unc5a | 1.45468 | 0.002949 | 0.014696 | Phf11b | -1.401163737 | 4.05E-18 | 8.51E-16 |
| Smad6 | 1.45122 | 3.14E-06 | 4.56E-05 | Pcp4 | -1.40631009 | 6.67E-06 | 8.69E-05 |
| Gm13479 | 1.4506 | 7.37E-24 | 2.57E-21 | Tmem154 | -1.406645432 | 0.000415 | 0.002936085 |
| Donson | 1.44963 | 1.47E-15 | 1.88E-13 | Myc | -1.408066318 | 2.59E-13 | 2.02E-11 |
| Hsf2bp | 1.444 | 5.37E-10 | 2.05E-08 | Aifm2 | -1.41228894 | 0.010222 | 0.039857362 |
| Znrf1 | 1.4427 | 2.35E-10 | 9.89E-09 | Gm7115 | -1.41354564 | 0.0101327 | 0.039570081 |
| Lrrc8d | 1.43962 | 0.001621 | 0.009065 | Nenf | -1.415185024 | 1.06E-14 | 1.14E-12 |
| Gm43821 | 1.43911 | 1.56E-10 | 6.90E-09 | Cd63-ps | -1.415525009 | 0.0004097 | 0.00290427 |
| Smarca5 | 1.43746 | 0.000183 | 0.001486 | Gadl1 | -1.41639039 | 6.65E-06 | 8.66E-05 |
| 4930516B21Ri | 1.43692 | 5.32E-08 | 1.27E-06 | Gm45768 | -1.417618334 | 0.0030496 | 0.015101924 |
| Aven | 1.43355 | 5.56E-06 | 7.38E-05 | Hspb11 | -1.423662147 | 7.92E-06 | 0.000100818 |
| Ube3c | 1.43272 | 1.89E-10 | 8.15E-09 | Abhd3 | -1.423996511 | 1.28E-12 | 8.60E-11 |
| Gm47283 | 1.4298 | 3.32E-08 | 8.50E-07 | Lrrc27 | -1.429631001 | 0.001765 | 0.009738528 |
| Ppp6r1 | 1.42859 | 5.55E-10 | 2.10E-08 | Slamf1 | -1.433679643 | 3.90E-12 | 2.35E-10 |
| Cytip | 1.42792 | 5.19E-10 | 1.99E-08 | Pdia5 | -1.438448099 | 6.20E-17 | 9.79E-15 |
| Fbxl17 | 1.42765 | 1.72E-07 | 3.60E-06 | Plk2 | -1.44062761 | 5.43E-07 | 9.65E-06 |
| Hist2h2be | 1.42749 | 4.49E-13 | 3.32E-11 | Nbl1 | -1.443995234 | 2.94E-06 | 4.31E-05 |
| Slc24a3 | 1.41992 | 1.21E-10 | 5.51E-09 | Enpp5 | -1.445913628 | 1.54E-14 | 1.55E-12 |
| Pde8a | 1.4182 | 1.03E-05 | 0.000126 | Il1rn | -1.446118443 | 0.0053625 | 0.023953708 |
| Col27a1 | 1.41751 | 9.64E-20 | 2.46E-17 | Anxa6 | -1.450723623 | 1.48E-26 | 6.65E-24 |
| Mfhas1 | 1.41747 | 2.51E-09 | 8.35E-08 | 2900026A02F | -1.457226965 | 3.18E-07 | 6.08E-06 |
| Sap30l | 1.41428 | 2.08E-05 | 0.000232 | Kcnn2 | -1.45816987 | 0.0125709 | 0.046974474 |
| Cdk13 | 1.41198 | 6.37E-08 | 1.49E-06 | Tnfsf9 | -1.464091288 | 0.0001122 | 0.000978464 |
| Cdhr1 | 1.40881 | 0.001826 | 0.010009 | Pfkfb3 | -1.469929082 | 4.03E-15 | 4.75E-13 |
| Tnks2 | 1.40662 | 2.74E-11 | 1.39E-09 | Gm14597 | -1.47062127 | 1.00E-05 | 0.000123071 |
| Pank1 | 1.40622 | 3.04E-05 | 0.000321 | Gm25313 | -1.472373901 | 0.0064201 | 0.027537739 |
| Gm15232 | 1.40453 | 3.98E-26 | 1.69E-23 | Gm25939 | -1.479668314 | 0.0001388 | 0.001178967 |
| Pde3b | 1.4038 | 3.87E-12 | 2.35E-10 | Sh3pxd2b | -1.480845096 | 0.0024404 | 0.012658418 |
| Colgalt1 | 1.40164 | 7.21E-14 | 6.35E-12 | Tmem204 | -1.481380547 | 3.22E-14 | 2.97E-12 |
| Adrb1 | 1.39873 | 1.15E-07 | 2.49E-06 | Pomgnt2 | -1.481560417 | 3.43E-10 | 1.37E-08 |
| Crybg3 | 1.39656 | 1.83E-05 | 0.000209 | Gm32591 | -1.481565305 | 1.49E-10 | 6.67E-09 |
| Arhgap33 | 1.3957 | 0.010703 | 0.041324 | Nim1k | -1.482756063 | 0.0032914 | 0.016112181 |
| Sipa1l2 | 1.38864 | 1.36E-14 | 1.41E-12 | 4930550C14F | -1.482831708 | 0.0021797 | 0.011592309 |
| Ctdp1 | 1.38823 | 0.010584 | 0.040968 | Cd63 | -1.491268508 | 1.03E-46 | 1.97E-43 |
| Nab1 | 1.38696 | 1.21E-07 | 2.61E-06 | Tmem205 | -1.492089335 | 1.95E-07 | 4.04E-06 |
| Ptger3 | 1.38013 | 3.81E-17 | 6.64E-15 | Gsto2 | -1.498643304 | 0.0010398 | 0.006260711 |
| Gm42648 | 1.37986 | 0.011018 | 0.042285 | 9-Sep | -1.50017629 | 1.03E-07 | 2.27E-06 |
| Igfbp4 | 1.37961 | 5.11E-28 | 2.90E-25 | Cd83 | -1.501580997 | 1.43E-27 | 7.54E-25 |
| Btbd2 | 1.37788 | 0.007823 | 0.032357 | Fam69b | -1.502377118 | 0.0005089 | 0.003478777 |
| Kctd6 | 1.37667 | 6.02E-06 | 7.94E-05 | Lipc | -1.504797517 | 0.004516 | 0.020866153 |
| Cd300c | 1.37451 | 0.000455 | 0.003162 | Ust | -1.505005959 | 4.15E-10 | 1.65E-08 |
| Gm44941 | 1.37382 | 0.004166 | 0.01948 | Plaur | -1.508234921 | 4.23E-16 | 5.79E-14 |
| Perp | 1.37362 | 4.15E-17 | 6.90E-15 | Cisd3 | -1.508799032 | 8.79E-10 | 3.20E-08 |
| Gm26984 | 1.3722 | 0.006293 | 0.027093 | Bex3 | -1.509786735 | 1.24E-09 | 4.44E-08 |
| Cables2 | 1.37115 | 0.002244 | 0.01187 | Gstt2 | -1.516452954 | 1.11E-10 | 5.07E-09 |
| Ppp2r2d | 1.37054 | 0.001929 | 0.010473 | Dlg3 | -1.518732601 | 1.59E-14 | 1.59E-12 |

| | | | | | | | |
|---|---|---|---|---|---|---|---|
| Hspb1 | 1.36877 | 0.000228 | 0.001803 | Smpd3 | -1.519458426 | 0.003153 | 0.015538845 |
| 2310008N11Ri | 1.36688 | 0.003579 | 0.017227 | Clec1a | -1.523039126 | 0.0002914 | 0.00220201 |
| Tmem150c | 1.36683 | 0.000403 | 0.002867 | Sdc4 | -1.523999177 | 3.29E-16 | 4.54E-14 |
| Lmnb1 | 1.36387 | 1.52E-06 | 2.41E-05 | Batf | -1.524240731 | 8.53E-12 | 4.77E-10 |
| Kcnh4 | 1.36132 | 0.000709 | 0.00455 | Ighj1 | -1.53267557 | 2.16E-10 | 9.26E-09 |
| Igsf8 | 1.35945 | 4.51E-17 | 7.34E-15 | Plin2 | -1.533297818 | 2.78E-26 | 1.22E-23 |
| Hist1h3d | 1.35621 | 0.003354 | 0.016329 | D830025C05F | -1.533356555 | 5.08E-05 | 0.000498614 |
| Ankrd40 | 1.35263 | 2.39E-09 | 8.02E-08 | Prdm5 | -1.534464902 | 0.0019734 | 0.010677255 |
| Ankrd13a | 1.35241 | 0.007026 | 0.029599 | Cpq | -1.53797465 | 2.98E-14 | 2.79E-12 |
| Nuf2 | 1.34859 | 0.001848 | 0.010103 | Ccdc8 | -1.557303035 | 0.0001115 | 0.000973863 |
| Slc44a1 | 1.34756 | 0.01038 | 0.040377 | Car9 | -1.558915543 | 5.61E-05 | 0.000540257 |
| Tmem243 | 1.34723 | 0.002033 | 0.010929 | Gem | -1.559841753 | 2.22E-11 | 1.14E-09 |
| Zswim5 | 1.34361 | 6.15E-08 | 1.45E-06 | Efhc1 | -1.565131077 | 0.0055337 | 0.02452552 |
| Adgra3 | 1.3432 | 1.35E-05 | 0.000159 | Col25a1 | -1.569229671 | 0.0009302 | 0.00573613 |
| Hs6st1 | 1.34178 | 5.43E-09 | 1.66E-07 | Ccr5 | -1.575500028 | 2.60E-16 | 3.74E-14 |
| Slc45a4 | 1.34166 | 5.90E-11 | 2.86E-09 | Gm4951 | -1.577132972 | 4.17E-05 | 0.000422088 |
| Gm26848 | 1.34015 | 0.012096 | 0.04569 | Meig1 | -1.57915183 | 0.0062313 | 0.026889459 |
| Rab21 | 1.33972 | 4.19E-08 | 1.04E-06 | Ptges | -1.579480081 | 0.0130842 | 0.048395282 |
| Tcim | 1.33942 | 0.004589 | 0.021154 | Gm24305 | -1.580206876 | 0.0023995 | 0.01251422 |
| Actr5 | 1.33918 | 0.007228 | 0.030317 | C030037D09F | -1.580447302 | 0.0010136 | 0.006131478 |
| Klhl21 | 1.3383 | 0.00762 | 0.031689 | Tceal8 | -1.582492591 | 3.95E-18 | 8.51E-16 |
| Map4k3 | 1.33813 | 6.87E-12 | 3.93E-10 | Selenom | -1.583012252 | 8.01E-07 | 1.37E-05 |
| Parp12 | 1.33676 | 0.003094 | 0.015286 | Sigirr | -1.583891118 | 0.0038936 | 0.018436256 |
| Cables1 | 1.33504 | 0.005455 | 0.024287 | Mdga1 | -1.584588072 | 0.0001678 | 0.001385829 |
| Dgkh | 1.33409 | 1.41E-05 | 0.000166 | Gstm5 | -1.58578119 | 7.12E-20 | 1.88E-17 |
| Fkbp5 | 1.33054 | 2.57E-17 | 4.68E-15 | Calr3 | -1.586981555 | 0.0004248 | 0.00299729 |
| Kynu | 1.32945 | 6.54E-05 | 0.000612 | Gm16287 | -1.589758367 | 0.000965 | 0.005900648 |
| Gm42502 | 1.32724 | 0.013353 | 0.049166 | Triqk | -1.59287779 | 1.20E-05 | 0.000144603 |
| Ppm1j | 1.32376 | 0.000268 | 0.002058 | Anxa9 | -1.595470283 | 0.0027005 | 0.01370548 |
| Acvr2a | 1.32336 | 1.14E-23 | 3.90E-21 | Nradd | -1.599119906 | 5.05E-14 | 4.60E-12 |
| Ncor2 | 1.32283 | 4.66E-07 | 8.47E-06 | Sugct | -1.600521681 | 0.0003665 | 0.002649277 |
| Btbd1 | 1.32227 | 8.27E-09 | 2.44E-07 | Gm23472 | -1.601492071 | 2.56E-06 | 3.81E-05 |
| Zdhhc14 | 1.32184 | 0.007348 | 0.0307 | P4htm | -1.60720489 | 0.0039296 | 0.018588488 |
| Gm11998 | 1.31872 | 1.59E-10 | 7.03E-09 | Gdpd2 | -1.607912677 | 0.0118744 | 0.044952938 |
| Gper1 | 1.31463 | 0.000682 | 0.004417 | Mettl27 | -1.608846015 | 5.78E-05 | 0.000553705 |
| NA | 1.31072 | 0.001941 | 0.010518 | Cib2 | -1.613043722 | 6.46E-15 | 7.17E-13 |
| Spred1 | 1.30965 | 2.88E-11 | 1.43E-09 | Efemp2 | -1.613626512 | 1.43E-09 | 5.06E-08 |
| Itga9 | 1.30867 | 1.64E-10 | 7.20E-09 | Gatm | -1.615039887 | 2.39E-17 | 4.40E-15 |
| Hist1h2be | 1.30672 | 2.49E-18 | 5.53E-16 | Ackr3 | -1.616295706 | 1.14E-05 | 0.000138386 |
| Ahdc1 | 1.3034 | 3.01E-08 | 7.79E-07 | Folh1 | -1.622132267 | 0.0131237 | 0.048484243 |
| Rhoq | 1.30222 | 4.60E-10 | 1.80E-08 | Angptl7 | -1.623182802 | 7.57E-12 | 4.30E-10 |
| Brip1 | 1.30221 | 0.002328 | 0.01222 | Palld | -1.626991164 | 0.0011655 | 0.006892529 |
| Reps1 | 1.30138 | 6.87E-08 | 1.60E-06 | Ccl5 | -1.631009627 | 0.0098398 | 0.038744632 |
| Arih1 | 1.30012 | 3.01E-11 | 1.49E-09 | Sfmbt2 | -1.631568595 | 3.45E-06 | 4.93E-05 |
| Lonrf1 | 1.29837 | 4.51E-09 | 1.41E-07 | Mmp11 | -1.634259518 | 2.99E-10 | 1.22E-08 |
| Heatr1 | 1.29736 | 5.24E-23 | 1.75E-20 | Cela1 | -1.637237212 | 0.0005184 | 0.003513254 |
| 6030400A10Ri | 1.29672 | 2.86E-07 | 5.56E-06 | Ccl4 | -1.641187269 | 4.30E-26 | 1.78E-23 |

| | | | | | | | |
|---|---|---|---|---|---|---|---|
| Ogfrl1 | 1.29583 | 3.01E-12 | 1.87E-10 | Pygl | -1.641840299 | 1.65E-11 | 8.70E-10 |
| Senp2 | 1.29583 | 4.12E-20 | 1.11E-17 | Wscd1 | -1.645028571 | 8.28E-14 | 7.25E-12 |
| Gm16193 | 1.29186 | 3.65E-06 | 5.16E-05 | Lag3 | -1.645161032 | 8.08E-16 | 1.08E-13 |
| Pkmyt1 | 1.29164 | 4.07E-05 | 0.000414 | Arhgef25 | -1.646691009 | 5.35E-10 | 2.05E-08 |
| Plpp6 | 1.29093 | 0.004924 | 0.022408 | C1ra | -1.648309934 | 3.37E-05 | 0.000348844 |
| Cenpa | 1.28955 | 0.000101 | 0.000893 | 1810034E14R | -1.649753377 | 0.0010091 | 0.006114117 |
| Hist1h4i | 1.28753 | 1.08E-05 | 0.000131 | Gpr4 | -1.653948779 | 0.0023443 | 0.012288597 |
| Csnk1g3 | 1.28698 | 0.003244 | 0.015917 | Zfp454 | -1.655496681 | 0.0123559 | 0.046443156 |
| Nckap5l | 1.28526 | 1.49E-16 | 2.19E-14 | Ahrr | -1.663796691 | 4.04E-05 | 0.000410146 |
| Gm6498 | 1.28508 | 0.005464 | 0.02432 | Gm23971 | -1.667034268 | 0.0004483 | 0.003125531 |
| Cmya5 | 1.28241 | 1.15E-17 | 2.15E-15 | Eno4 | -1.675357448 | 8.49E-05 | 0.000765994 |
| Mis18bp1 | 1.27997 | 0.005167 | 0.023303 | Slc35e4 | -1.678956409 | 7.08E-11 | 3.39E-09 |
| Otub2 | 1.27639 | 8.43E-09 | 2.48E-07 | Ccl3 | -1.683601699 | 1.21E-24 | 4.41E-22 |
| Itpr3 | 1.26992 | 1.26E-09 | 4.51E-08 | Haao | -1.686608643 | 8.77E-11 | 4.11E-09 |
| Mpp6 | 1.26921 | 5.05E-12 | 2.97E-10 | Gm26616 | -1.687800022 | 9.77E-06 | 0.000120342 |
| Sav1 | 1.26895 | 0.000674 | 0.004373 | Klhl13 | -1.707472274 | 3.51E-09 | 1.13E-07 |
| Zfp746 | 1.26821 | 0.003293 | 0.016112 | Dusp23 | -1.713428275 | 2.96E-07 | 5.72E-06 |
| Spint1 | 1.26397 | 7.82E-15 | 8.56E-13 | Maff | -1.717272641 | 0.0085887 | 0.034839107 |
| Scarb1 | 1.26374 | 1.30E-16 | 1.94E-14 | E230032D23F | -1.719267818 | 0.0002917 | 0.002203446 |
| Gm5281 | 1.26346 | 0.008855 | 0.03572 | C330018A13F | -1.722151653 | 0.0005182 | 0.003513254 |
| Gclc | 1.26236 | 0.00084 | 0.005255 | Clybl | -1.723258953 | 1.57E-11 | 8.39E-10 |
| Wsb2 | 1.26108 | 2.15E-07 | 4.37E-06 | Batf2 | -1.72472919 | 0.0004289 | 0.003017701 |
| Osgin1 | 1.26091 | 4.57E-08 | 1.12E-06 | Gm22068 | -1.729006098 | 0.0001273 | 0.00109075 |
| Klk9 | 1.25921 | 0.001938 | 0.010506 | Gm16118 | -1.731722739 | 6.35E-15 | 7.10E-13 |
| Cenpv | 1.25917 | 5.32E-05 | 0.000518 | Syt3 | -1.743663243 | 4.27E-08 | 1.06E-06 |
| Pelp1 | 1.25426 | 9.83E-14 | 8.41E-12 | D5Ertd605e | -1.751570141 | 6.36E-12 | 3.66E-10 |
| Amfr | 1.25418 | 0.000902 | 0.005591 | Gpx7 | -1.751719386 | 0.0121875 | 0.045956828 |
| Dennd3 | 1.25182 | 0.001203 | 0.007069 | Rab7b | -1.754858919 | 6.63E-05 | 0.000619557 |
| Gm1966 | 1.25012 | 5.66E-07 | 1.00E-05 | Igfbp5 | -1.759391996 | 0.0103207 | 0.040183402 |
| Lmbr1l | 1.24953 | 2.71E-13 | 2.10E-11 | Tmem40 | -1.766747669 | 0.0123629 | 0.046457754 |
| Sik3 | 1.24834 | 1.41E-06 | 2.25E-05 | Acbd4 | -1.767248305 | 1.55E-13 | 1.25E-11 |
| Dpep2 | 1.2479 | 0.009205 | 0.036786 | Itga7 | -1.769120372 | 0.0071579 | 0.030087048 |
| Neurl1a | 1.24551 | 5.23E-08 | 1.26E-06 | Gramd3 | -1.769451196 | 1.25E-07 | 2.69E-06 |
| Gm43909 | 1.24207 | 0.004372 | 0.020287 | Epor | -1.769740269 | 2.00E-07 | 4.11E-06 |
| Arhgef39 | 1.24148 | 2.00E-05 | 0.000225 | Ripor2 | -1.770258689 | 1.11E-13 | 9.28E-12 |
| Efnb2 | 1.24132 | 0.005514 | 0.024489 | Rnu3b1 | -1.77122799 | 2.46E-06 | 3.67E-05 |
| Tbkbp1 | 1.23906 | 1.46E-09 | 5.16E-08 | Lgals3 | -1.774209846 | 4.17E-12 | 2.48E-10 |
| B330016D10Ri | 1.239 | 0.000895 | 0.005559 | Icos | -1.775198321 | 2.67E-08 | 7.02E-07 |
| Mms22l | 1.2298 | 1.99E-14 | 1.93E-12 | Patl2 | -1.779154013 | 0.0004774 | 0.003289583 |
| Ube2o | 1.22876 | 8.41E-06 | 0.000106 | Smyd2 | -1.787016658 | 7.75E-06 | 9.91E-05 |
| Zfp316 | 1.22777 | 0.003576 | 0.017227 | Gm24924 | -1.789035019 | 1.55E-10 | 6.89E-09 |
| Peg10 | 1.22725 | 0.008284 | 0.03389 | Tspan32 | -1.793288565 | 7.46E-27 | 3.46E-24 |
| Itgb7 | 1.22709 | 0.004361 | 0.020253 | Gm12918 | -1.794998342 | 0.0080238 | 0.033028827 |
| Gm23530 | 1.22696 | 0.001932 | 0.010486 | Eva1a | -1.79984129 | 7.35E-18 | 1.44E-15 |
| Arfgef1 | 1.22062 | 8.17E-11 | 3.86E-09 | Hebp2 | -1.802229594 | 0.0017034 | 0.009460046 |
| Ncapd2 | 1.22053 | 9.22E-05 | 0.000824 | 4930589L23R | -1.806737023 | 0.0026154 | 0.01337303 |
| 5730435O14Ri | 1.2189 | 0.000559 | 0.003745 | Slc2a10 | -1.807194082 | 0.0026794 | 0.013614403 |

| | | | | | | | |
|---|---|---|---|---|---|---|---|
| Rcc2 | 1.21581 | 7.09E-08 | 1.64E-06 | Hist1h1d | -1.808646149 | 0.0002185 | 0.001739446 |
| Src | 1.21352 | 2.37E-09 | 8.00E-08 | Fcgrt | -1.811512381 | 1.32E-27 | 7.20E-25 |
| Pde7a | 1.21303 | 5.40E-06 | 7.21E-05 | Tmem26 | -1.819204541 | 0.0047149 | 0.021641601 |
| Ssbp3 | 1.20963 | 3.49E-06 | 4.99E-05 | Fkbp1b | -1.819719296 | 5.70E-07 | 1.01E-05 |
| Pom121 | 1.20657 | 6.83E-07 | 1.18E-05 | Cst7 | -1.821883542 | 6.18E-18 | 1.25E-15 |
| Notch4 | 1.206 | 4.88E-07 | 8.81E-06 | Ccl6 | -1.823206309 | 2.04E-29 | 1.49E-26 |
| Gm16897 | 1.20353 | 1.59E-09 | 5.53E-08 | Elovl6 | -1.851091759 | 6.29E-05 | 0.000592651 |
| Plxna4 | 1.19916 | 5.85E-11 | 2.84E-09 | Epb41l3 | -1.852180581 | 3.02E-20 | 8.26E-18 |
| Gm44686 | 1.1979 | 0.001011 | 0.006121 | Gpr65 | -1.859690625 | 1.17E-15 | 1.53E-13 |
| Gm45206 | 1.19646 | 0.011905 | 0.045037 | Apob | -1.860886062 | 0.0015418 | 0.008720671 |
| Ttc7b | 1.19628 | 0.000337 | 0.002469 | S1pr1 | -1.863041371 | 1.05E-16 | 1.62E-14 |
| Soga1 | 1.19353 | 0.008317 | 0.033982 | Mrgpre | -1.864701911 | 1.48E-14 | 1.51E-12 |
| Itga6 | 1.19333 | 1.75E-15 | 2.20E-13 | 4833445I07Ri | -1.868899539 | 2.22E-05 | 0.000244201 |
| Mcc | 1.19091 | 0.00014 | 0.001189 | Gstm7 | -1.870034701 | 0.0022529 | 0.011899313 |
| Tpbgl | 1.18914 | 0.00198 | 0.010704 | Gm15890 | -1.874954046 | 1.91E-07 | 3.97E-06 |
| Map3k3 | 1.18852 | 4.00E-08 | 1.00E-06 | Rai14 | -1.875389297 | 0.0002975 | 0.00223925 |
| Rad51 | 1.18691 | 0.012235 | 0.046111 | Rftn1 | -1.881754565 | 1.18E-13 | 9.79E-12 |
| Cdkn3 | 1.18422 | 0.006068 | 0.026336 | Aldh1b1 | -1.885101846 | 4.70E-06 | 6.41E-05 |
| Myo1c | 1.18359 | 1.57E-13 | 1.26E-11 | Rnu3b3 | -1.894856556 | 0.0004071 | 0.00288795 |
| Ppm1e | 1.18326 | 2.58E-09 | 8.52E-08 | Thsd1 | -1.898761445 | 4.17E-09 | 1.32E-07 |
| Csf2ra | 1.18195 | 1.15E-14 | 1.23E-12 | Gm43351 | -1.898788179 | 1.02E-05 | 0.000124809 |
| Rnf103 | 1.17895 | 1.74E-07 | 3.62E-06 | 4930412C18F | -1.902017461 | 0.0022961 | 0.012085563 |
| Jak2 | 1.17799 | 3.13E-10 | 1.27E-08 | Mfsd4b3 | -1.90780748 | 0.0057331 | 0.025181821 |
| Rxfp1 | 1.17461 | 8.56E-08 | 1.94E-06 | Mid1-ps1 | -1.908554881 | 0.0043862 | 0.020334185 |
| Unc13a | 1.17452 | 9.85E-10 | 3.55E-08 | 3110070M22 | -1.915307209 | 0.0003004 | 0.002254043 |
| Il18rap | 1.17178 | 0.004769 | 0.021819 | Lrrc34 | -1.929563236 | 0.0005609 | 0.003749498 |
| Cop1 | 1.16872 | 2.87E-09 | 9.47E-08 | Gstt3 | -1.933278049 | 4.58E-05 | 0.000455958 |
| Card11 | 1.16361 | 1.89E-07 | 3.91E-06 | Cd5l | -1.939413761 | 6.90E-05 | 0.000640101 |
| Gm26910 | 1.1636 | 6.68E-05 | 0.000623 | AF529169 | -1.948068223 | 0.0023346 | 0.012242506 |
| Stxbp5 | 1.16213 | 1.54E-08 | 4.28E-07 | Gm37125 | -1.952250568 | 4.93E-18 | 1.01E-15 |
| Suds3 | 1.16172 | 8.33E-05 | 0.000754 | Apoe | -1.956641673 | 2.59E-48 | 5.67E-45 |
| Ube2q1 | 1.16157 | 4.78E-13 | 3.49E-11 | Gulp1 | -1.98057005 | 0.0045395 | 0.020944907 |
| Gm26762 | 1.16048 | 5.05E-06 | 6.81E-05 | Gm28707 | -1.985276199 | 0.0003216 | 0.002379953 |
| A130006I12Rik | 1.15886 | 0.011898 | 0.045019 | 2810442N19F | -1.995188552 | 0.0005927 | 0.003931668 |
| Rgs14 | 1.15851 | 1.10E-13 | 9.28E-12 | Gm25360 | -1.999890856 | 6.29E-06 | 8.25E-05 |
| Eme1 | 1.15764 | 0.010656 | 0.041187 | Armcx6 | -2.000585496 | 4.64E-06 | 6.34E-05 |
| Gen1 | 1.15707 | 0.009976 | 0.039162 | A530064D06I | -2.004636089 | 0.0028747 | 0.014389348 |
| 9530085L11Ril | 1.15634 | 0.00374 | 0.01787 | Ispd | -2.007327446 | 0.0009066 | 0.005617524 |
| Smurf1 | 1.15366 | 0.011364 | 0.04343 | Rnu12 | -2.009185019 | 9.60E-05 | 0.000851801 |
| Gm17709 | 1.15249 | 0.004171 | 0.019495 | C1rl | -2.016092723 | 2.08E-19 | 5.14E-17 |
| Rapgef5 | 1.15245 | 4.00E-07 | 7.51E-06 | Sfn | -2.01789146 | 0.0008553 | 0.005340789 |
| Map4k4 | 1.15199 | 7.11E-11 | 3.39E-09 | Tlcd2 | -2.022716418 | 3.93E-17 | 6.69E-15 |
| Amigo1 | 1.14928 | 3.32E-08 | 8.50E-07 | Nrap | -2.024367026 | 0.0028981 | 0.014492435 |
| Hs3st3b1 | 1.14888 | 2.47E-09 | 8.24E-08 | Cpxm1 | -2.024482704 | 0.013087 | 0.048395282 |
| Cdc42bpb | 1.14788 | 1.68E-06 | 2.62E-05 | Map7d3 | -2.026288145 | 6.67E-05 | 0.000622678 |
| Tada2b | 1.14758 | 8.94E-06 | 0.000112 | Taco1os | -2.03344541 | 0.0065169 | 0.027859077 |
| Cd34 | 1.14756 | 2.62E-12 | 1.65E-10 | Snord13 | -2.033722588 | 2.05E-06 | 3.14E-05 |

| | | | | | | | |
|---|---|---|---|---|---|---|---|
| 1190002N15Ri | 1.14663 | 0.000177 | 0.00145 | Acvrl1 | -2.054391801 | 3.65E-21 | 1.05E-18 |
| Cmip | 1.14551 | 2.11E-05 | 0.000235 | 2810468N07F | -2.054837154 | 1.39E-11 | 7.45E-10 |
| Slc12a9 | 1.14523 | 3.88E-21 | 1.10E-18 | Ccdc89 | -2.060644982 | 0.0012545 | 0.007334235 |
| Gm996 | 1.14382 | 5.72E-05 | 0.000549 | Anxa5 | -2.062654244 | 2.26E-35 | 2.47E-32 |
| Sorl1 | 1.14351 | 2.92E-08 | 7.59E-07 | Gm35611 | -2.085608595 | 0.0094964 | 0.037653624 |
| Scrib | 1.1431 | 7.17E-09 | 2.13E-07 | Ctse | -2.086546371 | 0.0015489 | 0.008748601 |
| Srsf9 | 1.14096 | 0.001866 | 0.010185 | 2010204K13F | -2.089745984 | 0.000191 | 0.001548564 |
| Gigyf1 | 1.14078 | 2.89E-10 | 1.19E-08 | Gm11545 | -2.095057461 | 2.20E-05 | 0.000242786 |
| Ect2 | 1.13984 | 0.003296 | 0.016121 | Ms4a4c | -2.104559511 | 1.59E-06 | 2.50E-05 |
| Trim44 | 1.13939 | 6.77E-10 | 2.52E-08 | Gm14230 | -2.108379099 | 0.0049053 | 0.022328398 |
| E230001N04Ri | 1.13807 | 0.005219 | 0.023483 | Gm16045 | -2.108701485 | 0.0086466 | 0.035036928 |
| Treml2 | 1.13542 | 2.34E-10 | 9.87E-09 | Mfsd7a | -2.112065705 | 9.84E-10 | 3.55E-08 |
| Galnt1 | 1.1322 | 6.34E-09 | 1.92E-07 | D030055H07I | -2.118096041 | 0.0052888 | 0.02372504 |
| Gm43272 | 1.13187 | 0.001686 | 0.009383 | Chst7 | -2.119261581 | 1.20E-05 | 0.000144262 |
| Tnks | 1.1292 | 3.97E-09 | 1.26E-07 | Gm22513 | -2.122633613 | 4.18E-08 | 1.04E-06 |
| Sdk1 | 1.12819 | 1.64E-08 | 4.53E-07 | Gm26244 | -2.130397704 | 2.42E-06 | 3.62E-05 |
| Cd300lf | 1.12785 | 5.02E-06 | 6.77E-05 | 1700003F12R | -2.13962724 | 1.53E-05 | 0.000178219 |
| Gm15832 | 1.12678 | 0.010819 | 0.041678 | Gm23444 | -2.148972551 | 0.0018884 | 0.010289973 |
| Ccnyl1 | 1.12518 | 1.43E-05 | 0.000168 | Il22ra2 | -2.158390762 | 0.0015876 | 0.008920479 |
| Cebpa | 1.12436 | 0.00976 | 0.038481 | Gm24949 | -2.168440917 | 1.70E-07 | 3.56E-06 |
| Rnf139 | 1.12373 | 1.76E-14 | 1.73E-12 | Plin3 | -2.177794189 | 4.12E-27 | 1.97E-24 |
| Fgfrl1 | 1.12267 | 0.000161 | 0.001346 | Ascl2 | -2.184975779 | 4.10E-08 | 1.02E-06 |
| Tbc1d9 | 1.1222 | 2.30E-10 | 9.76E-09 | Cd79b | -2.185423329 | 1.66E-21 | 4.90E-19 |
| Cenpe | 1.12155 | 0.006818 | 0.028849 | Ccr3 | -2.18766168 | 4.26E-07 | 7.88E-06 |
| Ppp2r5a | 1.12135 | 1.01E-10 | 4.66E-09 | Bcl2l2 | -2.192959338 | 9.39E-05 | 0.000835829 |
| Tmem64 | 1.12071 | 3.56E-08 | 9.04E-07 | Krt10 | -2.221879639 | 3.10E-10 | 1.26E-08 |
| Ermap | 1.11994 | 4.16E-18 | 8.62E-16 | Saxo2 | -2.231020047 | 0.0018579 | 0.010148789 |
| Maz | 1.11698 | 1.77E-08 | 4.84E-07 | Gm31410 | -2.23123056 | 1.29E-08 | 3.66E-07 |
| Rbm15b | 1.11635 | 3.55E-06 | 5.05E-05 | Ttpa | -2.235847121 | 0.0131818 | 0.048640391 |
| Ankrd52 | 1.1151 | 3.92E-09 | 1.25E-07 | Ighg2b | -2.244995586 | 0.0109091 | 0.04193107 |
| Slc36a1 | 1.11402 | 6.26E-12 | 3.62E-10 | Slc14a1 | -2.253940913 | 1.26E-16 | 1.89E-14 |
| Slc19a2 | 1.11329 | 1.52E-09 | 5.34E-08 | Ankrd34a | -2.257030447 | 0.0041311 | 0.019350479 |
| Irs2 | 1.11176 | 7.10E-08 | 1.64E-06 | Gm36888 | -2.265148749 | 0.0090075 | 0.036167204 |
| Ago2 | 1.11145 | 5.03E-13 | 3.62E-11 | Rapsn | -2.265985833 | 5.97E-29 | 3.66E-26 |
| Klf16 | 1.11067 | 0.000377 | 0.002711 | Acot1 | -2.272103619 | 0.0002843 | 0.002164154 |
| Patz1 | 1.10943 | 2.47E-06 | 3.68E-05 | Nme4 | -2.286948238 | 8.58E-14 | 7.47E-12 |
| Pdcd7 | 1.10651 | 2.43E-08 | 6.48E-07 | Car11 | -2.287417714 | 2.17E-08 | 5.83E-07 |
| N4bp1 | 1.1062 | 6.44E-12 | 3.70E-10 | Gm13391 | -2.292677822 | 0.0037701 | 0.017989594 |
| Tmem86b | 1.10583 | 5.17E-10 | 1.99E-08 | C530050E15F | -2.314497673 | 0.0046471 | 0.021387988 |
| Fnip2 | 1.10512 | 8.41E-11 | 3.97E-09 | Hacd4 | -2.316414852 | 0.0001194 | 0.001030761 |
| Micall1 | 1.10491 | 8.59E-11 | 4.03E-09 | Gm9999 | -2.318653071 | 0.005072 | 0.022970995 |
| Postn | 1.10414 | 2.06E-12 | 1.32E-10 | Lgi4 | -2.320025348 | 2.70E-07 | 5.28E-06 |
| Sort1 | 1.10354 | 4.54E-09 | 1.41E-07 | 5930430L01R | -2.321128114 | 4.35E-06 | 5.98E-05 |
| Rad51ap1 | 1.10221 | 0.003395 | 0.016498 | Dapk1 | -2.322437729 | 5.03E-13 | 3.62E-11 |
| Srf | 1.10021 | 3.56E-06 | 5.06E-05 | Kazald1 | -2.327233704 | 0.0031739 | 0.015626455 |
| Trappc10 | 1.09784 | 4.74E-07 | 8.59E-06 | Krt15 | -2.342743234 | 0.0027852 | 0.014028355 |
| Gm39348 | 1.09716 | 0.001627 | 0.009091 | Gm44702 | -2.350969804 | 0.0010965 | 0.006550517 |

| | | | |
|---|---|---|---|
| Brpf3 | 1.09674 | 2.89E-05 | 0.000307 |
| Spast | 1.09596 | 2.71E-16 | 3.81E-14 |
| Ncor1 | 1.09481 | 2.77E-07 | 5.42E-06 |
| Rnf169 | 1.09458 | 1.37E-12 | 9.09E-11 |
| 4933439K11Ri | 1.09401 | 5.81E-05 | 0.000556 |
| Tnrc6c | 1.09265 | 5.61E-10 | 2.12E-08 |
| Ppfia1 | 1.08986 | 1.18E-07 | 2.55E-06 |
| Hist1h2bc | 1.08814 | 7.62E-11 | 3.62E-09 |
| Papd5 | 1.08487 | 9.69E-09 | 2.83E-07 |
| Nrep | 1.08429 | 4.03E-12 | 2.41E-10 |
| Pdgfb | 1.08271 | 4.63E-10 | 1.81E-08 |
| Dpy19l1 | 1.08247 | 1.21E-05 | 0.000145 |
| Gm47795 | 1.08194 | 0.001468 | 0.008366 |
| Tle1 | 1.08169 | 1.14E-08 | 3.27E-07 |
| Zfp318 | 1.08088 | 4.41E-06 | 6.05E-05 |
| Il7r | 1.07987 | 1.68E-11 | 8.80E-10 |
| Srpk3 | 1.07982 | 9.25E-11 | 4.29E-09 |
| Gpd1l | 1.07941 | 5.18E-15 | 6.01E-13 |
| Nedd9 | 1.0788 | 0.000698 | 0.004503 |
| Fxr2 | 1.07797 | 2.14E-06 | 3.27E-05 |
| Ankib1 | 1.07763 | 1.83E-12 | 1.20E-10 |
| Ammecr1 | 1.07734 | 0.000392 | 0.002809 |
| Itgav | 1.07687 | 4.46E-15 | 5.22E-13 |
| Vipr1 | 1.07369 | 1.55E-10 | 6.88E-09 |
| Dnah17 | 1.06961 | 0.006226 | 0.026888 |
| Ccne2 | 1.06705 | 0.001446 | 0.00827 |
| Igfbp7 | 1.06559 | 0.002459 | 0.012737 |
| Grk2 | 1.06183 | 3.60E-08 | 9.12E-07 |
| Mmp15 | 1.06154 | 0.002232 | 0.011815 |
| Hira | 1.06055 | 0.000615 | 0.00405 |
| Ache | 1.05914 | 4.24E-07 | 7.85E-06 |
| Atp1a3 | 1.05669 | 2.29E-05 | 0.000251 |
| 4930517O19Ri | 1.05476 | 0.000327 | 0.002407 |
| Cdc73 | 1.054 | 1.59E-11 | 8.43E-10 |
| Upf1 | 1.0531 | 8.07E-07 | 1.38E-05 |
| Slc16a6 | 1.05262 | 3.32E-19 | 7.71E-17 |
| Btg3 | 1.05186 | 0.001005 | 0.006095 |
| Il6ra | 1.04998 | 3.92E-17 | 6.69E-15 |
| Crim1 | 1.04914 | 1.84E-05 | 0.000209 |
| Necap1 | 1.04883 | 1.46E-14 | 1.51E-12 |
| Mcmbp | 1.04881 | 2.24E-07 | 4.52E-06 |
| Mast2 | 1.0466 | 3.16E-06 | 4.58E-05 |
| Bod1 | 1.04647 | 6.62E-07 | 1.15E-05 |
| Atp2a2 | 1.04604 | 7.67E-07 | 1.32E-05 |
| Rpia | 1.04462 | 8.08E-07 | 1.38E-05 |
| Taf5 | 1.04302 | 7.00E-05 | 0.000648 |
| Dgka | 1.03893 | 3.04E-16 | 4.23E-14 |

| | | | |
|---|---|---|---|
| Krt14 | -2.357276054 | 5.31E-07 | 9.49E-06 |
| Gm17025 | -2.396271307 | 1.48E-05 | 0.000173727 |
| Zfp57 | -2.403249703 | 0.0057903 | 0.025354332 |
| Gm24407 | -2.416150521 | 1.75E-07 | 3.64E-06 |
| Krt5 | -2.435075244 | 9.81E-05 | 0.000868863 |
| Stap2 | -2.456670075 | 0.0027072 | 0.013730654 |
| Pnck | -2.472261551 | 4.27E-06 | 5.90E-05 |
| AW112010 | -2.516134699 | 8.36E-07 | 1.42E-05 |
| Gm24950 | -2.581809013 | 0.0004863 | 0.003343487 |
| D630033O11 | -2.6153458 | 0.0010202 | 0.006163441 |
| Rnu3a | -2.640062695 | 0.000225 | 0.001784453 |
| Pdzk1ip1 | -2.642198055 | 1.37E-05 | 0.000162043 |
| Gm38091 | -2.644609658 | 9.46E-13 | 6.47E-11 |
| D7Ertd128e | -2.652965308 | 1.23E-06 | 2.00E-05 |
| Gm25099 | -2.685314078 | 0.0001618 | 0.001347339 |
| Cd22 | -2.69838021 | 4.87E-16 | 6.60E-14 |
| Tll2 | -2.718112045 | 0.0010928 | 0.006536097 |
| Gm24265 | -2.726387058 | 2.95E-13 | 2.26E-11 |
| Gm45778 | -2.737592636 | 3.65E-07 | 6.91E-06 |
| Fam71a | -2.745856662 | 3.42E-05 | 0.000353913 |
| Treml4 | -2.752204682 | 0.0007296 | 0.004663346 |
| Ly6k | -2.787426416 | 0.0092878 | 0.037047081 |
| Six5 | -2.827012145 | 0.0001645 | 0.001364752 |
| Emp1 | -2.875477395 | 6.19E-05 | 0.000585444 |
| Mmp12 | -2.88863559 | 1.08E-07 | 2.36E-06 |
| Calm4 | -2.895728345 | 3.81E-07 | 7.20E-06 |
| Krt1 | -2.916446117 | 9.01E-06 | 0.000112636 |
| Fam178b | -2.93157912 | 1.28E-06 | 2.06E-05 |
| Anxa8 | -2.955032493 | 0.0006409 | 0.00418658 |
| BC064078 | -3.072262694 | 6.31E-06 | 8.26E-05 |
| Ccl24 | -3.107744902 | 1.03E-16 | 1.60E-14 |
| Krt2 | -3.139069659 | 3.35E-06 | 4.80E-05 |
| Tmem82 | -3.184404338 | 1.77E-06 | 2.74E-05 |
| Gpnmb | -3.295433322 | 2.89E-05 | 0.000306919 |
| Aqp3 | -3.339111332 | 6.13E-05 | 0.000581047 |
| Ly6d | -3.372048842 | 1.41E-06 | 2.24E-05 |
| Dsc3 | -3.561889888 | 1.22E-06 | 1.99E-05 |
| Ccr1 | -3.590969212 | 1.65E-10 | 7.22E-09 |
| Serpinb2 | -3.640199767 | 6.49E-08 | 1.52E-06 |
| Krtdap | -3.901435305 | 1.58E-06 | 2.50E-05 |
| Lypd3 | -3.952039145 | 6.29E-05 | 0.000592651 |

| | | | |
|---|---|---|---|
| Atrnl1 | 1.03727 | 0.007902 | 0.032596 |
| Gm13563 | 1.03666 | 4.51E-05 | 0.00045 |
| Stk11 | 1.03562 | 9.26E-07 | 1.56E-05 |
| Galnt2 | 1.03507 | 0.004307 | 0.020022 |
| Tk1 | 1.03328 | 4.92E-06 | 6.67E-05 |
| Liph | 1.03323 | 5.66E-12 | 3.31E-10 |
| Bmt2 | 1.03009 | 7.74E-07 | 1.33E-05 |
| Mycbp | 1.02991 | 8.86E-09 | 2.61E-07 |
| Zfp710 | 1.02891 | 1.24E-16 | 1.89E-14 |
| Piezo1 | 1.02862 | 9.76E-07 | 1.62E-05 |
| Avpr2 | 1.02752 | 0.000675 | 0.004379 |
| Gm32098 | 1.02439 | 0.012789 | 0.047594 |
| Fam102b | 1.02384 | 6.61E-11 | 3.19E-09 |
| Ubp1 | 1.02306 | 3.96E-08 | 9.94E-07 |
| Gcnt2 | 1.02244 | 2.82E-08 | 7.37E-07 |
| Snap25 | 1.02033 | 0.000773 | 0.004906 |
| Gm15907 | 1.02024 | 0.008192 | 0.033578 |
| Phf10 | 1.0197 | 5.19E-13 | 3.72E-11 |
| Tgfbi | 1.01942 | 3.14E-17 | 5.60E-15 |
| Fam160b1 | 1.01724 | 6.55E-08 | 1.53E-06 |
| BC030336 | 1.01655 | 2.35E-07 | 4.71E-06 |
| Larp1 | 1.01648 | 8.05E-08 | 1.84E-06 |
| Gm29093 | 1.01599 | 0.001846 | 0.010099 |
| Faf1 | 1.01594 | 1.75E-05 | 0.000201 |
| Otud4 | 1.0158 | 1.69E-10 | 7.36E-09 |
| Ube2h | 1.01563 | 2.68E-11 | 1.36E-09 |
| Gm43273 | 1.01404 | 4.23E-05 | 0.000426 |
| Pask | 1.00827 | 0.003101 | 0.015318 |
| Zfp703 | 1.00802 | 1.96E-07 | 4.04E-06 |
| Tmem245 | 1.00665 | 9.60E-12 | 5.28E-10 |
| Rhobtb2 | 1.00631 | 1.40E-08 | 3.94E-07 |
| Dagla | 1.00614 | 5.95E-12 | 3.45E-10 |
| Rai1 | 1.00599 | 4.06E-09 | 1.29E-07 |
| Cdk19 | 1.0025 | 6.04E-14 | 5.41E-12 |
