## Supplementary File 2 for "Dicer deficiency in microglia leads to accelerated demyelination and failed remyelination"

**Supplementary file 2:** List of enriched biological processed of DEGs from flow sorted microglia isolated from naïve Cx3cr1$^{creERT2}$ Cre- control and Cre+ mutant animals.

| Category | Term | Count | % | PValue | Genes | List Total | Pop Hits | Pop Total | Fold Enrich | Bonferroni | Benjamini | FDR |
|---|---|---|---|---|---|---|---|---|---|---|---|---|
| GOTERM_I | GO:00513( | 49 | 4.64455 | 4.80E-11 | 242785, 26 | 811 | 374 | 18082 | 2.921125 | 1.51E-07 | 1.51E-07 | 1.50E-07 |
| GOTERM_I | GO:00070 | 64 | 6.066351 | 6.81E-10 | 242785, 26 | 811 | 614 | 18082 | 2.324006 | 2.14E-06 | 8.05E-07 | 8.02E-07 |
| GOTERM_I | GO:00070( | 39 | 3.696682 | 7.69E-10 | 242785, 26 | 811 | 277 | 18082 | 3.139138 | 2.41E-06 | 8.05E-07 | 8.02E-07 |
| GOTERM_I | GO:00321 | 11 | 1.042654 | 1.11E-06 | 22339, 13( | 811 | 33 | 18082 | 7.431977 | 0.003488 | 7.79E-04 | 7.76E-04 |
| GOTERM_I | GO:00064( | 53 | 5.023697 | 1.24E-06 | 17169, 171 | 811 | 576 | 18082 | 2.051535 | 0.003885 | 7.79E-04 | 7.76E-04 |
| GOTERM_I | GO:00163 | 52 | 4.92891 | 1.49E-05 | 17169, 171 | 811 | 612 | 18082 | 1.894426 | 0.045523 | 0.007765 | 0.007738 |
