## Supplementary File 3 for "Dicer deficiency in microglia leads to accelerated demyelination and failed remyelination"

**Supplementary file 3:** Ingenuity Canonical Pathways (IPA) analysis of DEGs from flow sorted microglia isolated from naïve Cx3cr1$^{creERT2}$ Cre- control and Cre+ mutant animals.

| Ingenuity Canonical Pathways | -log(p-va | Ratio | z-score | Molecules |
|---|---|---|---|---|
| Cardiac Hypertrophy Signaling (Enhanced) | 4.3 | 0.078 | 3.43 | ACVR2A,ACVR2B,ADRA2A,ADRB1,ATP2A2,CALML5,DVL1,ENPP6,FGFRL1,FZD8,GNG7,GSK3A,HSPB1,IL12 |
| Insulin Secretion Signaling Pathway | 0 | 0.037 | 2.828 | AGO2,GCK,ITPR3,JAK2,PRKACA,PRKAR2A,PRKCQ,SNAP25,SRC |
| Natural Killer Cell Signaling | 0.812 | 0.0558 | 2.714 | HSPA1A/HSPA1B,Hspa1b,IL12RB2,IL18RAP,JAK2,KLRD1,MAP3K3,MAP3K5,MAP3K9,PRKCQ,RAC1 |
| Kinetochore Metaphase Signaling Pathway | 5.03 | 0.149 | 2.673 | AURKB,BIRC5,BOD1,CCNB1,CDCA8,CENPA,CENPE,KNL1,MXD3,NEK2,NUF2,PLK1,PPP2R5A,SKA3,SPC24 |
| Cardiac Hypertrophy Signaling | 1.85 | 0.0708 | 2.673 | ADRA2A,ADRB1,CALML5,GNB5,GNG7,HSPB1,IL6R,MAP3K3,MAP3K5,MAP3K9,MAPKAPK2,PRKACA, |
| GNRH Signaling | 1.1 | 0.0636 | 2.53 | CALML5,GNG7,ITPR3,MAP3K3,MAP3K5,MAP3K9,PRKACA,PRKAR2A,PRKCQ,RAC1,SRC |
| BAG2 Signaling Pathway | 2.2 | 0.14 | 2.449 | HSP90AA1,HSPA1A/HSPA1B,Hspa1b,MAPKAPK2,MYC,PSMA8 |
| Unfolded protein response | 1.66 | 0.107 | 2.449 | AMFR,CEBPA,CEBPB,HSPA1A/HSPA1B,Hspa1b,MAP3K5 |
| Neuropathic Pain Signaling In Dorsal Horn Neurons | 0.686 | 0.0594 | 2.449 | ITPR3,KCNN4,PRKACA,PRKAR2A,PRKCQ,SRC |
| Gαs Signaling | 2.03 | 0.0935 | 2.333 | ADRB1,AVPR2,GNB5,GNG7,GPER1,HCAR2,PRKACA,PRKAR2A,SRC,VIPR1 |
| Dopamine-DARPP32 Feedback in cAMP Signaling | 0.963 | 0.0613 | 2.333 | ATP2A2,CALML5,CSNK1G2,CSNK1G3,ITPR3,PPM1J,PPP2R5A,PRKACA,PRKAR2A,PRKCQ |
| Cell Cycle Regulation by BTG Family Proteins | 3.29 | 0.189 | 2.236 | CCNE1,CCNE2,E2F1,E2F7,E2F8,PPM1J,PPP2R5A |
| IGF-1 Signaling | 1.67 | 0.0865 | 2.236 | IGFBP4,IGFBP5,IGFBP7,IRS2,JAK2,PRKACA,PRKAR2A,SFN,SRF |
| Agrin Interactions at Neuromuscular Junction | 1.07 | 0.0769 | 2.236 | DVL1,ITGA6,ITGB3,RAC1,RAPSN,SRC |
| NF-κB Activation by Viruses | 0.987 | 0.0732 | 2.236 | CCR5,ITGA6,ITGAV,ITGB3,ITGB5,PRKCQ |
| GPCR-Mediated Integration of Enteroendocrine Signaling Exemplified by an L Cell | 0.799 | 0.0685 | 2.236 | ADRB1,ITPR3,PRKACA,PRKAR2A,VIPR1 |
| Reelin Signaling in Neurons | 0.613 | 0.0543 | 2.236 | APBB1,APOE,ITGA6,ITGB3,MAP3K9,RAC1,SRC |
| CDK5 Signaling | 0.6 | 0.0556 | 2.236 | CABLES1,ITGA6,PPM1J,PPP2R5A,PRKACA,PRKAR2A |
| Thrombin Signaling | 0.365 | 0.0433 | 2.236 | GATA2,GNB5,GNG7,ITPR3,PRKCQ,RAC1,RHOBTB2,RHOQ,SRC |
| HGF Signaling | 0.354 | 0.045 | 2.236 | MAP3K3,MAP3K5,MAP3K9,PRKCQ,RAC1 |
| Estrogen-mediated S-phase Entry | 5.34 | 0.308 | 2.121 | CCNA2,CCNE1,CCNE2,E2F1,E2F7,E2F8,ESR1,MYC |
| Glioma Invasiveness Signaling | 3.21 | 0.135 | 2.121 | HMMR,ITGAV,ITGB3,ITGB5,MMP9,PLAUR,RAC1,RHOBTB2,RHOQ,TIMP4 |
| Superpathway of Inositol Phosphate Compounds | 0.281 | 0.0402 | 2.121 | CDC25C,DUSP23,INPP5A,PI4K2A,PLPP6,PPFIA1,PPP1R12C,PPP2R5A |
| Leukocyte Extravasation Signaling | 1.62 | 0.0711 | 2.111 | CLDN10,CLDN12,ITGA6,ITGB3,JAM3,MMP11,MMP12,MMP15,MMP8,MMP9,PRKCQ,RAC1,SRC,TIMP4 |
| IL-8 Signaling | 1.28 | 0.065 | 2.111 | GNB5,GNG7,ITGAV,ITGB3,ITGB5,MAP4K4,MMP9,PRKCQ,RAC1,RHOBTB2,RHOQ,SRC,VEGFA |
| cAMP-mediated signaling | 2.76 | 0.0833 | 2.065 | ADORA1,ADRA2A,ADRB1,AVPR2,CALML5,ENPP6,GPER1,GRK2,HCAR2,PDE3B,PDE7A,PDE8A,PRKACA, |
| Dermatan Sulfate Degradation (Metazoa) | 2.43 | 0.235 | 2 | FGFRL1,HYAL4,Spam1,SPAM1 |
| tRNA Splicing | 1.06 | 0.093 | 2 | ENPP6,PDE3B,PDE7A,PDE8A |
| Tec Kinase Signaling | 0.706 | 0.0549 | 2 | GNB5,GNG7,JAK2,PRKCQ,RAC1,RHOBTB2,RHOQ,SRC,TNF |
| Netrin Signaling | 0.599 | 0.0615 | 2 | PRKACA,PRKAR2A,RAC1,UNC5A |
| Growth Hormone Signaling | 0.517 | 0.0563 | 2 | CEBPA,JAK2,PRKCQ,SRF |
| FGF Signaling | 0.376 | 0.0476 | 2 | FGFRL1,MAP3K5,MAPKAPK2,RAC1 |
| Fcγ Receptor-mediated Phagocytosis in Macrophages and Monocytes | 0.295 | 0.0426 | 2 | ARF6,PRKCQ,RAC1,SRC |
| IL-15 Production | 0 | 0.0331 | 2 | CSF2RA,DYRK3,JAK2,SRC |
| Putrescine Degradation III | 2.08 | 0.19 | -2 | ALDH1B1,ALDH2,MAOA,SAT2 |
| Glutathione Redox Reactions I | 1.87 | 0.167 | -2 | GPX7,GSTM2,GSTT2/GSTT2B,MGST3 |
| BEX2 Signaling Pathway | 0.425 | 0.0506 | -2 | LGALS1,PPM1J,PPP2R5A,VEGFA |
| Neuroprotective Role of THOP1 in Alzheimer's Disease | 0 | 0.0345 | -2 | C1R,MMP9,PRKACA,PRKAR2A |
| Coronavirus Pathogenesis Pathway | 0 | 0.0267 | -2 | CCL5,E2F1,E2F7,E2F8 |
| Necroptosis Signaling Pathway | 0 | 0.0318 | -2.236 | CAPN2,DAPK1,PYGL,RIPK3,TNF |
| Xenobiotic Metabolism PXR Signaling Pathway | 5.68 | 0.12 | -2.294 | ALDH1B1,ALDH2,CHST10,CHST2,CHST7,GSTM2,GSTM3,GSTM4,GSTM5,GSTO2,GSTT2/GSTT2B,HS3ST3B1 |
| Apelin Adipocyte Signaling Pathway | 0.987 | 0.0732 | -2.449 | GPX7,GSTM2,GSTT2/GSTT2B,MGST3,PRKACA,PRKAR2A |
| Xenobiotic Metabolism AHR Signaling Pathway | 4.58 | 0.153 | -2.496 | AHRR,ALDH1B1,ALDH2,GSTM2,GSTM3,GSTM4,GSTM5,GSTO2,GSTT2/GSTT2B,HSP90AA1,MGST3,NCOA1,TNF |
| PPARα/RXRα Activation | 1.43 | 0.0684 | -2.496 | ACVR2A,ACVR2B,HSP90AA1,IL18RAP,IL1R2,ITGB5,JAK2,LPL,MAP4K4,NCOR1,NCOR2,PRKACA,PRKAR2A |
| VDR/RXR Activation | 2.46 | 0.115 | -2.646 | CCL5,CEBPA,CEBPB,CXCL10,IGFBP5,NCOA1,NCOR1,NCOR2,PRKCQ |
| Glutathione-mediated Detoxification | 4.6 | 0.25 | -2.828 | GSTM2,GSTM3,GSTM4,GSTM5,GSTO2,GSTT2/GSTT2B,Gstt3,MGST3 |
