## Supplementary File 4 for "Dicer deficiency in microglia leads to accelerated demyelination and failed remyelination"

**Supplementary file 4:** List of differentially expressed gene (DEGs, log2FoldChange <-1/>+1, pAdj<0.05) from flow sorted microglia isolated from Cx3cr1$^{creERT2}$ Cre- control and Cre+ mutant animals fed to CUP diet for 6 weeks.

| Gene.name | log2FoldChange | pvalue | padj | Gene.nam | log2FoldCl | pvalue | padj |
|---|---|---|---|---|---|---|---|
| Trim55 | 7.900618372 | 3.03E-05 | 0.000725 | Cd209a | -1.00553 | 0.00212 | 0.01927 |
| Gcm2 | 5.535049819 | 0.0002808 | 0.00411 | Tiam1 | -1.00799 | 0.001971 | 0.018301 |
| Gm44507 | 5.324182472 | 6.52E-05 | 0.001332 | Gm22513 | -1.0088 | 0.003332 | 0.027063 |
| Ostn | 5.132148605 | 2.92E-05 | 0.000706 | Myom1 | -1.0093 | 4.22E-06 | 0.000148 |
| Ripk4 | 5.101982431 | 5.69E-07 | 2.79E-05 | Ms4a4a | -1.01285 | 3.39E-09 | 3.24E-07 |
| Myt1l | 4.897539685 | 0.0010994 | 0.011683 | Gm23472 | -1.01306 | 0.002402 | 0.021159 |
| Ddo | 4.493354836 | 1.27E-06 | 5.58E-05 | Tet1 | -1.01724 | 0.002199 | 0.019854 |
| Gm826 | 4.309146771 | 2.84E-07 | 1.55E-05 | Tnni3 | -1.0173 | 0.000125 | 0.002201 |
| Gm34312 | 4.266906987 | 6.07E-06 | 0.000196 | Ccr5 | -1.02194 | 7.42E-08 | 4.99E-06 |
| Syt6 | 4.094843146 | 1.78E-05 | 0.000473 | Creg2 | -1.02357 | 0.002322 | 0.020678 |
| Gm8091 | 4.081094242 | 1.99E-12 | 3.73E-10 | Gm24265 | -1.03313 | 0.002699 | 0.023009 |
| Gm47926 | 3.991479362 | 4.10E-05 | 0.000927 | Gm42867 | -1.04109 | 3.49E-05 | 0.000815 |
| Lgi2 | 3.807122961 | 6.31E-05 | 0.0013 | Matk | -1.04111 | 8.08E-10 | 9.35E-08 |
| Gm12856 | 3.56615554 | 3.54E-05 | 0.000823 | S100a6 | -1.04877 | 3.97E-05 | 0.000899 |
| Olfr1330 | 3.535410455 | 1.14E-05 | 0.000327 | S100a4 | -1.07815 | 0.003163 | 0.025932 |
| Cxcl13 | 3.47674413 | 0.0001184 | 0.002123 | F13a1 | -1.08038 | 6.25E-07 | 3.03E-05 |
| Hormad2 | 3.339522208 | 1.96E-05 | 0.000511 | Dnaaf3 | -1.08208 | 2.49E-08 | 1.89E-06 |
| Gm48483 | 3.301522376 | 1.20E-05 | 0.000342 | Ahnak | -1.09063 | 0.00113 | 0.011901 |
| Gm12865 | 3.037193137 | 0.000934 | 0.010431 | Mfsd7a | -1.09352 | 4.07E-05 | 0.000922 |
| Mreg | 3.016651475 | 6.03E-06 | 0.000196 | Slc26a11 | -1.09804 | 3.78E-05 | 0.00087 |
| Olfr1328 | 2.9988816 | 0.0001862 | 0.002999 | Slc14a1 | -1.09858 | 0.000145 | 0.002452 |
| Unc5cl | 2.992012487 | 9.87E-08 | 6.51E-06 | Ly6g6e | -1.10048 | 0.001398 | 0.014059 |
| Sox7 | 2.90515804 | 2.65E-06 | 0.000102 | Dlc1 | -1.10463 | 5.22E-05 | 0.001116 |
| Dcstamp | 2.861743682 | 4.79E-07 | 2.41E-05 | Gm12279 | -1.1056 | 0.006451 | 0.043825 |
| Hyal6 | 2.839117799 | 4.51E-09 | 4.17E-07 | Gprasp2 | -1.11758 | 0.001278 | 0.013156 |
| Gm38832 | 2.776069949 | 6.83E-13 | 1.35E-10 | Zfp37 | -1.11908 | 0.000352 | 0.00492 |
| Asb4 | 2.754064269 | 0.0007533 | 0.008771 | Cxcr3 | -1.12178 | 3.32E-05 | 0.000781 |
| Kif14 | 2.7472069 | 2.07E-05 | 0.000535 | Cybb | -1.13393 | 4.97E-14 | 1.34E-11 |
| Fam19a1 | 2.741863335 | 2.05E-10 | 2.75E-08 | Trib3 | -1.14076 | 0.000161 | 0.002648 |
| Gm11707 | 2.735484853 | 3.29E-08 | 2.41E-06 | Gpr165 | -1.14942 | 0.00015 | 0.002511 |
| Gm12864 | 2.72080956 | 0.0028175 | 0.023755 | Gm26582 | -1.15138 | 0.005196 | 0.037595 |
| Kcnh4 | 2.718580954 | 5.90E-08 | 4.10E-06 | Arhgef10l | -1.15148 | 4.88E-12 | 8.69E-10 |
| Ctsw | 2.71479416 | 5.80E-14 | 1.48E-11 | Sgsh | -1.15508 | 9.98E-08 | 6.52E-06 |
| 5330426L24Rik | 2.704835882 | 1.69E-07 | 1.02E-05 | Gm39469 | -1.15867 | 3.56E-06 | 0.00013 |
| Raet1d | 2.686769123 | 6.86E-05 | 0.001376 | Gm10655 | -1.15876 | 0.00275 | 0.023294 |
| Perp | 2.677815368 | 1.46E-07 | 9.01E-06 | Slc41a2 | -1.1606 | 0.003134 | 0.025787 |
| Fam189a2 | 2.661103612 | 0.0001975 | 0.003139 | Cyp27a1 | -1.16224 | 1.37E-13 | 3.23E-11 |
| Rarb | 2.606787373 | 6.59E-05 | 0.001341 | Mamld1 | -1.16841 | 0.002041 | 0.018766 |
| B230311B06Rik | 2.573535534 | 3.70E-07 | 1.95E-05 | Gm21738 | -1.17908 | 0.005334 | 0.038328 |
| 4930455G09Rik | 2.541148759 | 0.0001755 | 0.002857 | Mxra7 | -1.19345 | 0.002906 | 0.0243 |
| Nid2 | 2.531076841 | 9.23E-06 | 0.000275 | Tlr8 | -1.21026 | 4.40E-05 | 0.000975 |

| | | | | | | | |
|---|---|---|---|---|---|---|---|
| Lrrc34 | 2.526468973 | 0.0060708 | 0.041864 | Ltb | -1.21193 | 0.000128 | 0.002229 |
| Upk1b | 2.484880609 | 2.62E-05 | 0.000648 | Ciita | -1.21529 | 2.97E-05 | 0.000714 |
| Lrr1 | 2.453435193 | 6.70E-06 | 0.000213 | Ptpn22 | -1.21555 | 1.88E-09 | 1.94E-07 |
| Gm35572 | 2.446774518 | 2.68E-05 | 0.000658 | Myof | -1.21789 | 6.11E-05 | 0.001271 |
| Kcnn4 | 2.433462021 | 0.0002165 | 0.00337 | Ctnna3 | -1.21838 | 0.002694 | 0.023009 |
| BC030867 | 2.431332007 | 7.32E-05 | 0.00144 | C1rl | -1.2256 | 6.51E-05 | 0.001332 |
| E430024P14Rik | 2.399808905 | 0.0006663 | 0.007986 | Plac8 | -1.22575 | 3.60E-06 | 0.000131 |
| Fbn2 | 2.376409753 | 0.0002506 | 0.003765 | Gm31410 | -1.22622 | 1.87E-06 | 7.70E-05 |
| Gramd1c | 2.365355751 | 0.0008279 | 0.009484 | Noxo1 | -1.22971 | 0.001282 | 0.01319 |
| Gm2065 | 2.34397858 | 1.33E-09 | 1.41E-07 | Asb11 | -1.23202 | 0.000969 | 0.010694 |
| Dennd3 | 2.341796431 | 8.02E-09 | 7.19E-07 | Slamf7 | -1.23595 | 1.02E-16 | 3.63E-14 |
| Skint3 | 2.333327695 | 1.75E-09 | 1.82E-07 | Gm21769 | -1.24791 | 0.005026 | 0.03682 |
| Gm12861 | 2.316868411 | 2.86E-06 | 0.000108 | Klhl3 | -1.25344 | 1.36E-05 | 0.000377 |
| Gm8696 | 2.309867629 | 4.92E-07 | 2.47E-05 | Rnd3 | -1.26151 | 0.007555 | 0.049077 |
| Runx3 | 2.30775485 | 5.83E-06 | 0.000191 | Gm10800 | -1.26506 | 1.43E-05 | 0.000391 |
| Alb | 2.294729255 | 4.40E-06 | 0.000153 | Itga4 | -1.28942 | 6.71E-07 | 3.22E-05 |
| Ube2c | 2.290147145 | 5.84E-45 | 2.08E-41 | Gm8113 | -1.29977 | 0.000243 | 0.0037 |
| Ache | 2.278009643 | 3.60E-14 | 1.01E-11 | Gm10801 | -1.30473 | 0.000461 | 0.006 |
| D630023F18Rik | 2.272721713 | 1.87E-06 | 7.70E-05 | Cd74 | -1.30619 | 7.32E-44 | 2.09E-40 |
| Depdc1a | 2.254293382 | 8.84E-09 | 7.72E-07 | AF529169 | -1.35751 | 2.21E-05 | 0.000565 |
| Dnaic1 | 2.230817157 | 0.0007757 | 0.008973 | Gnb3 | -1.36048 | 0.000206 | 0.003239 |
| Cited4 | 2.184317763 | 0.002393 | 0.021126 | Arhgef25 | -1.37269 | 0.004474 | 0.033872 |
| Rxrg | 2.175997312 | 0.0001565 | 0.002602 | Syt3 | -1.37685 | 0.001659 | 0.016032 |
| Hist2h3c2 | 2.174464677 | 0.0001755 | 0.002857 | Mycl | -1.38166 | 0.0003 | 0.004338 |
| Hsf2bp | 2.147204145 | 7.53E-06 | 0.000233 | Pstpip2 | -1.3833 | 0.001071 | 0.011514 |
| Htra3 | 2.132059755 | 1.32E-07 | 8.28E-06 | Gpr83 | -1.3918 | 0.000414 | 0.005548 |
| Adora1 | 2.108126414 | 0.0001194 | 0.002132 | Eno2 | -1.39219 | 0.000119 | 0.002132 |
| Prkcq | 2.078790353 | 2.52E-12 | 4.65E-10 | Ahr | -1.39984 | 0.000593 | 0.007249 |
| Sapcd2 | 2.075276878 | 2.52E-06 | 9.82E-05 | Diras2 | -1.40761 | 0.001162 | 0.012113 |
| Mcc | 2.067652564 | 5.12E-07 | 2.54E-05 | A430110L2 | -1.41953 | 0.005962 | 0.041414 |
| Olfm4 | 2.055629551 | 7.32E-05 | 0.00144 | Gm16348 | -1.43998 | 0.000247 | 0.003726 |
| Cd40 | 2.049105357 | 0.0022418 | 0.020126 | Sell | -1.44712 | 4.88E-06 | 0.000165 |
| Bub1 | 2.024385483 | 1.34E-22 | 1.01E-19 | Gm8774 | -1.44762 | 0.002577 | 0.022304 |
| Hspa1a | 2.010996335 | 0.0006729 | 0.008059 | Mir5107 | -1.45927 | 0.004799 | 0.035611 |
| Gm4316 | 1.996134776 | 0.0006913 | 0.008238 | Gda | -1.46727 | 0.000872 | 0.009893 |
| Rxfp1 | 1.988403414 | 6.00E-07 | 2.92E-05 | Ighj1 | -1.46817 | 1.27E-06 | 5.60E-05 |
| Pif1 | 1.984354694 | 5.20E-06 | 0.000174 | Ms4a4b | -1.48226 | 0.005508 | 0.03916 |
| D7Ertd443e | 1.973840342 | 9.96E-05 | 0.001843 | Olfr1033 | -1.51937 | 0.001299 | 0.013306 |
| Stil | 1.970664134 | 2.54E-11 | 3.93E-09 | Ccl22 | -1.53229 | 0.002285 | 0.020425 |
| Mki67 | 1.950768697 | 8.68E-38 | 1.77E-34 | Slc26a1 | -1.53492 | 0.000925 | 0.010334 |
| Hyal4 | 1.945747028 | 3.27E-07 | 1.74E-05 | Ahrr | -1.53582 | 0.005325 | 0.038325 |
| Il1r2 | 1.940970504 | 3.22E-05 | 0.000762 | Gm37306 | -1.53935 | 1.50E-07 | 9.21E-06 |
| Gm11998 | 1.929680946 | 1.68E-08 | 1.41E-06 | H2-Ab1 | -1.54546 | 1.34E-49 | 9.57E-46 |
| Hspa1b | 1.924742026 | 0.0002742 | 0.00405 | Rsph10b | -1.55493 | 0.004558 | 0.034236 |
| Ifi44 | 1.909163018 | 0.0023761 | 0.021029 | Stx1a | -1.57065 | 3.17E-05 | 0.000756 |
| Nek2 | 1.880793394 | 1.96E-15 | 5.93E-13 | Ccr2 | -1.57387 | 0.000358 | 0.004993 |
| Aspm | 1.878257298 | 8.28E-11 | 1.19E-08 | Gm38091 | -1.57582 | 1.09E-05 | 0.000315 |

| | | | | | | | |
|---|---|---|---|---|---|---|---|
| Etv4 | 1.864669872 | 4.08E-07 | 2.09E-05 | Gm26870 | -1.57588 | 0.000716 | 0.008478 |
| Shcbp1 | 1.855895335 | 2.40E-20 | 1.37E-17 | Zfp112 | -1.58611 | 9.00E-05 | 0.001699 |
| Dyrk4 | 1.855290818 | 0.0008987 | 0.010125 | Retnlg | -1.5945 | 6.98E-10 | 8.28E-08 |
| B3gnt3 | 1.853951378 | 6.76E-06 | 0.000215 | Dkk2 | -1.60241 | 5.32E-06 | 0.000176 |
| Fcna | 1.853896563 | 1.39E-06 | 5.99E-05 | H2-M2 | -1.64351 | 0.001001 | 0.010988 |
| Dyrk3 | 1.852716192 | 0.0052625 | 0.037962 | Gm9403 | -1.66498 | 0.007655 | 0.049524 |
| Nusap1 | 1.848140005 | 1.08E-19 | 5.92E-17 | Sirpb1a | -1.68324 | 0.006922 | 0.045889 |
| Camp | 1.840405038 | 0.0033047 | 0.026906 | Jag1 | -1.70583 | 0.000606 | 0.007372 |
| Chad | 1.838807582 | 7.99E-05 | 0.001547 | 9530046B1 | -1.73165 | 0.007572 | 0.049147 |
| Fbln1 | 1.831906486 | 2.20E-08 | 1.72E-06 | Cd79a | -1.7565 | 0.003928 | 0.030584 |
| Kif4 | 1.829142553 | 1.23E-09 | 1.35E-07 | Gm26885 | -1.81348 | 0.001049 | 0.01131 |
| Orm1 | 1.827384593 | 0.003993 | 0.030987 | H2-Aa | -1.82641 | 3.32E-63 | 4.73E-59 |
| Ifit1bl1 | 1.818849639 | 5.67E-07 | 2.79E-05 | Klra2 | -1.83375 | 1.03E-05 | 0.000301 |
| Hyal5 | 1.803158995 | 1.41E-05 | 0.000389 | Ighj4 | -1.84805 | 7.61E-05 | 0.001485 |
| Cd244 | 1.799734429 | 9.61E-06 | 0.000285 | Tenm4 | -1.87469 | 0.002867 | 0.024075 |
| Slco4c1 | 1.798455533 | 0.0044905 | 0.033977 | H2-Eb1 | -1.89048 | 1.09E-45 | 5.18E-42 |
| Ccnb1 | 1.795628508 | 4.20E-34 | 7.47E-31 | Vcan | -1.89181 | 0.003568 | 0.028498 |
| Gm42047 | 1.79314848 | 1.76E-12 | 3.34E-10 | Tmem150b | -1.9625 | 1.40E-06 | 6.04E-05 |
| Fam83d | 1.788337271 | 2.07E-06 | 8.28E-05 | Clec4e | -1.96626 | 5.16E-06 | 0.000173 |
| Tpx2 | 1.783885966 | 2.62E-22 | 1.86E-19 | Cd300e | -2.00918 | 0.002318 | 0.020678 |
| Gm8817 | 1.782096514 | 0.0010083 | 0.011037 | Adgre4 | -2.04873 | 2.01E-08 | 1.60E-06 |
| Pbk | 1.782020534 | 1.53E-26 | 1.45E-23 | Napsa | -2.05879 | 3.19E-08 | 2.35E-06 |
| Ccnd2 | 1.779889379 | 0.0035345 | 0.028359 | Sirpb1b | -2.10454 | 1.77E-05 | 0.000471 |
| Hmcn2 | 1.779727326 | 2.75E-06 | 0.000105 | Vnn3 | -2.10626 | 0.000398 | 0.005397 |
| Prr11 | 1.77506348 | 1.77E-06 | 7.33E-05 | Pdlim1 | -2.1234 | 8.80E-05 | 0.001672 |
| Gm47283 | 1.767310142 | 1.06E-18 | 4.72E-16 | Postn | -2.14317 | 0.000154 | 0.002565 |
| Kif18b | 1.761646129 | 4.57E-13 | 9.71E-11 | Serpinb10 | -2.21061 | 0.000176 | 0.002857 |
| Rhof | 1.76001644 | 4.42E-07 | 2.25E-05 | Gm14548 | -2.2148 | 0.000311 | 0.004463 |
| Sh2d1b1 | 1.751996181 | 8.10E-06 | 0.000246 | F10 | -2.25283 | 0.000133 | 0.002312 |
| Ccna2 | 1.750558442 | 1.30E-30 | 1.85E-27 | Asgr2 | -2.25417 | 0.00021 | 0.003288 |
| Cdca3 | 1.726875521 | 5.97E-29 | 6.08E-26 | Ighg2b | -2.2661 | 0.000188 | 0.003021 |
| Lockd | 1.726326129 | 5.43E-10 | 6.60E-08 | Treml4 | -2.34192 | 2.69E-05 | 0.00066 |
| Clspn | 1.722773979 | 1.19E-07 | 7.60E-06 | Sirpb1c | -2.34286 | 1.66E-07 | 9.99E-06 |
| Mastl | 1.722596385 | 2.96E-08 | 2.21E-06 | Grik2 | -2.36215 | 0.000113 | 0.002042 |
| Chrna1os | 1.721516078 | 0.0002493 | 0.003757 | Itgb7 | -2.44038 | 8.64E-10 | 9.84E-08 |
| Brip1 | 1.721374599 | 2.65E-10 | 3.50E-08 | Lilra6 | -2.47595 | 7.12E-07 | 3.40E-05 |
| Pimreg | 1.715051753 | 2.12E-19 | 1.12E-16 | Ighj3 | -2.48709 | 3.73E-05 | 0.000861 |
| Neil3 | 1.713380431 | 1.25E-09 | 1.36E-07 | Gpr141 | -2.50568 | 2.43E-09 | 2.42E-07 |
| Psrc1 | 1.705804069 | 8.11E-07 | 3.73E-05 | Ighj2 | -2.56949 | 2.36E-08 | 1.83E-06 |
| Birc5 | 1.694797347 | 1.64E-24 | 1.38E-21 | Gm9733 | -2.61431 | 3.66E-10 | 4.74E-08 |
| Depdc1b | 1.693968254 | 4.72E-06 | 0.000161 | Ear2 | -3.12022 | 2.85E-09 | 2.74E-07 |
| Psat1 | 1.693813729 | 5.29E-08 | 3.73E-06 | Epx | -3.97102 | 5.02E-07 | 2.50E-05 |
| Cenpm | 1.693720957 | 1.73E-07 | 1.03E-05 | Prg2 | -4.33371 | 5.29E-05 | 0.001128 |
| Ctla2a | 1.687755159 | 2.38E-19 | 1.13E-16 | Jchain | -4.50965 | 5.29E-12 | 9.31E-10 |
| Nuf2 | 1.683872 | 2.28E-17 | 8.76E-15 | Ear6 | -4.91549 | 0.002679 | 0.022942 |
| Kif23 | 1.6836914 | 3.54E-13 | 7.88E-11 | Igkc | -7.31439 | 5.98E-16 | 2.08E-13 |
| Troap | 1.681410652 | 1.36E-07 | 8.43E-06 | Igha | -10.4455 | 1.85E-11 | 2.96E-09 |

| | | | |
|---|---|---|---|
| Kif11 | 1.681098132 | 6.83E-19 | 3.14E-16 |
| Phf19 | 1.679601439 | 0.0011058 | 0.011734 |
| Fn1 | 1.673059748 | 1.31E-05 | 0.000367 |
| Rrm2 | 1.672654939 | 3.73E-30 | 4.09E-27 |
| Knl1 | 1.667099994 | 4.37E-22 | 2.97E-19 |
| E2f7 | 1.66663028 | 6.09E-05 | 0.001267 |
| Ska3 | 1.665933488 | 2.85E-07 | 1.55E-05 |
| Kifc1 | 1.66251415 | 0.0001382 | 0.002368 |
| Gbp8 | 1.661493933 | 0.0001922 | 0.003072 |
| Scml4 | 1.660291978 | 1.67E-06 | 7.00E-05 |
| Cenpn | 1.655145958 | 2.43E-11 | 3.80E-09 |
| Hmmr | 1.650127277 | 1.76E-20 | 1.05E-17 |
| Ccl2 | 1.638627092 | 2.37E-30 | 2.81E-27 |
| Ccne1 | 1.638369566 | 2.17E-09 | 2.19E-07 |
| Ncapg | 1.635791572 | 1.68E-14 | 4.78E-12 |
| Oip5 | 1.634351357 | 2.08E-09 | 2.12E-07 |
| Ccnb2 | 1.634215041 | 8.08E-31 | 1.28E-27 |
| Mxd3 | 1.633739337 | 1.58E-05 | 0.000425 |
| Anln | 1.632725124 | 8.51E-12 | 1.46E-09 |
| Mcm10 | 1.625576878 | 1.26E-07 | 7.98E-06 |
| Hcar2 | 1.622454746 | 2.52E-05 | 0.000624 |
| Mtfr2 | 1.617442058 | 0.0004166 | 0.005566 |
| mt-Tn | 1.616804081 | 1.35E-09 | 1.43E-07 |
| Spag5 | 1.616322437 | 2.47E-09 | 2.43E-07 |
| Atp10a | 1.615246479 | 8.56E-10 | 9.83E-08 |
| Rbm44 | 1.614253922 | 0.0020427 | 0.018766 |
| Ckap2 | 1.61396572 | 2.04E-13 | 4.62E-11 |
| Foxr1 | 1.61292783 | 0.0027812 | 0.023518 |
| Gna14 | 1.607405774 | 2.07E-09 | 2.12E-07 |
| Map3k19 | 1.60698138 | 0.000181 | 0.002929 |
| Stap2 | 1.605367151 | 0.0042227 | 0.032363 |
| Gm17201 | 1.60156926 | 0.0044057 | 0.033527 |
| Ankle1 | 1.599684263 | 1.29E-11 | 2.12E-09 |
| Cdca8 | 1.595558426 | 2.36E-19 | 1.13E-16 |
| Gck | 1.593421626 | 0.0018106 | 0.017098 |
| Cdc25c | 1.592317443 | 4.40E-06 | 0.000153 |
| Tmem154 | 1.589673845 | 0.00012 | 0.002138 |
| Top2a | 1.587121747 | 2.34E-30 | 2.81E-27 |
| Wfdc21 | 1.586075182 | 3.63E-06 | 0.000131 |
| Cdca2 | 1.583371032 | 7.06E-16 | 2.39E-13 |
| Kif20a | 1.578108578 | 5.99E-13 | 1.24E-10 |
| Ska1 | 1.574899403 | 0.0001005 | 0.001853 |
| Hist1h4h | 1.573753287 | 1.63E-07 | 9.87E-06 |
| Spc24 | 1.572705002 | 2.78E-12 | 5.08E-10 |
| Cenpf | 1.5713398 | 4.75E-23 | 3.76E-20 |
| Ehd1 | 1.569907702 | 0.0015788 | 0.015399 |
| Dtl | 1.568628063 | 1.02E-13 | 2.51E-11 |

| | | | |
|---|---|---|---|
| E2f8 | 1.563959289 | 3.05E-05 | 0.000729 |
| Mcoln3 | 1.562138651 | 2.78E-06 | 0.000106 |
| Plk1 | 1.558817128 | 7.65E-18 | 3.20E-15 |
| Pdcd1lg2 | 1.557618336 | 0.0076054 | 0.049295 |
| 1700001C19Rik | 1.554533771 | 0.000287 | 0.004175 |
| Aurka | 1.549734907 | 3.80E-14 | 1.04E-11 |
| Rad51ap1 | 1.549180029 | 1.17E-14 | 3.39E-12 |
| Melk | 1.546663123 | 5.31E-11 | 7.80E-09 |
| Dusp27 | 1.546588998 | 0.001419 | 0.014241 |
| Esco2 | 1.539868221 | 3.93E-07 | 2.04E-05 |
| Rad51 | 1.535754287 | 7.82E-14 | 1.95E-11 |
| Htr2b | 1.530124733 | 1.36E-10 | 1.89E-08 |
| Cped1 | 1.520628553 | 1.81E-08 | 1.48E-06 |
| Rad54b | 1.517591035 | 6.92E-06 | 0.000218 |
| Phlda3 | 1.516083223 | 0.0001366 | 0.002349 |
| Mms22l | 1.509792272 | 4.45E-11 | 6.67E-09 |
| Xlr4a | 1.509356141 | 0.0016592 | 0.016032 |
| Tmem51 | 1.507157708 | 2.68E-07 | 1.47E-05 |
| Ptpn7 | 1.506251545 | 1.21E-08 | 1.03E-06 |
| Itgb3 | 1.505112847 | 0.0001317 | 0.002287 |
| Pdlim7 | 1.498090091 | 0.0001372 | 0.002357 |
| Iqcg | 1.4945193 | 0.0001634 | 0.002678 |
| Cfb | 1.486812402 | 0.0005115 | 0.006486 |
| Cep55 | 1.484119763 | 7.86E-13 | 1.51E-10 |
| Spc25 | 1.48387218 | 1.66E-15 | 5.37E-13 |
| Prc1 | 1.480010034 | 3.91E-18 | 1.69E-15 |
| Adamtsl2 | 1.471541132 | 0.0003546 | 0.004945 |
| Sgo1 | 1.468122327 | 1.80E-07 | 1.07E-05 |
| Ccne2 | 1.467380561 | 1.76E-13 | 4.03E-11 |
| C130012C08Rik | 1.463133221 | 0.0023978 | 0.021155 |
| Acp5 | 1.446808483 | 6.89E-05 | 0.001378 |
| Pvrig | 1.438176263 | 5.88E-08 | 4.10E-06 |
| Gm27008 | 1.436588236 | 2.02E-05 | 0.000522 |
| Ddias | 1.432587341 | 7.64E-05 | 0.001487 |
| Ctf2 | 1.431834246 | 0.0009021 | 0.010147 |
| Mns1 | 1.428360834 | 0.0014365 | 0.014355 |
| S100a8 | 1.427547817 | 0.0073425 | 0.04794 |
| Kif2c | 1.426829947 | 5.11E-13 | 1.07E-10 |
| Cdc20 | 1.425494987 | 1.45E-21 | 9.01E-19 |
| Gm11223 | 1.421935933 | 8.21E-06 | 0.000249 |
| Pclaf | 1.421460752 | 7.90E-18 | 3.21E-15 |
| Bub1b | 1.420703704 | 1.81E-15 | 5.61E-13 |
| Cenpe | 1.41911404 | 1.09E-17 | 4.33E-15 |
| Tmem2 | 1.41873806 | 3.15E-07 | 1.68E-05 |
| Ect2 | 1.417176689 | 4.41E-13 | 9.51E-11 |
| St8sia1 | 1.410515965 | 8.75E-09 | 7.72E-07 |
| Gm9999 | 1.402627982 | 2.04E-06 | 8.22E-05 |

| | | | |
|---|---|---|---|
| Adcy3 | 1.395515077 | 5.50E-12 | 9.55E-10 |
| Ttk | 1.39394371 | 6.48E-07 | 3.13E-05 |
| Ndc80 | 1.384027126 | 1.38E-13 | 3.23E-11 |
| Sdk1 | 1.383565825 | 0.0026957 | 0.023009 |
| Racgap1 | 1.380773854 | 5.79E-22 | 3.75E-19 |
| Itpr3 | 1.379659098 | 3.76E-07 | 1.97E-05 |
| Cenph | 1.379103332 | 7.86E-06 | 0.000241 |
| Cxcl1 | 1.367901166 | 0.0014262 | 0.014293 |
| Xlr4c | 1.362603808 | 0.0003937 | 0.005369 |
| 4930479D17Rik | 1.359732583 | 0.0004525 | 0.005933 |
| Xcr1 | 1.358455023 | 0.0001207 | 0.002138 |
| Cit | 1.353289416 | 1.80E-08 | 1.48E-06 |
| Bcl11a | 1.352767362 | 0.0015211 | 0.015 |
| Trem3 | 1.349104505 | 6.61E-05 | 0.001343 |
| Foxm1 | 1.347777505 | 5.15E-08 | 3.66E-06 |
| Abca13 | 1.34499482 | 6.06E-05 | 0.001265 |
| Ccnf | 1.339692847 | 1.51E-10 | 2.09E-08 |
| Mmd | 1.335957659 | 7.89E-09 | 7.11E-07 |
| Serpinb1c | 1.334536546 | 0.006635 | 0.044507 |
| Cenpk | 1.333085957 | 7.25E-07 | 3.44E-05 |
| mt-Ti | 1.325635956 | 0.0037822 | 0.029756 |
| Zwilch | 1.321790883 | 4.17E-06 | 0.000147 |
| Sgol2a | 1.316137821 | 1.03E-05 | 0.000302 |
| Nrap | 1.31536755 | 9.00E-06 | 0.000269 |
| Pstpip1 | 1.311306211 | 4.28E-05 | 0.000953 |
| Myl2 | 1.311031489 | 2.78E-06 | 0.000106 |
| Kif22 | 1.306521683 | 2.35E-17 | 8.80E-15 |
| Hist1h2bc | 1.300158162 | 0.0002104 | 0.003293 |
| Trpm6 | 1.299453485 | 7.82E-05 | 0.00152 |
| 4930517O19Rik | 1.299244649 | 3.59E-05 | 0.000831 |
| Arhgef7 | 1.299127426 | 5.05E-11 | 7.48E-09 |
| Pcp4 | 1.293907282 | 5.37E-10 | 6.59E-08 |
| Slc38a10 | 1.292584893 | 0.000859 | 0.009762 |
| Ifi30 | 1.292099762 | 0.0004133 | 0.005547 |
| Spint1 | 1.284860611 | 0.0001632 | 0.002678 |
| Gper1 | 1.284823717 | 2.44E-05 | 0.000606 |
| 1700020N01Rik | 1.281728656 | 2.93E-08 | 2.21E-06 |
| Knstrn | 1.281647609 | 6.46E-13 | 1.31E-10 |
| Nedd9 | 1.280849092 | 2.76E-07 | 1.51E-05 |
| Diaph3 | 1.275993895 | 0.0001134 | 0.002055 |
| Cst7 | 1.27588802 | 1.81E-15 | 5.61E-13 |
| mt-Tc | 1.269818911 | 5.54E-05 | 0.001173 |
| Cdkn3 | 1.267605232 | 2.66E-09 | 2.58E-07 |
| Oas3 | 1.267093606 | 0.0010749 | 0.011544 |
| Tcim | 1.266638165 | 1.63E-09 | 1.71E-07 |
| Rab3b | 1.256106346 | 0.0040914 | 0.031585 |
| Wee1 | 1.254671689 | 7.47E-07 | 3.52E-05 |

| | | | |
|---|---|---|---|
| Scarb1 | 1.247716621 | 6.59E-10 | 7.89E-08 |
| Hlf | 1.246320888 | 0.0008116 | 0.00932 |
| Itgb2l | 1.239417011 | 0.0064599 | 0.043867 |
| Aurkb | 1.238712288 | 4.57E-15 | 1.36E-12 |
| Kynu | 1.238445985 | 9.61E-05 | 0.001787 |
| Stmn1 | 1.236865846 | 7.33E-43 | 1.74E-39 |
| Cdc6 | 1.228851021 | 3.91E-05 | 0.00089 |
| Mybph | 1.228568537 | 0.000275 | 0.004058 |
| Cdk1 | 1.22769681 | 6.86E-26 | 6.11E-23 |
| Gpsm2 | 1.223216206 | 0.0001423 | 0.002425 |
| Ass1 | 1.221177947 | 4.94E-06 | 0.000167 |
| Ckap2l | 1.2088665 | 5.05E-10 | 6.26E-08 |
| Trim59 | 1.206358853 | 3.73E-10 | 4.79E-08 |
| Shisa7 | 1.199285183 | 0.0011404 | 0.011969 |
| Dynlt1a | 1.188513341 | 1.05E-05 | 0.000306 |
| Coro2a | 1.187558869 | 4.18E-07 | 2.13E-05 |
| Cd5 | 1.182576895 | 4.23E-05 | 0.000946 |
| Cenpi | 1.180638123 | 1.36E-06 | 5.89E-05 |
| Cdca5 | 1.174886295 | 3.04E-07 | 1.63E-05 |
| Pglyrp1 | 1.171952895 | 8.83E-05 | 0.001676 |
| Ccl12 | 1.169292375 | 1.31E-11 | 2.12E-09 |
| Ly6g | 1.165506688 | 0.0067973 | 0.045297 |
| Gm6749 | 1.160118928 | 0.0004956 | 0.006329 |
| 4930486L24Rik | 1.159506605 | 2.59E-06 | 0.0001 |
| Gm5431 | 1.152408128 | 0.0011286 | 0.011896 |
| Rtkn2 | 1.152296422 | 0.002558 | 0.022184 |
| Brca1 | 1.149757615 | 0.0004077 | 0.005477 |
| Cspg4 | 1.145919498 | 4.64E-10 | 5.89E-08 |
| Gtse1 | 1.144742926 | 0.0012986 | 0.013306 |
| Rgs16 | 1.144002432 | 0.0001169 | 0.002099 |
| C330027C09Rik | 1.141196592 | 1.93E-07 | 1.13E-05 |
| Cdkn2b | 1.141073281 | 0.0009477 | 0.01052 |
| Mybl2 | 1.138895553 | 0.0054116 | 0.038744 |
| Cks1b | 1.1385104 | 2.32E-19 | 1.13E-16 |
| Chaf1a | 1.137417841 | 3.29E-05 | 0.000776 |
| mt-Tp | 1.132949788 | 2.56E-11 | 3.93E-09 |
| Kif20b | 1.129509369 | 1.01E-09 | 1.14E-07 |
| Btla | 1.129217563 | 0.0005889 | 0.007216 |
| Mis18bp1 | 1.129141532 | 2.01E-08 | 1.60E-06 |
| Smtn | 1.128152367 | 0.0005852 | 0.00719 |
| Cenpw | 1.127932711 | 3.62E-06 | 0.000131 |
| Qpct | 1.124453054 | 1.14E-13 | 2.76E-11 |
| Pkmyt1 | 1.122080541 | 9.63E-07 | 4.35E-05 |
| C130050O18Rik | 1.119866577 | 2.22E-07 | 1.26E-05 |
| Net1 | 1.115920926 | 0.0004298 | 0.005715 |
| Gdpd5 | 1.115843176 | 5.21E-06 | 0.000174 |
| Gcnt2 | 1.114424824 | 4.34E-09 | 4.04E-07 |

| | | | |
|---|---|---|---|
| Fam89a | 1.114345395 | 1.99E-06 | 8.08E-05 |
| Wwtr1 | 1.111572889 | 0.0050682 | 0.037008 |
| Eldr | 1.098978306 | 0.0003621 | 0.005036 |
| Agap2 | 1.098720514 | 0.0010208 | 0.011122 |
| Rad51c | 1.098297093 | 0.0050744 | 0.037018 |
| Esr1 | 1.097973158 | 2.09E-07 | 1.20E-05 |
| Ticrr | 1.088623042 | 0.0058473 | 0.040917 |
| Gm15928 | 1.081684055 | 0.0055982 | 0.039678 |
| Hspa12a | 1.079893767 | 0.0004542 | 0.005945 |
| Rgs7bp | 1.065967181 | 4.37E-05 | 0.000972 |
| Otub2 | 1.063327769 | 0.000153 | 0.002552 |
| Pear1 | 1.062240302 | 8.82E-09 | 7.72E-07 |
| Hist1h1c | 1.05973206 | 4.64E-07 | 2.34E-05 |
| Rab19 | 1.057148526 | 0.0013206 | 0.0135 |
| Ppm1j | 1.055586959 | 0.0015943 | 0.015528 |
| Igsf8 | 1.05250813 | 0.0055337 | 0.039302 |
| Gm26902 | 1.042668839 | 2.52E-07 | 1.41E-05 |
| Dock9 | 1.034359545 | 4.04E-07 | 2.08E-05 |
| Cldn12 | 1.033165256 | 0.0002196 | 0.003393 |
| Smc4 | 1.031678754 | 3.44E-17 | 1.26E-14 |
| Klrb1b | 1.031056874 | 6.61E-06 | 0.000212 |
| Plpp1 | 1.029938993 | 2.46E-09 | 2.43E-07 |
| Hist1h4i | 1.026439811 | 6.08E-05 | 0.001267 |
| Gm16897 | 1.025825603 | 7.83E-05 | 0.00152 |
| Gm16894 | 1.020091171 | 0.003548 | 0.028432 |
| 2310022B05Rik | 1.019937224 | 2.91E-06 | 0.000109 |
| Masp1 | 1.019306498 | 9.89E-07 | 4.44E-05 |
| Plp2 | 1.010415149 | 2.81E-06 | 0.000107 |
| Tnfsf8 | 1.010094218 | 7.89E-08 | 5.28E-06 |
| Cd300lf | 1.00905645 | 2.61E-08 | 1.98E-06 |
| Trip13 | 1.006380096 | 8.16E-05 | 0.001571 |
| Serf1 | 1.003037138 | 1.52E-06 | 6.41E-05 |
