## Supplementary File 5 for "Dicer deficiency in microglia leads to accelerated demyelination and failed remyelination"

**Supplementary file 5:** List of enriched biological processed of DEGs from flow sorted microglia isolated from Cx3cr1$^{creERT2}$ Cre- control and Cre+ mutant animals fed to CUP diet for 6 weeks.

| Category | Term | Count | % | PValue | Genes | List Total | Pop Hits | Pop Total | Fold Enrich | Bonferroni | Benjamini | FDR |
|---|---|---|---|---|---|---|---|---|---|---|---|---|
| GOTERM_ | GO:0007049~cell cycle | 77 | 16.70282 | 3.50E-38 | 240641, 26 | 369 | 614 | 18082 | 6.145291 | 5.95E-35 | 5.95E-35 | 5.86E-35 |
| GOTERM_ | GO:0051301~cell division | 62 | 13.44902 | 1.63E-37 | 240641, 26 | 369 | 374 | 18082 | 8.123444 | 2.78E-34 | 1.39E-34 | 1.37E-34 |
| GOTERM_ | GO:0007067~mitotic nuclear division | 54 | 11.71367 | 2.90E-36 | 240641, 26 | 369 | 277 | 18082 | 9.552875 | 4.93E-33 | 1.64E-33 | 1.62E-33 |
| GOTERM_ | GO:0007059~chromosome segregation | 23 | 4.989154 | 3.61E-18 | 66442, 66 | 369 | 89 | 18082 | 12.66362 | 6.14E-15 | 1.53E-15 | 1.51E-15 |
| GOTERM_ | GO:0002376~immune system process | 30 | 6.507592 | 1.35E-09 | 246727, 14 | 369 | 383 | 18082 | 3.838332 | 2.29E-06 | 4.58E-07 | 4.51E-07 |
| GOTERM_ | GO:0000281~mitotic cytokinesis | 10 | 2.169197 | 5.33E-09 | 16571, 19 | 369 | 30 | 18082 | 16.33424 | 9.06E-06 | 1.51E-06 | 1.49E-06 |
| GOTERM_ | GO:0000070~mitotic sister chromatid segregation | 9 | 1.952278 | 1.02E-08 | 12704, 18 | 369 | 23 | 18082 | 19.17497 | 1.74E-05 | 2.49E-06 | 2.45E-06 |
| GOTERM_ | GO:0007080~mitotic metaphase plate congression | 9 | 1.952278 | 3.12E-07 | 56742, 22 | 369 | 34 | 18082 | 12.97131 | 5.30E-04 | 6.62E-05 | 6.52E-05 |
| GOTERM_ | GO:0007052~mitotic spindle organization | 8 | 1.735358 | 1.12E-06 | 66442, 76 | 369 | 28 | 18082 | 14.00077 | 0.001899 | 2.11E-04 | 2.08E-04 |
| GOTERM_ | GO:0051310~metaphase plate congression | 6 | 1.301518 | 2.39E-06 | 229841, 7 | 369 | 12 | 18082 | 24.50136 | 0.004057 | 4.07E-04 | 4.00E-04 |
| GOTERM_ | GO:0032467~positive regulation of cytokinesis | 8 | 1.735358 | 3.70E-06 | 12704, 24 | 369 | 33 | 18082 | 11.87944 | 0.006263 | 5.62E-04 | 5.53E-04 |
| GOTERM_ | GO:0007018~microtubule-based movement | 11 | 2.386117 | 3.97E-06 | 240641, 22 | 369 | 78 | 18082 | 6.910639 | 0.00672 | 5.62E-04 | 5.53E-04 |
| GOTERM_ | GO:0051988~regulation of attachment of spindle microtubules to kinetochore | 5 | 1.084599 | 5.63E-06 | 18005, 13 | 369 | 7 | 18082 | 35.00194 | 0.009517 | 7.36E-04 | 7.24E-04 |
| GOTERM_ | GO:0000910~cytokinesis | 8 | 1.735358 | 8.31E-06 | 12704, 23 | 369 | 37 | 18082 | 10.59518 | 0.014018 | 0.001008 | 9.93E-04 |
| GOTERM_ | GO:0007051~spindle organization | 6 | 1.301518 | 1.23E-05 | 16551, 54 | 369 | 16 | 18082 | 18.37602 | 0.020744 | 0.001397 | 0.001376 |
| GOTERM_ | GO:0034501~protein localization to kinetochore | 5 | 1.084599 | 3.22E-05 | 76464, 22 | 369 | 10 | 18082 | 24.50136 | 0.053187 | 0.003416 | 0.003364 |
| GOTERM_ | GO:0006955~immune response | 18 | 3.904555 | 4.52E-05 | 246727, 1 | 369 | 272 | 18082 | 3.242826 | 0.073964 | 0.00448 | 0.004412 |
| GOTERM_ | GO:0000086~G2/M transition of mitotic cell cycle | 7 | 1.518438 | 4.75E-05 | 12704, 12 | 369 | 33 | 18082 | 10.39451 | 0.077481 | 0.00448 | 0.004412 |
| GOTERM_ | GO:0090307~mitotic spindle assembly | 7 | 1.518438 | 6.72E-05 | 72119, 18 | 369 | 35 | 18082 | 9.800542 | 0.107925 | 0.006011 | 0.005919 |
| GOTERM_ | GO:0019886~antigen processing and presentation of exogenous peptide antigen via MHC class II | 5 | 1.084599 | 1.44E-04 | 14969, 65 | 369 | 14 | 18082 | 17.50097 | 0.216686 | 0.01221 | 0.012023 |
| GOTERM_ | GO:0051256~mitotic spindle midzone assembly | 4 | 0.867679 | 1.60E-04 | 16571, 71 | 369 | 6 | 18082 | 32.66847 | 0.237744 | 0.012926 | 0.012728 |
| GOTERM_ | GO:0045087~innate immune response | 21 | 4.555315 | 2.14E-04 | 246727, 26 | 369 | 400 | 18082 | 2.572642 | 0.304948 | 0.016533 | 0.01628 |
| GOTERM_ | GO:0006468~protein phosphorylation | 26 | 5.639913 | 3.28E-04 | 268697, 26 | 369 | 576 | 18082 | 2.211928 | 0.427158 | 0.02422 | 0.023849 |
| GOTERM_ | GO:0006974~cellular response to DNA damage stimulus | 21 | 4.555315 | 4.09E-04 | 21973, 20 | 369 | 420 | 18082 | 2.450136 | 0.500659 | 0.02893 | 0.028487 |
| GOTERM_ | GO:0032147~activation of protein kinase activity | 6 | 1.301518 | 5.04E-04 | 72119, 13 | 369 | 33 | 18082 | 8.909584 | 0.575685 | 0.034283 | 0.033758 |
| GOTERM_ | GO:0030071~regulation of mitotic metaphase/anaphase transition | 4 | 0.867679 | 6.41E-04 | 23834, 22 | 369 | 9 | 18082 | 21.77898 | 0.66366 | 0.040344 | 0.039726 |
| GOTERM_ | GO:0008608~attachment of spindle microtubules to kinetochore | 4 | 0.867679 | 6.41E-04 | 229841, 7 | 369 | 9 | 18082 | 21.77898 | 0.66366 | 0.040344 | 0.039726 |
| GOTERM_ | GO:0006935~chemotaxis | 10 | 2.169197 | 6.92E-04 | 20201, 20 | 369 | 118 | 18082 | 4.152772 | 0.691518 | 0.041989 | 0.041346 |
| GOTERM_ | GO:0006954~inflammatory response | 18 | 3.904555 | 7.31E-04 | 20201, 20 | 369 | 344 | 18082 | 2.564095 | 0.711237 | 0.042817 | 0.042162 |
