## Supplementary File 6 for "Dicer deficiency in microglia leads to accelerated demyelination and failed remyelination"

**Supplementary file 6:** Ingenuity Canonical Pathways (IPA) analysis of DEGs from flow sorted microglia isolated from Cx3cr1$^{creERT2}$ Cre- control and Cre+ mutant animals fed to CUP diet for 6 weeks.

| | | | | |
|---|---|---|---|---|
| Kinetochore Metaphase Signaling Pathway | 24.3 | 0.267 | 3.128 | AURKB,BIRC5,BUB1,BUB1B,CCNB1,CDC20,CDCA8,CDK1,CENPE,CENPH,CENPK,CENPN,CENPW,KIF2C,KNL1,MASTL,MXD3,NDC80, |
| Estrogen-mediated S-phase Entry | 6.72 | 0.269 | 2.646 | CCNA2,CCNE1,CCNE2,CDK1,E2F7,E2F8,ESR1 |
| Mitotic Roles of Polo-Like Kinase | 8.98 | 0.182 | 2.333 | CCNB1,CCNB2,CDC20,CDC25C,CDK1,KIF11,KIF23,PKMYT1,PLK1,PPM1J,PRC1,WEE1 |
| Pyridoxal 5'-phosphate Salvage Pathway | 2.28 | 0.0758 | 2.236 | CDK1,NEK2,PLK1,PRKCQ,TTK |
| Salvage Pathways of Pyrimidine Ribonucleotides | 1.59 | 0.051 | 2.236 | CDK1,NEK2,PLK1,PRKCQ,TTK |
| cAMP-mediated signaling | 0.461 | 0.0219 | 2.236 | ADCY3,ADORA1,GPER1,HCAR2,XCR1 |
| Cell Cycle Regulation by BTG Family Proteins | 3.42 | 0.135 | 2 | CCNE1,CCNE2,E2F7,E2F8,PPM1J |
| RhoA Signaling | 1.23 | 0.0407 | 2 | ANLN,CIT,DLC1,MYL2,RND3 |
| GPCR-Mediated Nutrient Sensing in Enteroendocrine Cells | 0.91 | 0.0357 | 2 | ADCY3,GNA14,ITPR3,PRKCQ |
| White Adipose Tissue Browning Pathway | 0.752 | 0.031 | 2 | ADCY3,CAMP,RARB,RXRG |
| Dopamine-DARPP32 Feedback in cAMP Signaling | 0.523 | 0.0245 | 2 | ADCY3,ITPR3,PPM1J,PRKCQ |
| Insulin Secretion Signaling Pathway | 0.402 | 0.0206 | 2 | ADCY3,GCK,GNA14,ITPR3,PRKCQ |
| Estrogen Receptor Signaling | 0.489 | 0.0213 | 1.89 | ADCY3,ESR1,GNA14,GNB3,GPER1,MYL2,PRKCQ |
| Cyclins and Cell Cycle Regulation | 7.92 | 0.148 | 1.732 | CCNA2,CCNB1,CCNB2,CCND2,CCNE1,CCNE2,CDK1,CDKN2B,E2F7,E2F8,PPM1J,WEE1 |
| Adrenomedullin signaling pathway | 0.921 | 0.0305 | 1.633 | ADCY3,GNA14,ITPR3,KCNN4,MATK,RXRG |
| GNRH Signaling | 0.764 | 0.0289 | 1.342 | ADCY3,GNA14,GNB3,ITPR3,PRKCQ |
| Hepatic Fibrosis Signaling Pathway | 1.01 | 0.0272 | 1.265 | CCL2,CD40,CYBB,IL1R2,ITGA4,ITGB3,MYL2,PRKCQ,RHOF,RND3 |
| Aryl Hydrocarbon Receptor Signaling | 3.11 | 0.0629 | 1 | AHR,AHRR,CCNA2,CCND2,CCNE1,CCNE2,ESR1,RARB,RXRG |
| T Cell Exhaustion Signaling Pathway | 1.52 | 0.04 | 1 | BTLA,HLA-DQA1,HLA-DQB1,HLA-DRB5,PDCD1LG2,PPM1J,PRKCQ |
| PD-1, PD-L1 cancer immunotherapy pathway | 1.46 | 0.0472 | 1 | HLA-DQA1,HLA-DQB1,HLA-DRB5,PDCD1LG2,PRKCQ |
| Glioma Invasiveness Signaling | 1.43 | 0.0541 | 1 | HMMR,ITGB3,RHOF,RND3 |
| Gαs Signaling | 0.963 | 0.0374 | 1 | ADCY3,GNB3,GPER1,HCAR2 |
| CREB Signaling in Neurons | 0.848 | 0.029 | 1 | ADCY3,GNA14,GNB3,GRIK2,ITPR3,PRKCQ |
| Cholecystokinin/Gastrin-mediated Signaling | 0.839 | 0.0336 | 1 | ITPR3,PRKCQ,RHOF,RND3 |
| Semaphorin Neuronal Repulsive Signaling Pathway | 0.745 | 0.0308 | 1 | CSPG4,ITGA4,MYL2,VCAN |
| Cardiac Hypertrophy Signaling (Enhanced) | 0.682 | 0.0226 | 1 | ADCY3,CYBB,DIAPH3,GNA14,GNB3,IL1R2,ITGA4,ITPR3,LTB,PRKCQ,TNFSF8 |
| Huntington's Disease Signaling | 0.666 | 0.0253 | 1 | GNA14,GNB3,HSPA1A/HSPA1B,Hspa1b,PRKCQ,STX1A |
| D-myo-inositol (1,4,5,6)-Tetrakisphosphate Biosynthesis | 0.654 | 0.0282 | 1 | CDC25C,DUSP27,PTPN22,PTPN7 |
| D-myo-inositol (3,4,5,6)-tetrakisphosphate Biosynthesis | 0.654 | 0.0282 | 1 | CDC25C,DUSP27,PTPN22,PTPN7 |
| 3-phosphoinositide Degradation | 0.562 | 0.0256 | 1 | CDC25C,DUSP27,PTPN22,PTPN7 |
| D-myo-inositol-5-phosphate Metabolism | 0.558 | 0.0255 | 1 | CDC25C,DUSP27,PTPN22,PTPN7 |
| 3-phosphoinositide Biosynthesis | 0.506 | 0.0241 | 1 | CDC25C,DUSP27,PTPN22,PTPN7 |
| NF-κB Signaling | 0.441 | 0.0223 | 1 | CD40,IL1R2,PRKCQ,TLR8 |
| Endothelin-1 Signaling | 0.402 | 0.0213 | 1 | ADCY3,GNA14,ITPR3,PRKCQ |
| Synaptic Long Term Depression | 0.398 | 0.0212 | 1 | GNA14,ITPR3,PPM1J,PRKCQ |
| Superpathway of Inositol Phosphate Compounds | 0.359 | 0.0201 | 1 | CDC25C,DUSP27,PTPN22,PTPN7 |
| Role of NFAT in Cardiac Hypertrophy | 0.306 | 0.0187 | 1 | ADCY3,GNB3,ITPR3,PRKCQ |
| PKCθ Signaling in T Lymphocytes | 0.569 | 0.0258 | -1 | HLA-DQA1,HLA-DQB1,HLA-DRB5,PRKCQ |
| Wnt/β-catenin Signaling | 0.471 | 0.0231 | -1 | DKK2,PPM1J,RARB,SOX7 |
| Role of CHK Proteins in Cell Cycle Checkpoint Control | 5.29 | 0.14 | -1.134 | BRCA1,CDC25C,CDK1,CLSPN,E2F7,E2F8,PLK1,PPM1J |
| ATM Signaling | 2.87 | 0.0722 | -1.134 | BRCA1,CCNB1,CCNB2,CDC25C,CDK1,PPM1J,RAD51 |
| Senescence Pathway | 1.71 | 0.0364 | -1.265 | CCNB1,CCNB2,CDC25C,CDK1,CDKN2B,E2F7,E2F8,ITPR3,PHF19,PPM1J |
| Cell Cycle: G2/M DNA Damage Checkpoint Regulation | 9.32 | 0.224 | -1.508 | AURKA,BRCA1,CCNB1,CCNB2,CDC25C,CDK1,CKS1B,PKMYT1,PLK1,TOP2A,WEE1 |
| Cell Cycle: G1/S Checkpoint Regulation | 3.02 | 0.0896 | -1.633 | CCND2,CCNE1,CCNE2,CDKN2B,E2F7,E2F8 |
| Dendritic Cell Maturation | 1.03 | 0.0328 | -1.633 | CD40,HLA-DQA1,HLA-DQB1,HLA-DRB5,Ighg2b,LTB |
| Endocannabinoid Cancer Inhibition Pathway | 1.01 | 0.035 | -2.236 | ADCY3,CCND2,CCNE1,CCNE2,TRIB3 |
