## Supplementary File 7 for "Dicer deficiency in microglia leads to accelerated demyelination and failed remyelination"

**Supplementary file 7:** List of differentially expressed gene (DEGs, og2FoldChange <-1/>+1, pAdj<0.05) from flow sorted microglia isolated from Cx3cr1[creERT2] Cre- control and Cre+ mutant animals undergoing remyelination after six weeks of CUP feeding.

| Gene.nam | log2FoldCl | pvalue | padj |
|---|---|---|---|
| Gdf3 | 2.574117 | 1.35E-05 | 0.005251 |
| Gm8696 | 1.691741 | 5.89E-13 | 1.08E-09 |
| Gm27008 | 1.627705 | 2.94E-05 | 0.008559 |
| Tie1 | 1.593762 | 3.89E-05 | 0.009952 |
| Lrrc32 | 1.54418 | 0.000257 | 0.034616 |
| Hspa1a | 1.468918 | 1.23E-44 | 7.89E-41 |
| Hspa1b | 1.43572 | 7.22E-66 | 9.25E-62 |
| Hspb1 | 1.38273 | 1.77E-07 | 0.000162 |
| Vstm4 | 1.37526 | 0.000167 | 0.026144 |
| Abcc9 | 1.360336 | 1.13E-05 | 0.004521 |
| Mylk | 1.351654 | 6.97E-05 | 0.01487 |
| Jcad | 1.333223 | 5.28E-06 | 0.002791 |
| Hsph1 | 1.299402 | 3.10E-24 | 1.32E-20 |
| Cxcl12 | 1.294322 | 1.98E-06 | 0.001204 |
| Dcn | 1.281291 | 0.000238 | 0.032488 |
| S1pr3 | 1.276032 | 1.63E-05 | 0.005811 |
| Esam | 1.246138 | 3.29E-05 | 0.008849 |
| Atp13a5 | 1.210191 | 0.000183 | 0.027273 |
| Vtn | 1.204381 | 1.23E-10 | 1.58E-07 |
| Col4a2 | 1.195646 | 0.000197 | 0.028728 |
| Rgs4 | 1.167055 | 0.00014 | 0.022962 |
| Rgs5 | 1.108342 | 0.000182 | 0.027273 |
| Igfbp7 | 1.100997 | 6.51E-06 | 0.003047 |
| Adgrf5 | 1.073351 | 5.22E-05 | 0.012607 |
| Slc6a20a | 1.054379 | 7.29E-05 | 0.015289 |
| Col14a1 | 1.025981 | 1.85E-05 | 0.006264 |
| Crtap | 1.014156 | 2.16E-12 | 3.45E-09 |

| Gene.nam | log2FoldCl | pvalue | padj |
|---|---|---|---|
| Itgal | -1.16352 | 4.67E-06 | 0.0026 |
| Gm24265 | -1.22622 | 2.72E-05 | 0.008142 |
| Cd24a | -1.54182 | 0.000126 | 0.022061 |
| Gm24407 | -1.58532 | 1.86E-05 | 0.006264 |
| Trem3 | -1.74957 | 8.03E-05 | 0.016589 |
| Cxcr2 | -2.05503 | 3.26E-05 | 0.008849 |
| Fpr1 | -2.08137 | 1.17E-07 | 0.000115 |
| Xlr4a | -2.20167 | 5.45E-06 | 0.002791 |
