## Supplementary File 8 for "Dicer deficiency in microglia leads to accelerated demyelination and failed remyelination"

**Supplementary file 8:** List of enriched biological processed of DEGs from flow sorted microglia isolated from Cx3cr1$^{creERT2}$ Cre- control and Cre+ mutant animals undergoing remyelination after six weeks of CUP feeding.

| Category | Term | Count | % | PValue | Genes | List Total | Pop Hits | Pop Total | Fold Enrich | Bonferroni | Benjamini | FDR |
|---|---|---|---|---|---|---|---|---|---|---|---|---|
| GOTERM_ | GO:0009408~response to heat | 4 | 12.5 | 9.20E-05 | 193740, 15 | 29 | 57 | 18082 | 43.7556 | 0.02597 | 0.026312 | 0.026312 |
| GOTERM_ | GO:0007155~cell adhesion | 6 | 18.75 | 8.02E-04 | 12818, 223 | 29 | 485 | 18082 | 7.713615 | 0.205066 | 0.114702 | 0.114702 |
| GOTERM_ | GO:0006986~response to unfolded protein | 3 | 9.375 | 0.002813 | 193740, 15 | 29 | 51 | 18082 | 36.67748 | 0.553221 | 0.268185 | 0.268185 |
| GOTERM_ | GO:0007166~cell surface receptor signaling pathway | 4 | 12.5 | 0.004592 | 12484, 224 | 29 | 219 | 18082 | 11.38844 | 0.731866 | 0.316177 | 0.316177 |
| GOTERM_ | GO:0016525~negative regulation of angiogenesis | 3 | 9.375 | 0.005528 | 12827, 131 | 29 | 72 | 18082 | 25.97989 | 0.795108 | 0.316177 | 0.316177 |
| GOTERM_ | GO:0006935~chemotaxis | 3 | 9.375 | 0.014289 | 14293, 203 | 29 | 118 | 18082 | 15.85213 | 0.983691 | 0.681095 | 0.681095 |
| GOTERM_ | GO:0022407~regulation of cell-cell adhesion | 2 | 6.25 | 0.019951 | 12484, 164 | 29 | 13 | 18082 | 95.92573 | 0.996861 | 0.695446 | 0.695446 |
| GOTERM_ | GO:0045744~negative regulation of G-protein coupled receptor protein signaling pathway | 2 | 6.25 | 0.019951 | 19737, 197 | 29 | 13 | 18082 | 95.92573 | 0.996861 | 0.695446 | 0.695446 |
| GOTERM_ | GO:0007204~positive regulation of cytosolic calcium ion concentration | 3 | 9.375 | 0.021885 | 14293, 124 | 29 | 148 | 18082 | 12.63886 | 0.998215 | 0.695446 | 0.695446 |
| GOTERM_ | GO:0022409~positive regulation of cell-cell adhesion | 2 | 6.25 | 0.030534 | 12484, 164 | 29 | 20 | 18082 | 62.35172 | 0.999859 | 0.837726 | 0.837726 |
| GOTERM_ | GO:0007159~leukocyte cell-cell adhesion | 2 | 6.25 | 0.036533 | 12484, 164 | 29 | 24 | 18082 | 51.95977 | 0.999976 | 0.837726 | 0.837726 |
| GOTERM_ | GO:0050850~positive regulation of calcium-mediated signaling | 2 | 6.25 | 0.036533 | 12484, 164 | 29 | 24 | 18082 | 51.95977 | 0.999976 | 0.837726 | 0.837726 |
| GOTERM_ | GO:0035987~endodermal cell differentiation | 2 | 6.25 | 0.041008 | 22370, 128 | 29 | 27 | 18082 | 46.18646 | 0.999994 | 0.837726 | 0.837726 |
| GOTERM_ | GO:2001234~negative regulation of apoptotic signaling pathway | 2 | 6.25 | 0.041008 | 15507, 155 | 29 | 27 | 18082 | 46.18646 | 0.999994 | 0.837726 | 0.837726 |
| GOTERM_ | GO:0006955~immune response | 3 | 9.375 | 0.065969 | 22370, 124 | 29 | 272 | 18082 | 6.877028 | 1 | 1 | 1 |
| GOTERM_ | GO:0007157~heterophilic cell-cell adhesion via plasma membrane cell adhesion molecules | 2 | 6.25 | 0.077528 | 12484, 164 | 29 | 52 | 18082 | 23.98143 | 1 | 1 | 1 |
| GOTERM_ | GO:0009968~negative regulation of signal transduction | 2 | 6.25 | 0.078961 | 19737, 197 | 29 | 53 | 18082 | 23.52895 | 1 | 1 | 1 |
| GOTERM_ | GO:0032526~response to retinoic acid | 2 | 6.25 | 0.08467 | 21846, 298 | 29 | 57 | 18082 | 21.8778 | 1 | 1 | 1 |
| GOTERM_ | GO:0043408~regulation of MAPK cascade | 2 | 6.25 | 0.087511 | 14562, 124 | 29 | 59 | 18082 | 21.13618 | 1 | 1 | 1 |
| GOTERM_ | GO:0007200~phospholipase C-activating G-protein coupled receptor signaling pathway | 2 | 6.25 | 0.088929 | 14293, 127 | 29 | 60 | 18082 | 20.78391 | 1 | 1 | 1 |
| GOTERM_ | GO:0009612~response to mechanical stimulus | 2 | 6.25 | 0.093169 | 13179, 203 | 29 | 63 | 18082 | 19.7942 | 1 | 1 | 1 |
| GOTERM_ | GO:0006954~inflammatory response | 3 | 9.375 | 0.098728 | 14293, 136 | 29 | 344 | 18082 | 5.43765 | 1 | 1 | 1 |
