## Supplementary File 9 for "Dicer deficiency in microglia leads to accelerated demyelination and failed remyelination"

**Supplementary file 9:** List of differentially expressed gene (DEGs, log2FoldChange <-1/>+1, pAdj<0.05) from flow sorted microglia isolated from $Cx3cr1^{creERT2}$ Cre- control and Cre+ mutant animals treated with TMX at peak of demyelination and undergoing remyelination after six weeks of CUP feeding.

| Gene.nam | log2FoldCl | pvalue | padj |
|---|---|---|---|
| Lgi2 | 5.646702 | 0.000178 | 0.003324 |
| Cenpm | 5.433124 | 3.75E-06 | 0.000129 |
| Pimreg | 5.011539 | 6.04E-28 | 4.36E-25 |
| Elane | 4.955209 | 0.003725 | 0.036971 |
| Kif18b | 4.879842 | 6.08E-26 | 3.20E-23 |
| Parpbp | 4.834011 | 2.60E-08 | 1.56E-06 |
| E2f7 | 4.442156 | 8.13E-13 | 1.08E-10 |
| Ankle1 | 4.400154 | 1.88E-28 | 1.71E-25 |
| Stil | 4.348262 | 6.17E-21 | 1.73E-18 |
| Mki67 | 4.347122 | 1.64E-92 | 2.25E-88 |
| Pbk | 4.318284 | 1.77E-39 | 4.85E-36 |
| Nek2 | 4.292235 | 1.69E-22 | 5.14E-20 |
| Rrm2 | 4.272233 | 7.22E-35 | 8.99E-32 |
| Fam83d | 4.231904 | 2.94E-08 | 1.74E-06 |
| Depdc1a | 4.181683 | 9.09E-09 | 5.96E-07 |
| Rad51 | 4.176633 | 2.23E-16 | 4.30E-14 |
| Sapcd2 | 4.159095 | 6.80E-08 | 3.70E-06 |
| Troap | 4.075276 | 2.77E-14 | 4.41E-12 |
| Gm8091 | 4.057479 | 0.000101 | 0.002103 |
| Ccnb2 | 4.051208 | 2.85E-27 | 1.70E-24 |
| Cdc25c | 4.03217 | 5.70E-08 | 3.13E-06 |
| 4930438A( | 3.978154 | 0.000111 | 0.00227 |
| Spc24 | 3.947154 | 3.19E-14 | 4.96E-12 |
| Birc5 | 3.93299 | 7.28E-24 | 2.49E-21 |
| Kif4 | 3.913908 | 3.51E-14 | 5.35E-12 |
| Kif23 | 3.896366 | 5.68E-31 | 5.55E-28 |
| Adgrg3 | 3.894503 | 0.004477 | 0.04195 |
| Ube2c | 3.873427 | 2.09E-24 | 7.75E-22 |
| Chil4 | 3.859863 | 3.10E-05 | 0.000793 |
| Depdc1b | 3.856729 | 1.37E-09 | 1.09E-07 |
| Neil3 | 3.831967 | 1.32E-16 | 2.63E-14 |
| Kif11 | 3.755412 | 1.21E-39 | 4.14E-36 |
| E2f8 | 3.736407 | 6.22E-14 | 9.26E-12 |
| Top2a | 3.734245 | 1.93E-65 | 1.32E-61 |
| Aspm | 3.690344 | 3.54E-25 | 1.57E-22 |
| Cdca2 | 3.663938 | 1.59E-24 | 6.41E-22 |
| Esco2 | 3.655959 | 1.14E-14 | 1.89E-12 |
| Ccnb1 | 3.643653 | 2.97E-39 | 6.79E-36 |
| Cep55 | 3.592782 | 5.58E-12 | 6.48E-10 |

| Gene.nam | log2FoldCl | pvalue | padj |
|---|---|---|---|
| P3h3 | -1.00065 | 2.82E-08 | 1.68E-06 |
| Gjb6 | -1.00066 | 0.000572 | 0.008589 |
| Aatk | -1.00372 | 2.21E-08 | 1.35E-06 |
| Ccdc122 | -1.00477 | 0.005005 | 0.0454 |
| Mmd2 | -1.00663 | 8.20E-05 | 0.001788 |
| Scg3 | -1.00797 | 9.91E-05 | 0.002087 |
| Nceh1 | -1.00957 | 8.93E-11 | 8.67E-09 |
| Cib2 | -1.01625 | 0.000293 | 0.004979 |
| Gm44321 | -1.01633 | 0.004148 | 0.039898 |
| Tmem221 | -1.01775 | 2.19E-05 | 0.000589 |
| Mfsd13a | -1.01797 | 4.50E-06 | 0.000149 |
| Acsl6 | -1.02168 | 1.17E-05 | 0.000345 |
| Angptl7 | -1.02387 | 8.71E-07 | 3.67E-05 |
| Fam171b | -1.02454 | 0.003575 | 0.035848 |
| 4833411C( | -1.02494 | 1.51E-06 | 5.91E-05 |
| Aldh1l1 | -1.02578 | 0.000205 | 0.003736 |
| Gpm6b | -1.02602 | 1.91E-09 | 1.45E-07 |
| Ly6a | -1.027 | 1.11E-05 | 0.000329 |
| Slamf8 | -1.0324 | 4.44E-07 | 2.00E-05 |
| Cacna1a | -1.03527 | 0.000251 | 0.004374 |
| Id4 | -1.03571 | 0.002718 | 0.029051 |
| Nckap1 | -1.03594 | 0.000179 | 0.003325 |
| 4-Sep | -1.0391 | 0.001368 | 0.017154 |
| Trappc6a | -1.03926 | 1.72E-06 | 6.65E-05 |
| Mta1 | -1.04428 | 0.000536 | 0.008152 |
| Gm10602 | -1.04746 | 0.001779 | 0.021168 |
| Maoa | -1.04762 | 0.002059 | 0.023643 |
| Cd83 | -1.04938 | 7.46E-14 | 1.10E-11 |
| Myom1 | -1.05661 | 0.000144 | 0.002804 |
| Htra1 | -1.05784 | 6.26E-07 | 2.74E-05 |
| Fam26f | -1.06222 | 0.000422 | 0.006718 |
| Fmnl2 | -1.06354 | 0.000409 | 0.00656 |
| Plp1 | -1.07302 | 8.09E-07 | 3.45E-05 |
| Pirb | -1.08057 | 1.28E-13 | 1.83E-11 |
| Nat8f4 | -1.08441 | 0.00458 | 0.042621 |
| Zeb2os | -1.0874 | 1.76E-07 | 8.76E-06 |
| Mex3d | -1.08936 | 0.001941 | 0.022539 |
| H2-K2 | -1.09486 | 1.90E-06 | 7.20E-05 |
| Tnfrsf19 | -1.09645 | 0.001044 | 0.013787 |

| | | | |
|---|---|---|---|
| Cdca5 | 3.592533 | 1.83E-11 | 1.97E-09 |
| Hmmr | 3.562198 | 8.05E-26 | 4.08E-23 |
| Ckap2 | 3.557374 | 1.51E-15 | 2.68E-13 |
| Sgo1 | 3.509399 | 5.78E-09 | 3.92E-07 |
| Sgol2a | 3.490094 | 1.03E-09 | 8.42E-08 |
| Prc1 | 3.469542 | 1.67E-38 | 3.26E-35 |
| Nusap1 | 3.461704 | 1.06E-22 | 3.31E-20 |
| Acap1 | 3.46085 | 0.000178 | 0.003324 |
| Nuf2 | 3.459903 | 5.75E-18 | 1.29E-15 |
| Tpx2 | 3.440887 | 1.50E-27 | 9.36E-25 |
| Shcbp1 | 3.426246 | 5.86E-16 | 1.08E-13 |
| Psrc1 | 3.414363 | 5.16E-07 | 2.29E-05 |
| Knstrn | 3.410726 | 2.94E-17 | 6.20E-15 |
| Aurkb | 3.406154 | 1.03E-25 | 5.04E-23 |
| Hp | 3.396252 | 0.005223 | 0.046796 |
| Kif22 | 3.357928 | 1.07E-31 | 1.13E-28 |
| Cdc20 | 3.335035 | 3.47E-33 | 3.96E-30 |
| Kif20a | 3.325234 | 1.27E-27 | 8.31E-25 |
| Gck | 3.310732 | 5.41E-14 | 8.14E-12 |
| Brip1 | 3.305837 | 1.01E-10 | 9.78E-09 |
| Pif1 | 3.253031 | 1.14E-05 | 0.000337 |
| Lockd | 3.249407 | 1.04E-08 | 6.77E-07 |
| Cdca3 | 3.24232 | 3.85E-20 | 1.06E-17 |
| Cdkn3 | 3.213108 | 3.21E-08 | 1.87E-06 |
| Cdca8 | 3.207515 | 1.95E-19 | 5.03E-17 |
| Ticrr | 3.2046 | 6.51E-09 | 4.39E-07 |
| Gtse1 | 3.202653 | 1.74E-11 | 1.89E-09 |
| Cenpe | 3.193921 | 1.42E-23 | 4.62E-21 |
| Ncapg | 3.189553 | 4.30E-21 | 1.23E-18 |
| Spag5 | 3.179315 | 4.35E-19 | 1.08E-16 |
| Knl1 | 3.171155 | 2.61E-22 | 7.77E-20 |
| Plk1 | 3.161222 | 7.19E-23 | 2.29E-20 |
| Bub1b | 3.159303 | 5.44E-35 | 7.45E-32 |
| Racgap1 | 3.154197 | 2.39E-44 | 1.09E-40 |
| Mxd3 | 3.13394 | 3.18E-08 | 1.87E-06 |
| Ndc80 | 3.130228 | 1.16E-17 | 2.52E-15 |
| Melk | 3.117711 | 1.29E-12 | 1.65E-10 |
| Xkr5 | 3.111801 | 2.90E-05 | 0.000746 |
| Bub1 | 3.094936 | 2.01E-13 | 2.81E-11 |
| Dlgap5 | 3.08243 | 3.06E-13 | 4.23E-11 |
| Ccna2 | 3.081903 | 9.16E-38 | 1.39E-34 |
| Aurka | 3.075249 | 7.29E-19 | 1.78E-16 |
| Ccnf | 3.0426 | 3.07E-28 | 2.48E-25 |
| Trim59 | 3.029515 | 5.33E-17 | 1.11E-14 |
| Uhrf1 | 3.026647 | 1.65E-24 | 6.47E-22 |
| Cdc6 | 3.004737 | 1.75E-09 | 1.34E-07 |
| Foxm1 | 2.965679 | 3.51E-24 | 1.27E-21 |

| | | | |
|---|---|---|---|
| Gm16576 | -1.0978 | 1.06E-06 | 4.32E-05 |
| Gm27676 | -1.09794 | 0.000691 | 0.009969 |
| Acsf2 | -1.09852 | 0.000127 | 0.002536 |
| Gstm5 | -1.09883 | 7.33E-07 | 3.15E-05 |
| Cd63 | -1.10094 | 5.73E-25 | 2.45E-22 |
| Cx3cr1 | -1.10125 | 1.92E-25 | 8.75E-23 |
| Bcar3 | -1.10488 | 0.001906 | 0.022311 |
| Tnfsf12 | -1.10682 | 0.000934 | 0.012632 |
| Tmem47 | -1.10847 | 3.38E-05 | 0.000854 |
| A930007I1 | -1.1115 | 3.18E-05 | 0.000807 |
| Slc6a11 | -1.11504 | 2.50E-09 | 1.87E-07 |
| Timp4 | -1.11609 | 0.000133 | 0.002616 |
| Sh3bp4 | -1.1168 | 0.004486 | 0.041998 |
| C920006O | -1.13116 | 0.003741 | 0.036998 |
| Cbs | -1.13214 | 0.005451 | 0.048207 |
| Dmtn | -1.13796 | 0.000204 | 0.003713 |
| Sdc4 | -1.13859 | 3.30E-09 | 2.35E-07 |
| Dhtkd1 | -1.13877 | 0.002174 | 0.024656 |
| Gfap | -1.14562 | 0.000618 | 0.009118 |
| AI464131 | -1.15359 | 0.000182 | 0.003378 |
| D5Ertd605 | -1.15912 | 0.000947 | 0.012788 |
| Gm37310 | -1.16027 | 0.00464 | 0.043029 |
| Wscd1 | -1.16169 | 2.73E-08 | 1.63E-06 |
| Slc14a1 | -1.1659 | 5.09E-05 | 0.001198 |
| Clybl | -1.16867 | 0.000792 | 0.011061 |
| Nbl1 | -1.17011 | 5.51E-05 | 0.00128 |
| Zfp783 | -1.17371 | 1.54E-07 | 7.78E-06 |
| Rorb | -1.17492 | 0.000826 | 0.01147 |
| Serpina3n | -1.17617 | 0.00506 | 0.045747 |
| Sat2 | -1.17649 | 6.82E-05 | 0.001534 |
| Tril | -1.17677 | 0.000238 | 0.004206 |
| Mtss1l | -1.17871 | 0.001457 | 0.018051 |
| Gpc5 | -1.18288 | 0.000118 | 0.002383 |
| Gstt2 | -1.18341 | 6.75E-05 | 0.001523 |
| Gpm6a | -1.18552 | 0.005698 | 0.049843 |
| Dgke | -1.1899 | 4.41E-06 | 0.000147 |
| Galnt3 | -1.19389 | 4.22E-05 | 0.001019 |
| Scara3 | -1.19811 | 0.001125 | 0.014701 |
| Lrguk | -1.19971 | 0.004677 | 0.043211 |
| Ddah1 | -1.20059 | 0.000516 | 0.00796 |
| Gadd45b | -1.20059 | 0.000734 | 0.010441 |
| Gm43890 | -1.20383 | 0.000891 | 0.012159 |
| 1700011I0 | -1.21244 | 0.005196 | 0.046642 |
| Mfsd7a | -1.21424 | 0.002608 | 0.02815 |
| Rab11fip4 | -1.21895 | 0.000285 | 0.004864 |
| Mbd2 | -1.22916 | 1.77E-07 | 8.78E-06 |
| S1pr1 | -1.23354 | 5.11E-22 | 1.49E-19 |

| | | | | | | | |
|---|---|---|---|---|---|---|---|
| Spc25 | 2.941627 | 1.16E-16 | 2.34E-14 | Gstt3 | -1.23909 | 0.001334 | 0.016814 |
| Ttk | 2.902917 | 2.21E-11 | 2.33E-09 | Gm37306 | -1.24074 | 7.24E-08 | 3.92E-06 |
| Anln | 2.851683 | 1.15E-23 | 3.83E-21 | Atp6v0e2 | -1.24818 | 0.002364 | 0.026345 |
| Dtl | 2.81798 | 1.17E-18 | 2.76E-16 | Slc39a4 | -1.25591 | 1.98E-08 | 1.22E-06 |
| Clspn | 2.798633 | 2.95E-07 | 1.39E-05 | Gm42970 | -1.26664 | 0.001591 | 0.019402 |
| Gm22267 | 2.789835 | 1.09E-05 | 0.000325 | Tmem205 | -1.268 | 6.14E-06 | 0.000195 |
| Ccne1 | 2.763897 | 1.19E-05 | 0.000349 | Gabrg1 | -1.27023 | 0.000312 | 0.005246 |
| Cdk1 | 2.73304 | 1.98E-24 | 7.53E-22 | Sod3 | -1.27321 | 0.00388 | 0.037909 |
| Rad54b | 2.685192 | 1.10E-06 | 4.45E-05 | Ptprf | -1.27526 | 0.003779 | 0.037236 |
| Mcm10 | 2.680779 | 3.03E-06 | 0.000108 | AW11201( | -1.27668 | 1.41E-09 | 1.11E-07 |
| Eme1 | 2.667882 | 1.88E-05 | 0.000519 | Dnaaf3 | -1.28718 | 4.10E-06 | 0.000139 |
| Foxr1 | 2.650595 | 1.39E-05 | 0.000401 | Nr1d1 | -1.29528 | 7.60E-05 | 0.001682 |
| Ska3 | 2.642357 | 9.03E-08 | 4.76E-06 | Gm10524 | -1.30443 | 8.67E-05 | 0.001869 |
| Skint3 | 2.641178 | 1.31E-12 | 1.66E-10 | Klf15 | -1.30518 | 0.004217 | 0.040335 |
| Kif2c | 2.61671 | 4.89E-13 | 6.57E-11 | Cyp2j9 | -1.32661 | 0.00037 | 0.006043 |
| Mis18bp1 | 2.597554 | 2.36E-12 | 2.78E-10 | C1rl | -1.32864 | 3.45E-09 | 2.44E-07 |
| C330027C( | 2.575741 | 2.59E-11 | 2.67E-09 | Ighj1 | -1.3317 | 7.90E-08 | 4.20E-06 |
| Prkcq | 2.565509 | 3.09E-27 | 1.76E-24 | Arhgap23 | -1.33501 | 9.22E-05 | 0.001965 |
| Ect2 | 2.54309 | 1.27E-18 | 2.94E-16 | Gm44789 | -1.33955 | 0.000291 | 0.004964 |
| Arhgef39 | 2.468443 | 4.30E-19 | 1.08E-16 | Rxrg | -1.34366 | 5.55E-05 | 0.001288 |
| Wee1 | 2.448773 | 8.89E-11 | 8.67E-09 | Ltbp3 | -1.34419 | 0.000675 | 0.009782 |
| Prr11 | 2.422249 | 9.88E-07 | 4.09E-05 | Soat2 | -1.35181 | 0.003457 | 0.034894 |
| Klrd1 | 2.421844 | 3.51E-13 | 4.76E-11 | Apcdd1 | -1.36071 | 0.002149 | 0.024413 |
| Gm5431 | 2.395796 | 1.64E-11 | 1.79E-09 | Sh2d6 | -1.36308 | 0.000478 | 0.007451 |
| Oip5 | 2.39403 | 3.75E-05 | 0.000923 | Adgre4 | -1.3744 | 1.18E-05 | 0.000345 |
| Pole | 2.379359 | 7.86E-12 | 8.98E-10 | Rgs4 | -1.37589 | 0.005095 | 0.045913 |
| Pi16 | 2.37224 | 0.0049 | 0.044719 | Gm26244 | -1.37941 | 0.000698 | 0.010046 |
| Ddias | 2.360941 | 1.93E-05 | 0.000532 | Wwc1 | -1.39585 | 0.002368 | 0.026345 |
| Fanci | 2.341165 | 4.31E-06 | 0.000144 | Clec1a | -1.39969 | 0.0006 | 0.008908 |
| 4930479D: | 2.337571 | 3.04E-06 | 0.000108 | Zkscan7 | -1.40002 | 0.001951 | 0.022611 |
| Cdkn1a | 2.332736 | 2.85E-38 | 4.87E-35 | Kank1 | -1.40274 | 0.001053 | 0.013878 |
| Brca1 | 2.321423 | 2.48E-10 | 2.28E-08 | Gm24924 | -1.40799 | 1.21E-06 | 4.86E-05 |
| Pclaf | 2.314915 | 3.00E-09 | 2.17E-07 | Ackr3 | -1.42277 | 0.000917 | 0.012434 |
| Il1r2 | 2.299289 | 2.67E-05 | 0.000696 | Gm5547 | -1.42421 | 0.000771 | 0.010805 |
| Pkmyt1 | 2.292898 | 1.16E-15 | 2.10E-13 | Gm128 | -1.42603 | 0.005073 | 0.045809 |
| Alb | 2.289958 | 7.39E-09 | 4.87E-07 | Fblim1 | -1.45373 | 1.85E-14 | 3.02E-12 |
| Padi4 | 2.284146 | 0.000536 | 0.008152 | Tnni3 | -1.46756 | 5.71E-05 | 0.001323 |
| Rad51ap1 | 2.28009 | 3.04E-11 | 3.11E-09 | Epb41l3 | -1.473 | 7.53E-11 | 7.47E-09 |
| Cit | 2.275928 | 9.16E-14 | 1.32E-11 | Pdgfrb | -1.50964 | 7.23E-05 | 0.001612 |
| Syt6 | 2.259814 | 6.40E-06 | 0.000202 | Ighj4 | -1.51673 | 0.000178 | 0.003324 |
| Cdc45 | 2.257093 | 4.61E-07 | 2.07E-05 | Rapsn | -1.53861 | 3.69E-10 | 3.26E-08 |
| Tcf19 | 2.23529 | 4.23E-11 | 4.26E-09 | Lag3 | -1.54058 | 2.96E-16 | 5.62E-14 |
| Plac8 | 2.234339 | 0.000184 | 0.003409 | Abcc9 | -1.54103 | 0.004076 | 0.039428 |
| Cenpf | 2.234033 | 7.84E-17 | 1.60E-14 | 2900026A( | -1.5705 | 3.52E-11 | 3.57E-09 |
| Ercc6l | 2.22343 | 2.80E-09 | 2.05E-07 | Gm31410 | -1.5782 | 1.27E-05 | 0.00037 |
| 5330426L2 | 2.204534 | 4.68E-06 | 0.000153 | Atp9a | -1.58299 | 4.32E-05 | 0.001038 |
| Cenpi | 2.204457 | 9.02E-08 | 4.76E-06 | Gm13710 | -1.61245 | 1.73E-06 | 6.65E-05 |

| | | | |
|---|---|---|---|
| Gm42047 | 2.201563 | 1.08E-15 | 1.97E-13 |
| Spn | 2.180259 | 0.003076 | 0.031733 |
| Cxcr2 | 2.176424 | 0.004942 | 0.044947 |
| Plp2 | 2.169639 | 0.000311 | 0.005235 |
| Gm38832 | 2.158259 | 2.49E-07 | 1.20E-05 |
| Zwilch | 2.157495 | 8.39E-07 | 3.56E-05 |
| Hyal5 | 2.134157 | 5.47E-09 | 3.73E-07 |
| Chaf1a | 2.132162 | 3.87E-08 | 2.22E-06 |
| Kif18a | 2.122658 | 1.97E-06 | 7.44E-05 |
| Kifc1 | 2.118906 | 0.001207 | 0.015569 |
| Ncapd2 | 2.100669 | 8.05E-25 | 3.34E-22 |
| Fignl1 | 2.086788 | 6.14E-15 | 1.06E-12 |
| Tmc5 | 2.069746 | 0.00121 | 0.015594 |
| Kif15 | 2.069383 | 1.10E-10 | 1.04E-08 |
| Mrap2 | 2.048776 | 0.002377 | 0.026383 |
| Gata2 | 2.035262 | 2.44E-05 | 0.000645 |
| Stmn1 | 2.029817 | 2.57E-28 | 2.20E-25 |
| Coro2a | 2.013203 | 3.04E-09 | 2.19E-07 |
| Ube2t | 2.011224 | 4.84E-05 | 0.001149 |
| Cenpa | 2.01036 | 2.71E-17 | 5.79E-15 |
| Perp | 2.004173 | 3.66E-10 | 3.25E-08 |
| Ckap2l | 2.004051 | 3.20E-13 | 4.39E-11 |
| Hells | 1.98685 | 3.13E-07 | 1.47E-05 |
| Tnfrsf26 | 1.984024 | 0.00065 | 0.009517 |
| Exo1 | 1.97241 | 0.000122 | 0.002451 |
| Cenpk | 1.970712 | 0.003054 | 0.031526 |
| Diaph3 | 1.967536 | 4.95E-05 | 0.001171 |
| Phf19 | 1.966867 | 0.000207 | 0.003755 |
| Cenpn | 1.96444 | 6.47E-06 | 0.000204 |
| Asf1b | 1.963681 | 2.36E-14 | 3.80E-12 |
| Nid2 | 1.956319 | 8.59E-28 | 5.89E-25 |
| Smc4 | 1.942486 | 5.78E-20 | 1.55E-17 |
| Gen1 | 1.934091 | 0.000164 | 0.003123 |
| B230311B( | 1.931472 | 0.000739 | 0.01049 |
| Mcm5 | 1.926144 | 1.52E-25 | 7.18E-23 |
| Mastl | 1.918273 | 9.58E-06 | 0.00029 |
| Ccne2 | 1.916617 | 3.35E-10 | 3.00E-08 |
| Pilrb2 | 1.91053 | 0.003778 | 0.037236 |
| Ptger3 | 1.906196 | 4.38E-16 | 8.23E-14 |
| Hcar2 | 1.869511 | 2.81E-06 | 0.000102 |
| Kif20b | 1.862241 | 1.71E-08 | 1.07E-06 |
| Ezh2 | 1.834173 | 9.67E-15 | 1.62E-12 |
| 1700020N( | 1.827529 | 1.68E-05 | 0.000476 |
| Ly75 | 1.825013 | 0.001365 | 0.017141 |
| Chtf18 | 1.823389 | 1.30E-06 | 5.18E-05 |
| Tacc3 | 1.799024 | 9.05E-19 | 2.18E-16 |
| Ptpn7 | 1.776189 | 0.000422 | 0.006718 |

| | | | |
|---|---|---|---|
| Phactr3 | -1.61963 | 0.005316 | 0.047283 |
| Etnppl | -1.62615 | 4.50E-06 | 0.000149 |
| Il12rb2 | -1.63696 | 0.000333 | 0.005529 |
| Slc16a9 | -1.63846 | 0.000144 | 0.002803 |
| Gm17025 | -1.64012 | 0.003501 | 0.035267 |
| Bdh1 | -1.6479 | 0.003193 | 0.032593 |
| Anxa9 | -1.64822 | 0.004069 | 0.039391 |
| Gm13391 | -1.69345 | 0.000994 | 0.013303 |
| Ptgfrn | -1.69989 | 1.79E-05 | 0.0005 |
| Gm22068 | -1.70896 | 0.004702 | 0.043344 |
| Dusp2 | -1.71147 | 1.16E-05 | 0.00034 |
| Xlr4a | -1.71404 | 0.000494 | 0.007668 |
| Gabbr2 | -1.7264 | 0.000237 | 0.004197 |
| Mt2 | -1.74302 | 0.000153 | 0.002942 |
| Xlr3b | -1.76062 | 1.11E-10 | 1.04E-08 |
| Gm25360 | -1.76719 | 0.00376 | 0.037131 |
| Lox | -1.77435 | 7.22E-07 | 3.11E-05 |
| Gpr4 | -1.79166 | 0.002697 | 0.028887 |
| Megf10 | -1.80149 | 0.000408 | 0.00656 |
| Rhobtb3 | -1.82293 | 0.000865 | 0.01191 |
| Ighj2 | -1.83846 | 1.26E-07 | 6.53E-06 |
| Gm38091 | -1.88055 | 0.000619 | 0.009121 |
| Car11 | -1.92688 | 0.000531 | 0.008112 |
| Gm2115 | -1.951 | 0.001942 | 0.022539 |
| Rap1gap | -1.98406 | 4.16E-05 | 0.001008 |
| Xlr3a | -1.99464 | 0.002368 | 0.026345 |
| Atp13a5 | -2.03497 | 0.001193 | 0.015427 |
| Gpr162 | -2.09175 | 1.91E-08 | 1.18E-06 |
| Car4 | -2.12422 | 0.000111 | 0.00227 |
| Cd22 | -2.18589 | 3.91E-24 | 1.37E-21 |
| Clic5 | -2.19005 | 0.001221 | 0.015703 |
| Aass | -2.26569 | 0.002936 | 0.030616 |
| AF529169 | -2.28131 | 0.000168 | 0.003174 |
| D7Ertd128 | -2.50074 | 1.56E-05 | 0.000445 |

| | | | |
|---|---|---|---|
| Pask | 1.763097 | 6.90E-07 | 2.98E-05 |
| Unc5cl | 1.761707 | 8.96E-05 | 0.001922 |
| E2f1 | 1.758944 | 1.66E-12 | 2.01E-10 |
| Upk1b | 1.727895 | 3.28E-27 | 1.80E-24 |
| Rbm44 | 1.722587 | 0.003553 | 0.035678 |
| Slc43a3 | 1.716344 | 4.40E-08 | 2.49E-06 |
| Gm27008 | 1.698537 | 1.12E-05 | 0.000331 |
| Adamtsl2 | 1.694725 | 3.90E-10 | 3.42E-08 |
| Hlf | 1.688394 | 7.87E-14 | 1.15E-11 |
| H2afx | 1.677043 | 1.71E-12 | 2.05E-10 |
| Chad | 1.674773 | 5.09E-05 | 0.001198 |
| Cdkn2c | 1.67277 | 5.09E-10 | 4.36E-08 |
| Itgb3 | 1.671323 | 1.57E-08 | 9.94E-07 |
| Il18rap | 1.666621 | 5.13E-05 | 0.001206 |
| Timeless | 1.658889 | 2.92E-06 | 0.000105 |
| Ccl2 | 1.657506 | 3.40E-14 | 5.24E-12 |
| Tedc1 | 1.655759 | 1.80E-08 | 1.11E-06 |
| 4930579G2 | 1.653877 | 0.000114 | 0.002316 |
| Hist1h1c | 1.639617 | 2.59E-12 | 3.03E-10 |
| Phlda3 | 1.633704 | 0.000777 | 0.010881 |
| D7Ertd443 | 1.632593 | 9.18E-05 | 0.001961 |
| Dnph1 | 1.620811 | 0.00493 | 0.044914 |
| Fbln1 | 1.607061 | 2.18E-05 | 0.000588 |
| Glipr2 | 1.582112 | 0.004028 | 0.039107 |
| Gpsm2 | 1.560407 | 0.002259 | 0.025436 |
| Tbc1d5 | 1.538898 | 8.28E-06 | 0.000255 |
| Hyal6 | 1.517634 | 0.000198 | 0.003629 |
| Atad2 | 1.514703 | 4.68E-18 | 1.07E-15 |
| Gm7931 | 1.506563 | 0.003123 | 0.032001 |
| Fam212b | 1.504686 | 0.002543 | 0.027716 |
| Cdc7 | 1.50331 | 1.17E-06 | 4.68E-05 |
| Prim2 | 1.489246 | 2.35E-11 | 2.43E-09 |
| Siglec5 | 1.475284 | 1.57E-16 | 3.08E-14 |
| Gm15232 | 1.444909 | 5.96E-08 | 3.26E-06 |
| Traip | 1.434052 | 0.002894 | 0.030238 |
| Mcm2 | 1.416896 | 1.98E-11 | 2.12E-09 |
| Kntc1 | 1.408665 | 1.39E-08 | 8.95E-07 |
| Fmr1nb | 1.408287 | 0.001586 | 0.019396 |
| Lmnb1 | 1.406201 | 1.46E-09 | 1.14E-07 |
| Abca13 | 1.402809 | 1.54E-05 | 0.000439 |
| Cks1b | 1.400535 | 7.13E-06 | 0.000222 |
| Cldn12 | 1.398329 | 1.49E-08 | 9.52E-07 |
| Hmgb2 | 1.394006 | 1.15E-12 | 1.50E-10 |
| Cks2 | 1.385238 | 4.97E-08 | 2.79E-06 |
| Slc5a10 | 1.382409 | 2.15E-06 | 7.99E-05 |
| Cdc25b | 1.375388 | 3.04E-06 | 0.000108 |
| Trip13 | 1.374091 | 0.001647 | 0.019924 |

| | | | |
|---|---|---|---|
| Mcc | 1.370812 | 6.93E-05 | 0.00155 |
| Mmp28 | 1.360045 | 2.86E-05 | 0.000738 |
| Chaf1b | 1.343588 | 0.004123 | 0.039726 |
| Acvr2a | 1.341573 | 7.55E-15 | 1.29E-12 |
| Wdhd1 | 1.331253 | 2.15E-07 | 1.05E-05 |
| Prim1 | 1.327824 | 8.46E-07 | 3.58E-05 |
| Smc2 | 1.326923 | 1.12E-09 | 9.06E-08 |
| Hist1h4i | 1.323464 | 0.001261 | 0.016077 |
| Gm19708 | 1.31597 | 0.002839 | 0.030033 |
| Spdl1 | 1.301192 | 0.00555 | 0.048918 |
| Ncapg2 | 1.300976 | 4.35E-10 | 3.77E-08 |
| C5ar2 | 1.293234 | 9.65E-15 | 1.62E-12 |
| Gm43305 | 1.292498 | 1.83E-05 | 0.000508 |
| Lig1 | 1.277041 | 2.36E-07 | 1.15E-05 |
| Mycbp | 1.273672 | 2.52E-10 | 2.30E-08 |
| Tk1 | 1.266409 | 6.80E-09 | 4.56E-07 |
| Dock9 | 1.257572 | 1.34E-11 | 1.51E-09 |
| Gpr165 | 1.255062 | 3.61E-05 | 0.000891 |
| Dusp8 | 1.242327 | 0.001797 | 0.021358 |
| Ppm1e | 1.232293 | 2.48E-06 | 9.05E-05 |
| Mcm6 | 1.226385 | 1.42E-12 | 1.77E-10 |
| Ska2 | 1.226242 | 4.64E-06 | 0.000152 |
| Incenp | 1.211662 | 1.55E-10 | 1.44E-08 |
| Map3k19 | 1.208175 | 4.21E-05 | 0.001019 |
| Il7r | 1.207552 | 3.07E-14 | 4.83E-12 |
| Rgs14 | 1.202404 | 1.06E-10 | 1.01E-08 |
| Hirip3 | 1.201201 | 6.04E-05 | 0.001384 |
| Kpna2 | 1.192281 | 4.33E-15 | 7.61E-13 |
| Myl2 | 1.182477 | 2.67E-05 | 0.000696 |
| Frat2 | 1.17953 | 0.005673 | 0.049766 |
| Acvr2b | 1.176889 | 3.36E-08 | 1.94E-06 |
| E2f2 | 1.17684 | 3.59E-05 | 0.000885 |
| Tmc7 | 1.173547 | 9.57E-12 | 1.08E-09 |
| Postn | 1.16772 | 7.08E-09 | 4.73E-07 |
| Hist1h2be | 1.16199 | 1.65E-08 | 1.04E-06 |
| Rxfp1 | 1.15959 | 3.52E-05 | 0.000877 |
| Myo18b | 1.158652 | 1.08E-08 | 7.03E-07 |
| Hsf2bp | 1.137554 | 0.00439 | 0.041383 |
| Tmem2 | 1.136277 | 8.85E-07 | 3.72E-05 |
| Pop1 | 1.130945 | 0.001115 | 0.014597 |
| Adrb2 | 1.128877 | 3.54E-28 | 2.69E-25 |
| Mms22l | 1.128705 | 1.68E-07 | 8.43E-06 |
| Smtn | 1.127534 | 0.000191 | 0.003518 |
| Ccl7 | 1.11777 | 0.003308 | 0.033565 |
| Lmtk2 | 1.116547 | 0.001619 | 0.01967 |
| Scarb1 | 1.11089 | 1.26E-09 | 1.01E-07 |
| Hspa1a | 1.104994 | 6.49E-18 | 1.43E-15 |

| | | | |
|---|---|---|---|
| Fanca | 1.103584 | 0.000271 | 0.004669 |
| Dennd3 | 1.102804 | 0.005442 | 0.048187 |
| Mcm7 | 1.100544 | 3.12E-09 | 2.24E-07 |
| Ube2g2 | 1.097812 | 1.37E-11 | 1.53E-09 |
| Heatr1 | 1.096007 | 6.81E-12 | 7.84E-10 |
| Igsf8 | 1.09388 | 0.001736 | 0.020748 |
| Cdkn2d | 1.068845 | 3.87E-05 | 0.000949 |
| Rrm1 | 1.048544 | 1.26E-12 | 1.62E-10 |
| Sipa1l1 | 1.042104 | 2.24E-11 | 2.34E-09 |
| Hist1h2bg | 1.039514 | 0.001943 | 0.022539 |
| Hspa1b | 1.030915 | 1.84E-13 | 2.60E-11 |
| Entpd6 | 1.02762 | 2.72E-09 | 2.01E-07 |
| Wdr62 | 1.02636 | 6.88E-06 | 0.000215 |
| Adora1 | 1.022467 | 0.00132 | 0.016677 |
| F9 | 1.013769 | 2.74E-06 | 9.94E-05 |
| Kdm1b | 1.011064 | 3.62E-09 | 2.54E-07 |
| Cdc25a | 1.007327 | 5.59E-06 | 0.00018 |
| Igfbp4 | 1.004393 | 1.38E-12 | 1.73E-10 |
| Rad54l | 1.002424 | 0.000603 | 0.008928 |
